# Exploring the possible role of hybridization in the evolution of photosynthetic pathways in *Flaveria* (Asteraceae), the prime model of C_4_ photosynthesis evolution

**DOI:** 10.1101/2022.01.31.478436

**Authors:** Diego F. Morales-Briones, Gudrun Kadereit

## Abstract

*Flaveria* (Asteraceae) is the prime model for the study of C_4_ photosynthesis evolution and seems to support a stepwise acquisition of the pathway through C_3_-C_4_ intermediate phenotypes, still existing in *Flaveria* today. Molecular phylogenies of *Flaveria* based on concatenated data matrices are currently used to reconstruct the complex sequence of trait shifts during C_4_ evolution. To assess the possible role of hybridization in C_4_ evolution in *Flaveria*, we re-analyzed transcriptome data of 17 *Flaveria* species to infer the extent of gene tree discordance and possible reticulation events. We found massive gene tree discordance as well as reticulation along the backbone and within clades containing C_3_-C_4_ intermediate and C_4_-like species. An early hybridization event between two C_3_ species might have triggered C_4_ evolution in the genus. The clade containing all C_4_ species plus the C_4_-like species *F. vaginata* and *F. palmeri* is robust but of hybrid origin involving *F. angustifolia* and *F. sonorensis* (both C_3_-C_4_ intermediate) as parental lineages. Hybridization seems to be a driver of C_4_ evolution in *Flaveria* and likely promoted the fast acquisition of C_4_ traits. This new insight can be used in further exploring C_4_ evolution and can inform C_4_ bioengineering efforts.

## INTRODUCTION

The detection of gene tree discordance is common in the phylogenomic era. Discordance can be the product of multiple processes and is commonly attributed to either incomplete lineage sorting (ILS) and/or hybridization (Pamilo and Nei, 1988; Doyle, 1992; Galtier and Daubin, 2008). Hybridization is a fundamental process in the evolution of animals, plants, and fungi (Giraud et al., 2008; Schwenk et al., 2008; Soltis and Soltis, 2009; Payseur and Rieseberg, 2016), and methods to investigate hybridization in a phylogenetic context recently have been developed greatly. These include methods that estimate phylogenetic networks while accounting for ILS and hybridization simultaneously (e.g., Solís-Lemus and Ané, 2016; Wen et al., 2018), and methods that detect hybridization based on site patterns or phylogenetic invariants (e.g., (Green et al., 2010; Durand et al., 2011; Kubatko and Chifman, 2019). The current ease to produce phylogenomic data sets and the availability of new analytical methods facilitate the exploration of reticulate evolution in any clade across the Tree of Life, including those that have particular significance as model lineages, such as the flowering plant genus *Flaveria* (Asteraceae) for the study of C_4_ photosynthesis evolution.

*Flaveria* belongs to the sunflower tribe Heliantheae (Anderberg et al., 2007). According to the most recent revision by Powell (1978), *Flaveria* includes 21 morphologically rather similar species distributed mainly in southern USA and northern Mexico, with few species occurring in the Caribbean and South America. The two weedy and self-compatible C_4_ species, *F. trinervia* and *F. bidentis*, have been introduced almost worldwide (https://powo.science.kew.org/). Species of *Flaveria* usually show scattered occurrences in unconnected, localized populations near rivers, creeks, irrigation canals, fields, roadsides, and ponds, often on saline or gypseous soils (Powell, 1978). They are either robust shrubs or herbaceous perennials, or annuals (mainly the C_4_ species). The genus stands out in Asteraceae for its reduced floral features and reduced and secondarily aggregated capitula (Anderberg et al., 2007). Reduction is most evident in *F. trinervia* (C_4_) and aggregation of capitula mimicking a single capitulum in *F. anomala* (C_3_-C_4_). *Flaveria* is consistently diploid (see Powell, 1978 and ref. therein; only exceptions are some tetraploid populations of *F. pringlei*) with a haploid chromosome number of *n* = 18. Artificial hybridization among 16 species of *Flaveria* was successful, and F1 hybrids could be obtained in most species’ combinations (see Table 1 in Powell, 1978). F2 and backcross crosses also resulted in offspring in a high number of combinations, but only four of these were fertile. Powell (1978) excluded frequent natural hybridization in *Flaveria* mainly because of geographical isolation.

**Table 1.** Photosynthetic types in *Flaveria* according to Sage et al. (2014 and 2018 and ref. therein); species names marked * were not sampled in this study, mesophyll (M), bundle sheath (BS), glycine decarboxylase (GDC).

| Photosynthetic type | Characteristics | *Flaveria* species representing this type<sup>1</sup> |
| --- | --- | --- |
| C<sub>3</sub> | Photorespiratory cycle operates completely within single M cells, BS cells small<sup>2</sup> with few organelles, veins widely spaced | *F. cronquistii* <sup>2</sup>,<br>*F. mcdougallii* * |
| C<sub>3</sub> proto-kranz | Functionally C<sub>3</sub>, activated BS cells, greater vein density, mitochondria localized at the inner BS wall adjacent to the vasculature | *F. pringlei*,<br>*F. robusta* |
| C<sub>2</sub> Type I | Low or no GDC expression in M cells, high number of centripetally located organelles in the BS cells, CO<sub>2</sub> compensation point reduced in comparison to C<sub>3</sub>, lack of any C<sub>4</sub> cycle | *F. angustifolia*,<br>*F. chlorifolia*,<br>*F. sonorensis* |
| C<sub>2</sub> Type II | In addition to type I, modest C<sub>4</sub> cycle enhancement | *F. anomala*,<br>*F. floridana*,<br>*F. linearis* *,<br>*F. pubescens*,<br>*F. ramosissima* |
| C<sub>4</sub>-like | Strong C<sub>4</sub> metabolic cycle but also weak C<sub>3</sub> cycle in the M cells | *F. brownii*,<br>*F. palmeri*,<br>*F. vaginata* |
| C<sub>4</sub> | No C<sub>3</sub> cycle, CO<sub>2</sub>-saturate photosynthesis below 500 ppm CO<sub>2</sub> | *F. bidentis*,<br>*F. campestre* *,<br>*F. kochiana*,<br>*F. trinervia*,<br>*F. australasica*, |

<sup>1</sup> *Flaveria oppositifolia*\* so far only classified as C<sub>2</sub> without further specification.
<sup>2</sup> *Flaveria cronquistii* qualifies as a C<sub>3</sub>+ species (see Sage et al., 2018) because it has larger and photosynthetically more active BS cells (McKown and Dengler, 2007).

C_4_ photosynthesis in *Flaveria* was first recognized by Smith and Turner (1975), and the presence of C_3_-C_4_ intermediate species was first noted by Brown (pers. comm. in Powell, 1978), first verified by Apel and Maass (1981) and then studied in detail biochemically in four species by Ku et al. (1983) and Nakamoto et al. (1983). Numerous publications characterizing the physiology and biochemistry of C_3_-C_4_ intermediate species of *Flaveria* followed (Ku et al., 1991 and ref. therein). At the same time crossing experiments of C_3_ and C_4_ *Flaveria* species as well as backcrosses or crosses between C_3_-C_4_ intermediate species revealed the transfer of C_4_ properties as well as the simultaneous functioning of C_3_ and C_4_ pathways in hybrids (see Apel et al., 1988 as an example and Kadereit et al., 2017 for review). Of all genera that contain C_3_-C_4_ intermediate species, *Flaveria* has the highest diversity of C_3_-C_4_ phenotypes including C_2_ photosynthesis (Sage et al., 2012; briefly described in Table 1), and arguably is the only lineage that allows to infer a detailed sequence of increasing C_4_-ness (Sage et al., 2012). Against this background, *Flaveria* qualified as the model group for the establishment (Monson and Moore, 1989) and subsequent refinement of a model of stepwise acquisition of C_4_ photosynthesis through intermediate steps (Sage et al., 2014 and ref. therein).

One challenge of the *Flaveria* model is that C_3_-C_4_ intermediate phenotypes might also have resulted from reticulation during the diversification of the genus, especially when reproductive barriers are leaky among extant species as soon as they get into contact (Powell, 1978) and hybridization between C_3_ and C_4_ species is possible. Therefore, a phylogenetic study exploring the occurrence and location of past reticulation events is needed. Comprehensive molecular phylogenetic studies of *Flaveria* published so far were either based on few molecular markers only (McKown et al., 2005), or on concatenated data matrices and inference methods unable to reveal tree discordance, possible reticulation, or incomplete lineage sorting (Lyu et al., 2015). Irrespective of this, numerous current studies of evolutionary change during the establishment of the C_4_ pathway rely on the *Flaveria* model (e.g., Lyu et al., 2021; Taniguchi et al., 2021).

The aim of this study is to use available transcriptome data of 17 species of *Flaveria* to assess the extent of reticulation during the diversification of the genus, and to evaluate these findings with respect to the evolution of C_4_ photosynthesis in the genus and its suitability as general model of C_4_ evolution.

## MATERIAL AND METHODS

### Taxon sampling

We included publicly available transcriptomes from 17 species of *Flaveria* (Table S1). In addition, we included outgroups from four genomes of Asteraceae (*Chrysanthemum, Helianthus, Lactuca* and *Stevia*) following (Mandel et al., 2019; Table S1).

### Homology and orthology inference

Raw read processing, transcriptome assembly, low-quality and chimeric transcript removal, transcript clustering into putative genes, translation, and final coding sequences (CDS) redundancy assessment were carried out following Morales-Briones et al. (2021) with minor modifications as follows. Sequencing errors in raw reads were corrected with Rcorrector (Song and Florea, 2015) and reads flagged as uncorrectable were discarded. Sequencing adapters and low-quality bases were removed with Trimmomatic v 0.39 (Bolger et al., 2014). Additionally, chloroplast and mitochondrial reads were filtered out with Bowtie2 v 2.4.4 (Langmead and Salzberg, 2012) using publicly available Asterales organelle genomes from the Organelle Genome Resources database (RefSeq; [Pruitt et al., 2007]; last accessed on June 4, 2021) as references. Read quality was assessed with FastQC v 0.11.9 (http://www.bioinformatics.bbsrc.ac.uk/projects/fastqc) and overrepresented sequences were discarded. *De novo* assembly was carried out with Trinity v 2.13.2 (Grabherr et al., 2011) with default settings, but without in silico normalization. Assembly quality was assessed with Transrate v 1.0.3 (Smith-Unna et al., 2016). Low quality and poorly supported transcripts were removed using individual cut-off values for three contig score components of Transrate: 1) proportion of nucleotides in a contig that agrees in identity with the aligned read, s(Cnuc) ≤ 0.25; 2) proportion of nucleotides in a contig that have one or more mapped reads, s(Ccov) ≤0.25; and 3) proportion of reads that map to the contig in correct orientation, s(Cord) ≤ 0.5. Furthermore, chimeric transcripts (*trans*-self and *trans*-multi-gene) were removed following the approach described in Yang and Smith (2013) using *Helianthus annuus* as the reference proteome, and a percentage similarity and length cutoff of 30 and 100, respectively. To remove isoforms and assembly artifacts, filtered reads were remapped to filtered transcripts with Salmon v 1.5.2 (Patro et al., 2017) and putative genes were clustered with Corset v 1.09 (Davidson and Oshlack, 2014) using default settings, except that we used a minimum of five reads as threshold to remove transcripts with low coverage (-m 5). Only the longest transcript of each putative gene inferred by Corset was retained as suggested in Chen et al. (2019). Filtered transcripts were translated with TransDecoder v 5.3.0 (Haas et al., 2013) with default settings and the proteomes of *Arabidopsis thaliana, Helianthus annuus*, and *Lactuca sativa* to identify open reading frames. Finally, coding sequences (CDS) from translated amino acids were further reduced with CD-HIT v 4.8.1 (-c 0.99; [Fu et al., 2012]) to remove near-identical sequences. Scripts used can be found at https://bitbucket.org/yanglab/phylogenomic_dataset_construction/src/master/ (Morales-Briones et al., 2021).

Homology inference was done with an all-by-all BLASTN search on CDS with an *E* value cutoff of 10. BLAST hits were filtered with a minimal hit coverage of 40%. Homolog groups were clustered with MCL v 14-137 (van Dongen, 2000) using a minimum minus log-transformed *E* value cutoff of 5 and an inflation value of 1.4, and only clusters with at least 17 taxa were retained. Homolog cluster sequences were aligned using the OMM_MACSE v 11.05 pipeline (Scornavacca et al., 2019). Alignments were further trimmed to remove columns with more than 90% missing data using Phyx (Brown et al., 2017). Homolog trees were inferred using RAxML v 8.2.11 (Stamatakis, 2014) with the GTRCAT model and 200 rapid bootstrap (BS) replicates. Monophyletic and paraphyletic tips of the same species were removed, keeping the tip with the highest number of characters in the trimmed alignment following Yang and Smith (2014). Spurious tips were detected and removed using TreeShrink v 1.3.9 (Mai and Mirarab, 2018) with the ‘per-gene’ mode, a false positive error rate threshold (α) of 0.05, and excluding the outgroups. Trees were visually inspected, and deep paralogs producing internal branch lengths longer than 0.20 were cut apart retaining subclades with at least 10 taxa to obtain final homolog trees. Orthology inference was done using the ‘monophyletic outgroup’ (MO) approach from Yang and Smith (2014). The MO approach filters for trees that have outgroup taxa being monophyletic and single-copy, and therefore filters for single- and low-copy genes. This approach roots the gene tree by the outgroups, traverses the rooted tree from root to tip, and removes the side with less taxa when gene duplication is detected (Yang and Smith, 2014). If no taxon duplication is detected in a homolog tree, the MO approach outputs a one-to-one ortholog. We set all species of *Flaveria* as ingroups, and *Chrysanthemum, Helianthus, Lactuca*, and *Stevia* as outgroups, keeping orthologs that included at least 10 taxa.

### Tree inference and detection of gene tree conflict

Sequences from individual orthologs were aligned using the OMM_MACSE pipeline. Columns with more than 20% missing data were trimmed with Phyx, and only alignments with at least 500 characters and all 21 taxa were retained and concatenated. We estimated a maximum likelihood (ML) tree of the concatenated matrix with IQ-TREE v 2.1.3 (Minh et al., 2020) searching for the best partition scheme (Lanfear et al., 2012) followed by ML gene tree inference and 1000 ultrafast bootstrap replicates for clade support. To estimate a coalescent-based species tree, first, we inferred individual gene trees with IQ-TREE using extended model selection (Kalyaanamoorthy et al., 2017) and 200 non-parametric bootstrap replicates for clade support. Gene trees were then used to infer a species tree with ASTRAL-III v 5.7.7 (Zhang et al., 2018) using local posterior probabilities (LPP; Sayyari and Mirarab, 2016) to assess clade support.

We explored gene tree discordance by calculating the number of concordant and discordant bipartitions on each node of the concatenated and ASTRAL trees using Phyparts (Smith et al., 2015). We used individual gene trees with BS support of at least 50% for each node. Additionally, to distinguish conflict from poorly supported branches, we carried out a Quartet Sampling (QS; Pease et al., 2018) analysis using the concatenated matrix with a partition by gene (--genetrees), the concatenated IQ-TREE and ASTRAL trees, and 1000 replicates. To further visualize gene tree conflict, we built a cloudogram with the DensiTree function from Phangorn v 2.7.1 (Schliep, 2011) in R (R Core Team 2021). We first time-calibrated individual ortholog gene trees, for visualization purposes only, with TreePL v 1.0 (Smith and O’Meara, 2012). The most recent common ancestor (MRCA) of *Helianthus* and *Flaveria* was fixed to 21.5 MYA, and the MRCA of *Flaveria* was fixed to 4.3 MYA based on Mandel et al. (2019).

### Testing for potential reticulation

First, to investigate if gene tree discordance can be explained by ILS alone, we performed coalescent simulations like Cloutier et al. (2019). An ultrametric tree with branch lengths in mutational units (μT) was inferred with PAUP v 4.0a (build 168; Swofford, 2002) by constraining a ML tree search to the ASTRAL tree and using the concatenated alignment, a GTRGAMMA model, and enforcing a strict molecular clock. The mutational branch lengths from the constrained tree and branch lengths in coalescent units (τ = T/4N_e_) from the ASTRAL tree were used to estimate the population size parameter theta (θ = μT/τ; Degnan and Rosenberg, 2009) for internal branches. Terminal branches were set with θ = 1. We then used Phybase v 1.4 (Liu and Yu, 2010), that implements the formula from Rannala and Yang (2003), to simulate 10,000 gene trees using the constraint tree and the estimated theta values. Lastly, we calculated the distribution of Robinson and Foulds (1981) tree-to-tree distances between the ASTRAL tree and each original gene tree using Phangorn and compared this with the distribution of tree-to-tree distances between the ASTRAL tree and the simulated gene trees.

To test for potential reticulation, we inferred species networks using maximum pseudo-likelihood (Yu and Nakhleh, 2015) in PhyloNet v 3.8.2 (Than et al., 2008) with the command “InferNetworks_MPL’’ and using individual ML gene trees as input. We included all 17 species of *Flaveria* and *Helianthus* as an outgroup (18-taxon data set). Network searches were performed allowing for up to ten reticulation events and using only nodes in the gene trees that have BS ≥ 50%. To find the network with optimal number of reticulations, we plotted the number of reticulations versus the pseudo-likelihood score to determine when the score stabilizes (Blair and Ané, 2020). Given that the pseudo-likelihood scores did not stabilize after networks with ten reticulation events (see results) and to reduce network complexity, we performed one additional round of searches by removing taxa involved in reticulation from the previous search. We removed *F. brownii* (C_4_-like) and *F. pringlei* (C_3_; 16-taxon data set). These two species were inferred to be product of reticulation events in all ten original searches and were not involved in additional reticulation events (i.e., they are not a parental lineage of other reticulation events; see results). Network searches for the reduced data set were carried out similarly as with the original data set.

Additionally, we tested for hybridization with HyDe (Blischak et al., 2018), which uses site pattern frequencies (Kubatko and Chifman, 2019) to quantify the hybridization parameter γ between two parental lineages that form a hybrid lineage. We tested all triplet combinations using the ‘run_hyde.py’ script, the concatenated alignment, and a mapping file to assign species. Test significance was assessed with a Bonferroni correction (α= 0.05) for the number of hypothesis tests conducted with estimates of γ between 0 and 1 (Blischak et al., 2018). We carried out HyDe hybridization test using all species and without *F. pringlei* (C_3_), as it has been identified as a potential artificial hybrid (Lyu et al., 2015).

### Assessment of whole genome duplication

To investigate potential whole genome duplication, product of reticulation events in *Flaveria* (see results), we mapped gene duplication events onto the inferred species tree following (Yang et al., 2018). First, we extracted rooted ingroup clades (orthogroup) from the final homolog trees by requiring at least 15 taxa and only orthogroups with an average BS ≥ 50 were used for mapping. Then gene duplication events were then mapped onto the MRCA on the species tree when two or more taxa overlapped between the two daughter clades on the rooted ingroup clade. Each node on a species tree can be counted only once from each gene tree to avoid nested gene duplications inflating the number of recorded duplications (Yang et al., 2018). Orthogroup extraction and mapping were carried out using the scripts “extract_clades.py” and “map_dups_mrca.py” from https://bitbucket.org/blackrim/clustering (Yang et al., 2018).

## RESULTS

### Orthology inference and phylogenetic analysis

The final number of orthologs with at least ten species was 5,374 with a mean of 4,809 orthologs per species (Table S1). The number of orthologs that included all 21 species was 1,295. The concatenated matrix (≥ 500 bp per ortholog) consisted of 1,582,170 aligned columns with a character occupancy of 93% from 1,249 orthologs.

The topologies from the IQ-TREE and ASTRAL trees were similar and most nodes had maximum support (BS =100, LPP = 100; Fig. 1; Fig. S1). The two trees only differed in the placement of *Flaveria bidentis* (C_4_) which had lower support on both cases (Fig. 1; Fig. S1). *Flaveria cronquistii* (C_3_), *F. robusta* (C_3_) and *F. pringle*i (C_3_) + *F. angustifolia* (C_3_-C_4_) formed a grade which is sister to Clade A + Clade B (clades names followed McKown et al., 2005). Clade A from IQ-TREE comprised the same species and relationships as in Lyu et al. (2015). *Flaveria ramosissima* (C_3_-C_4_) and *F. palmeri* (C_4_-like) were successive sisters to *F. kochiana* (C_4_) + *F. vaginata* (C_4_-like) and the C_4_ clade [(*F. bidentis, F. trinervia* + *F australasica*); BS = 83]. The ASTRAL tree recovered *F. bidentis* (C_4_) sister to *F. kochiana* (C_4_) + *F. vaginata* (C_4_-like; LLP = 0.96; Fig. S1). Clade B included the same species as in Lyu et al. (2015) but also included *F. sonorensis* (C_3_-C_4_). Our analyses recovered *F. brownii* (C_4_-like) + *F. sonorensis* (C_3_-C_4_) as sister to the remaining species in Clade B. The remaining species, all C_3_-C_4_, had similar relationships as in Lyu et al. (2015) with *F. anomala* as sister to the clade *F. pubescens, F. chlorifolia* + *F. floridana*.

**Fig. 1.**
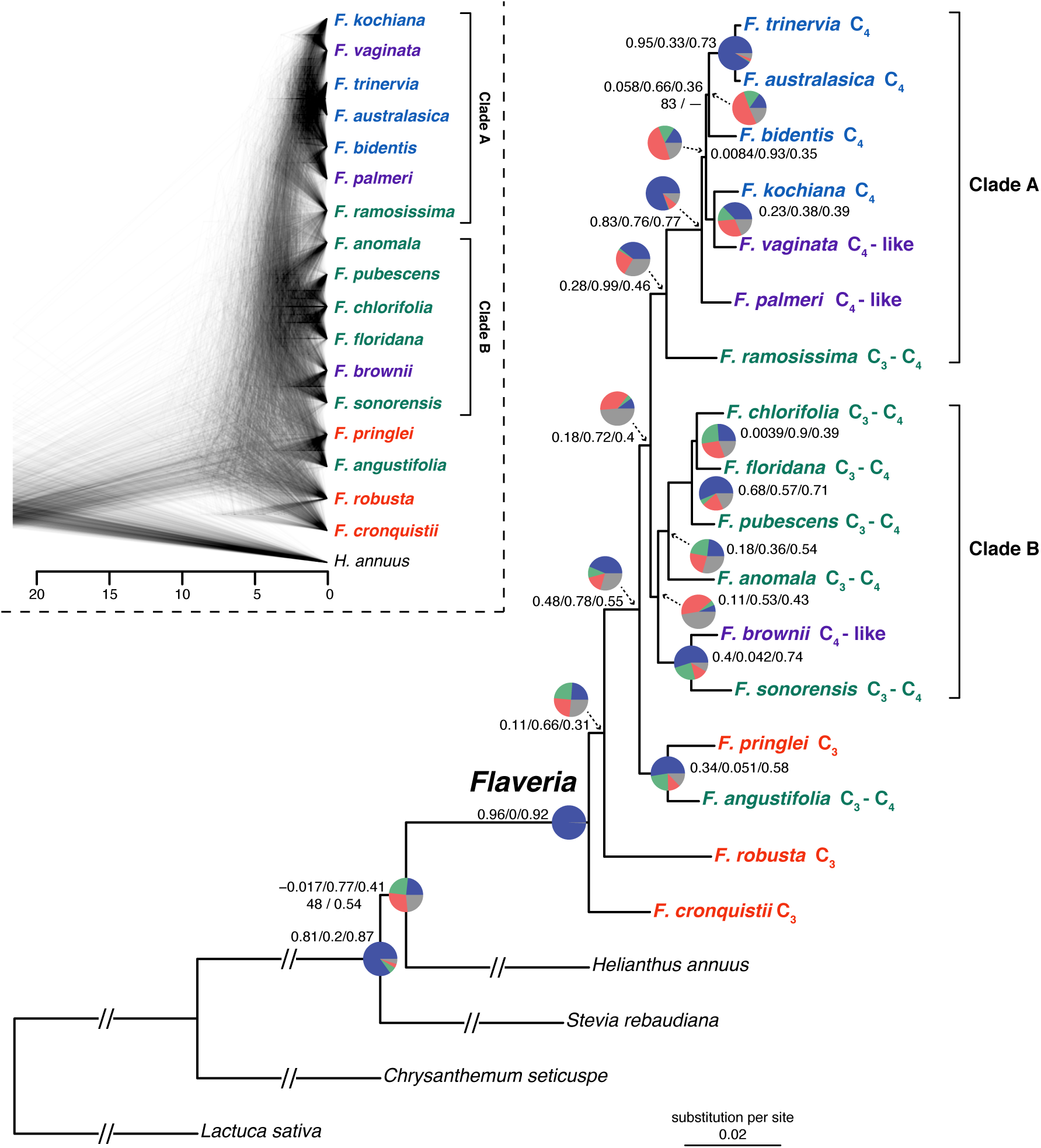
Maximum likelihood phylogeny of *Flaveria* inferred with IQ-TREE from the concatenated 1249-nuclear gene supermatrix. Species names are colored by photosynthetic type. Quartet Sampling (QS) scores are shown above branches. QS scores: Quartet concordance/Quartet differential/Quartet informativeness. All nodes have full bootstrap support (BS =100) and local posterior probability (LLP =1) unless noted next to branches. Em dashes (—) denotes an alternative topology compared to the ASTRAL tree (Fig. S1). Pie charts represent the proportion of ortholog trees that support a clade (blue), the main alternative bifurcation (green), the remaining alternatives (red), and bifurcations (conflict or support) with <50% bootstrap support (gray). Branch lengths as number of substitutions per site (scale bar). Exceptionally long branches were shortened with a broken segment (//) for illustration purposes (See Fig. S1 for original branch lengths). Inset: Cloudogram inferred from 1288 nuclear ortholog trees. Scale in millions of years ago (MA).

### Phylogenetic conflict

Overall, conflict analyses and cloudogram visualization revealed rampant gene tree discordance in *Flaveria* (Fig. 1; Fig S2). The cloudogram showed significant conflict along the backbone of the phylogeny as well as within clades A and B (Fig. 1). The placement of *F. robusta* (C_3_) was supported only by 308 (of 947) informative gene trees and had low QS support (0.11/0.66/0.3) with a clear signal of an alternative topology involving *F. cronquistii*. The sister relationship of *F. pringlei* (C_3_) and *F. angustifolia* (C_3_-C_4_) was supported by 677 (of 1,123) gene trees and had moderate QS support (0.34/0.051/0.58) and a strong signal of an alternative topology. The placement of *F. pringlei* (C_3_) + *F. angustifolia* (C_3_-C_4_) as sister of Clade A + Clade B was supported by 562 (of 909) gene trees and had strong QS support (0.48/0.78/0.55) with moderate signal for an alternative topology. The clade composed of clades A and B was supported only by 131 (of 661) gene trees and had low QS support (0.18/0.72/0.4) but had no signals of an alternative topology. Clade A was supported by 505 (of 858) gene trees and had moderate QS support (0.28/0.99/0.46) but showed no signals of alternative topologies. The remaining species of Clade A [excluding *F. ramosissima* (C_3_-C_4_)] formed a clade supported by most gene trees (1,037 of 1,151) and a strong QS score (0.83/0.76/0.77) with no signals of an alternative topology. *Flaveria kochiana* (C_4_) + *F. vaginata* (C_4_-like) was supported by 479 (of 1,049) gene trees and had moderate QS support (0.23/0.38/0.39) with a clear signal of an alternative topology. *Flaveria trinervia* (C_4_) + *F. australasica* (C_4_) was supported by most gene trees (1,172 of 1,220) and had strong QS support with no signals of alternative topologies. *Flaveria bidentis* (C_4_) as sister to *F. trinervia* (C_4_) + *F. australasica* (C_4_; IQ-TREE) was supported by 197 (of 1,054) and had very low QS support with a signal of an alternative topology. When *F. bidentis* (C_4_) was placed as sister to *F. kochiana* (C_4_) + *F. vaginata* (C_4_-like; ASTRAL) the clade formed by these three species was supported by even fewer gene trees (170 of 1,037) and QS counter-support (−0.027/0.72/0.38; Fig. S3). The clade composed of *F. kochiana* (C_4_) + *F. vaginata* (C_4_-like) and the C_4_ clade *F. bidentis, F. trinervia* + *F australasica* was supported by 206 (of 1,028) gene trees and had very low QS support (0.0084/0.93/0.35) with signals of alternative topologies. Clade B was supported by only 85 (of 679) gene trees and had low QS support (0.11/0.53/0.43) with signals of an alternative topology. *Flaveria brownii* (C_4_-like) + *F. sonorensi*s (C_3_-C_4_) was supported by 708 (of 1,180) gene trees and had moderate QS support (0.4/0.042/0.74) with clear signals of an alternative topology. The four remaining species of Clade B formed a clade supported by 299 (of 910) gene trees and had low QS support (0.11/0.53/0.43) with signals of an alternative topology. *Flaveria chlorifolia* (C_3_-C_4_) + *F. floridana* (C_3_-C_4_) was supported by 337 (of 1,042) gene trees and had very low QS support (0.0039/0.9/0.39) with clear signals of an alternative topology. Lastly, the C_3_-C_4_ clade (*F. pubescens, F. chlorifolia* + *F. floridana*) was supported by 727 (of 1,055) gene trees and had strong QS support (0.68/0.57/0.71) with a slight signal of an alternative topology.

### Potential widespread reticulation

The distribution of tree-to-tree distances of the empirical and simulated gene trees to the ASTRAL tree showed some overlap (Fig. S4), but there was a skew towards larger distances in the empirical trees (mean 16.61) compared to distances in the simulated trees (mean 11.80). This suggested that ILS alone cannot explain most of the observed gene tree incongruence (Maureira-Butler et al., 2008).

Species network analyses recovered topologies with up to ten reticulations events for the 18-taxon data set (Fig. S6). The pseudo-likelihood score (Fig. S5) continually improved with the inclusion of additional reticulation events, and the network with ten reticulations (Fig. 2A) had the best score. Although the best-scored network showed complicated and nested reticulation patterns (Fig. 2A), there are several clear patterns among most networks (Fig. S6). *Flaveria brownii* (C_4_-like) and *F. pringlei* (C_3_) were consistently recovered as hybrids in all ten networks (Fig. S6). In both cases, parental lineages and inheritance probabilities were consistent across networks. Other reticulation patterns recovered across networks were the hybrid origin of clades A (mainly C_4_ and C_4_-like) and B (mainly C_3_-C_4_), which both had *F. sonorensis* (C_3_-C_4_; itself a hybrid in some networks) and *F. angustifolia* (C_3_-C_4_; or closely related to lineage) as potential parental lineages. The last reticulation event recovered in all networks involved the C_3_ species *F. cronquistii* and *F. robusta* which were potential parental lineages of all remaining *Flaveria* species. The analyses of the 16-taxon data set (Fig. S6) resulted in a best-scoring network with five reticulation events (Fig. 2B). This showed patterns like the 18-taxon dataset regarding the hybrid origin of clades A and B, and the deep reticulation involving *F. cronquistii* (C_3_) and *F. robusta* (C_3_). Also, it recovered a reticulation event (consistent with the 18-taxon dataset) where *F. vaginata* (C_4_-like) and *F. kochiana* (C_4_) within Clade A had a hybrid origin.

**Fig. 2.**
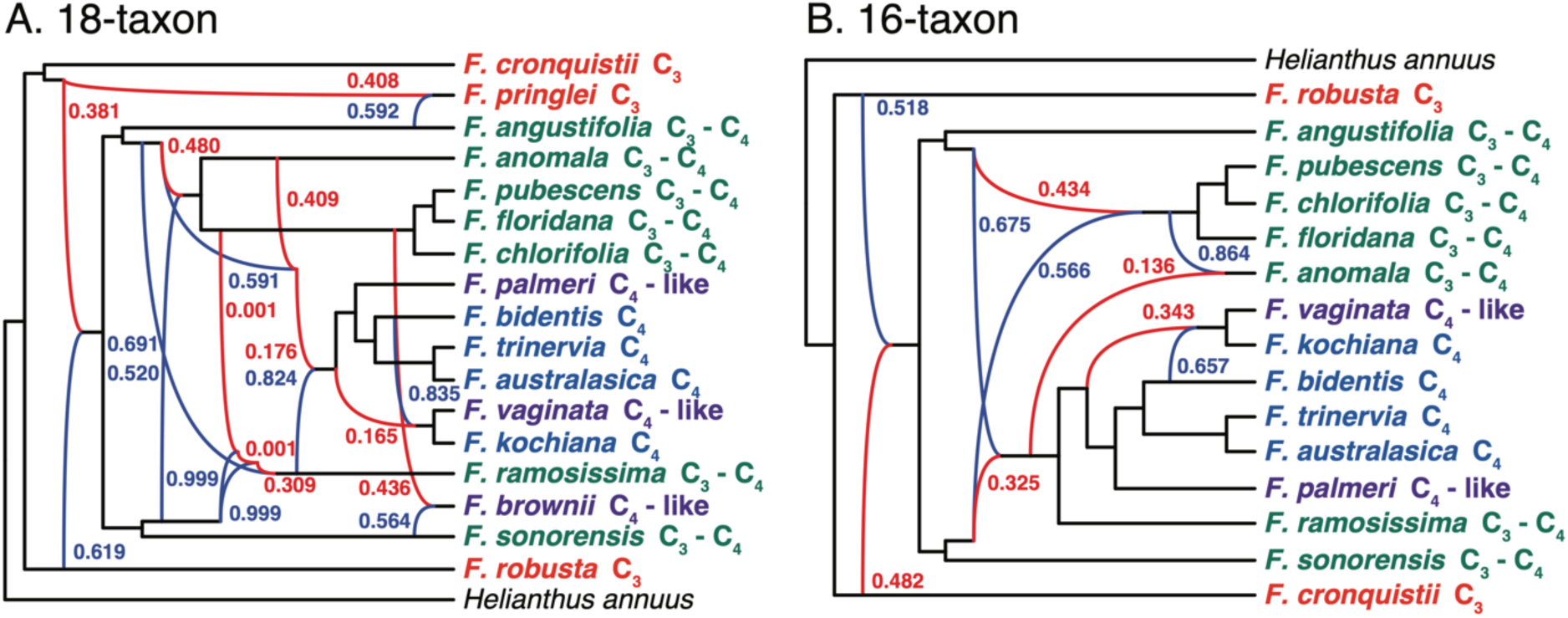
*Flaveria* species networks with the best maximum pseudo-likelihood scores with PhyloNet for the (A) 18-taxon and (B) 16-taxon data sets. Species names are colored by photosynthetic type. Red and blue curved branches indicate the minor and major edges, respectively, of hybrid nodes. Numbers next to curved branches indicate inheritance probabilities for each hybrid node.

The HyDe analysis of all possible triples using the 17 species of *Flaveria* resulted in 2,040 hybridization tests, of which 311 triples were significant (Table S2). The analyses without *F. pringlei* (C_3_) resulted in 247 significant hybridization tests of 1,680 overall tests (Table S3). HyDe analyses detected 12 species as potential hybrids (Fig. 3). These species included all members of Clade B which comprises five C_3_-C_4_ and one C_4_-like species. Hybrids detected in Clade B had several potential parental lineages from across *Flaveria* and admixture (γ) values were either closer to zero or one (Fig. 3). *Flaveria chlorifolia, F. floridana, F. pubescens, F. anomala* showed similar hybridization patterns consistent with a single ancient hybrid origin of these species as shown in the PhyloNet analyses. On the other hand, *F. sonorensis* and *F. brownii* showed hybridization patterns different from the rest of the species of the clade, suggesting their independent hybrid origins as also seen in the PhyloNet analyses. In Clade A, and unlike PhyloNet, HyDe detected only four instances of hybridization. These included the C_3_-C_4_ species *F. ramosissima* and the two C_4_-like species, *F. vaginata* and *F. palmeri*, of the clade.

**Fig. 3.**
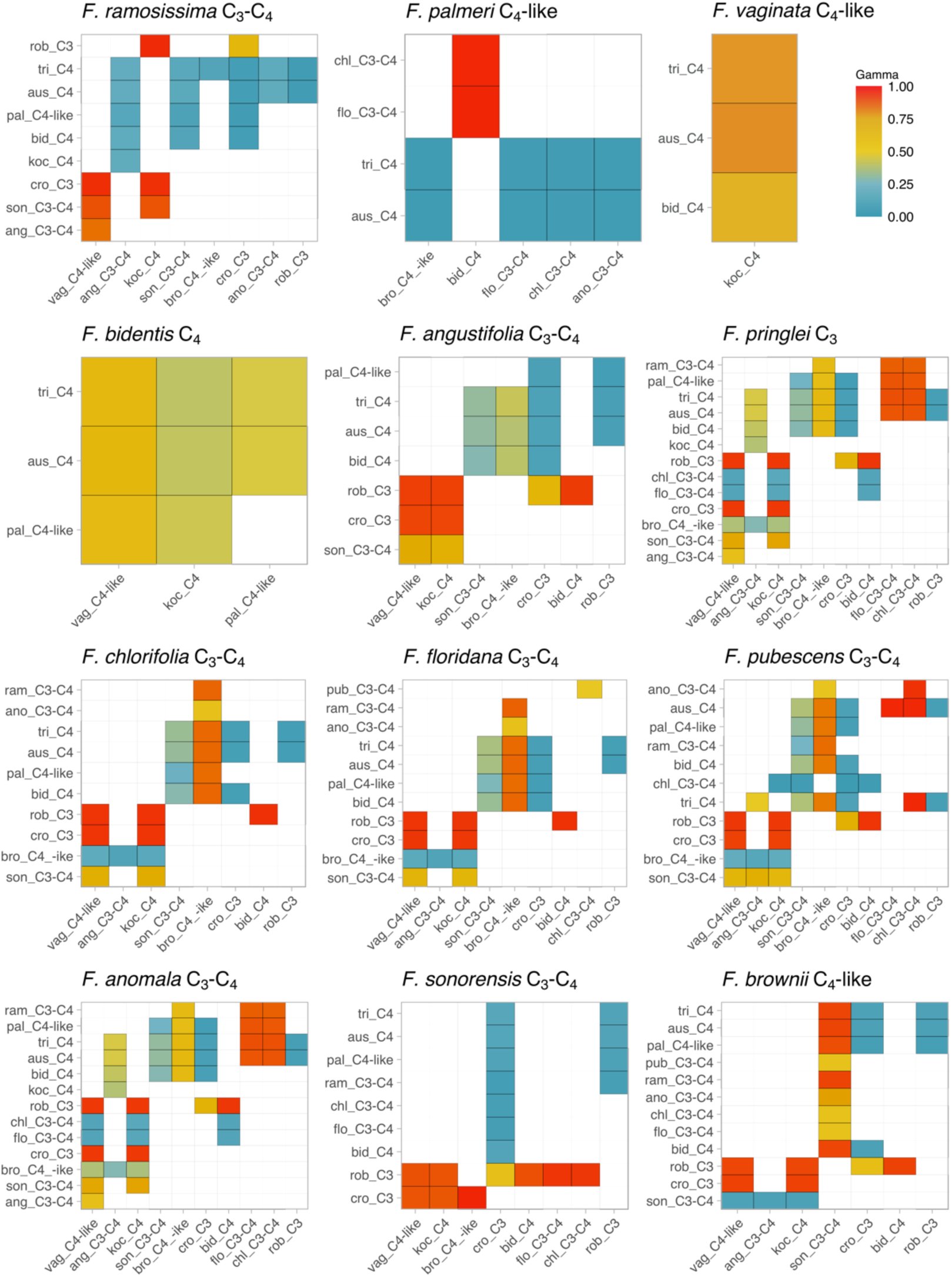
HyDe significant hybridization test for the 12 species of *Flaveria* identified as potential hybrids. Species on the x-axis are parental lineage 1 (P1) and species on the y-axis are parental lineage 2 (P2). Only colored boxes denote possible combinations of P1 and P2 as parents of hybrid species. The color scale represents the value of the hybridization parameter γ for each hybridization event. Recent 50:50 hybrids would be represented by a γ ∼0.5. Values of γ approaching 0 indicate a major hybrid contribution from P1, and values approaching 1 indicate a major hybrid contribution from P2, with both cases representing ancient hybridization.

In these three cases γ was consistent with ancient hybridization. *Flaveria vaginata* had notably fewer potential parental lineages suggesting a more recent reticulation event. The fourth species, *F. bidentis*, was the only C_4_ member of the clade which was detected as a hybrid. In this case, all potential parental lineages are members of Clade A, and γ values around 0.5 suggested a recent reticulation event. Outside clades A and B, *F. angustifolia* (C_3_-C_4_) and *F. pringlei* (C_3_) were detected as hybrids. In both cases, the potential parental lineages come from across *Flaveria* with γ values suggesting ancient reticulation. Overall, HyDe detected all C_3_-C_4_ and C_4_-like species as well as *F. bidentis* (C_4_) and *F. pringlei* (C_3_) as potential ancient hybrids.

### Lack of whole genome duplications

The orthogroup mapping did not reveal any node in *Flaveria* with elevated levels of gene duplication (Fig S7), showing the absence of whole genome duplication in the genus. In part this is expected as *Flaveria* is consistently diploid (see Powell, 1978 and ref. therein) with a haploid chromosome number of *n* = 18. Furthermore, the lack of whole genome duplication in *Flaveria* suggests that reticulation events in this group are homoploid hybridizations.

## DISCUSSION

In C_4_ photosynthesis, high rates of net photosynthesis and a highly competitive water and nitrogen use efficiency are achieved by spatially separated carbon fixation in the outer mesophyll cells preceding the Calvin-Benson cycle, and by effectively fueling Rubisco with always high CO_2_ concentration in the controlled seclusion of the Kranz cells (Long, 1999). Resulting low levels of photorespiration make C_4_ plants competitive in various stressful environments where carbon deficiency poses a problem to C_3_ plants (Sage et al., 2012). C_4_ evolved more than 60 times in angiosperms with hotspots of C_4_ origins in Poaceae and Amaranthaceae (Sage et al., 2018). Due to the anatomical and gene regulatory complexity of the C_4_ pathway, it seems clear that there must have been intermediate stable phenotypes during the evolution of C_4_ from a C_3_ ancestor (Monson and Moore, 1989). The established generalized model of C_4_ evolution tries to explain the sequence of intermediate adaptive events during the transition from the ancestral C_3_ to C_4_ using the naturally occurring C_3_-C_4_ intermediate phenotypes of *Flaveria* (and other lineages) as proxies of intermediate stages in C_4_ evolution (Sage et al., 2014; Bräutigam and Gowik, 2016). However, *Flaveria* is also known for rampant interfertility of its 21 (mostly diploid) species (Long and Rhamstine, 1968; Powell, 1978), and for the transferability of photosynthetic traits to hybrid offspring in hybridization experiments (Apel et al., 1988; Kadereit et al., 2017 and ref. therein). Therefore, a clear understanding of the phylogenetic history of this model lineage of C_4_ evolution, including tests for possible reticulate evolution, is fundamental as this might have strong implications for our understanding of the evolution of C_4_ photosynthesis in general.

Our analyses using transcriptome data of 17 *Flaveria* species revealed rampant gene tree discordance along the backbone of the phylogeny as well as the clades containing most C_3_-C_4_ intermediate and C_4_-like species (clades A and B; visualized in the cloudogram in Fig. 1). Coalescence simulations showed that gene tree discordance in *Flaveria* cannot be attributed to ILS alone. Our initial species network analyses including all 17 species sampled consistently identified *Flaveria brownii* (C_4_-like) and *F. pringlei* (C_3_) as hybrids (Figs. 2 and 3). After exclusion of these two most likely recent hybrids, deep reticulation events were recovered more clearly. The network analyses of the remaining species suggested an early reticulation event between two C_3_ lineages (*F. robusta* and *F. cronquistii*) giving rise to the ancestor of the lineage containing all C_3_-C_4_ intermediate species, C_4_-like species and C_4_ species. This result is supported by the HyDe analysis which identified all C_3_-C_4_ and C_4_-like species as potential ancient hybrids. The ancestors of *F. robusta* and *F. cronquistii* seem to have contributed equally to the origin of the hybrid lineage (Fig. 2). Within this lineage there exist two parental lineages, *F. angustifolia* (C_3_-C_4_) and *F. sonorensis* (C_3_-C_4_), that seem to have contributed to a robust clade containing all C_4_ species plus *F. ramosissima* (C_3_-C_4_) and *F. vaginata* (C_4_-like; clade A in Fig. 1), and also to another robust clade of four C_3_-C_4_ intermediate species (*F. pubescens, F. chlorifolia, F. floridana, F. anomala;* partial clade B in Fig. 1; supported also by the HyDe analysis). Of these four species, *F. anomala* was introgressed by the ancestor of clade A (Fig. 2).

These results have several important implications that will be discussed in the following three sections: 1) natural C_3_-C_4_ intermediates species in *Flaveria* do not seem to result from hybridization between a C_3_ and a C_4_ lineage (as similar C_3_-C_4_ intermediate phenotypes resulting from crossing experiments might suggest); 2) recurrent homoploid hybridization possibly played a major role in the evolution of C_4_ photosynthesis in *Flaveria* with an initial hybridization event between two C_3_ species (*F. robusta* and *F. cronquistii*) as a possible trigger, 3) only the ancestral lineages of *F. angustifolia* and *F. sonorensis* (both C_3_-C_4_ intermediates) seem to be involved in the formation of clades A and B, of which only clade A shows C_4_ photosynthesis.

Hybridization is an important factor in plant evolution and speciation in general (Abbott et al., 2013) and *Flaveria* is no exception to this but also shows hybridization in a more complex way than we expected. From the phenotypic outcome of the numerous C_3_ × C_4_ hybridization experiments in *Flaveria* it was conceivable that naturally occurring C_3_-C_4_ intermediate species might be the result of crosses between parental lineages with different photosynthetic types (Monson and Moore, 1989; Kadereit et al., 2017). However, our results indicate that origin of C_3_-C_4_ intermediates might be more complex: in *Flaveria* there are several recent hybrids expressing intermediate or deviating phenotypes compared to the parental lineages which perturb phylogenetic reconstruction (in particular, *F. pringlei* and *F. brownii*). There also were ancient hybridization events challenging the reconstruction of the backbone (Fig. 1; see below). McKown et al. (2005) and Lyu et al., 2015 interpreted *F. brownii* as representing an independent and recent origin of C_4_-like photosynthesis from C_3_-C_4_ ancestors. According to our analyses, *F. brownii* is a recent hybrid between *F. sonorensis* (C_3_-C_4_) and one of the other species in clade B (all C_3_-C_4_; Fig. 1–3), probably *F. floridana* with which it forms fertile F1 and F2 hybrids (Powell, 1978). This is an important finding for the interpretation of trait evolution in *Flaveria*. The C_4_-like photosynthesis in *F. brownii* seems to have resulted from a cross between a C_2_ type I (*F. sonorensis*) and C_2_ type 2 species (possibly *F. floridana*; see Table 1) resulting in stronger C_4_-ness of *F. brownii*. Since *F. brownii* possesses C_3_ isoforms of the key enzymes of the C_4_ pathway (e.g., Kubien et al., 2008; Gowik and Westhoff, 2011; Ludwig, 2011), the C_4_-like metabolism in *F. brownii* seems to have resulted from a highly effective gene regulatory change in comparison to its parental lineages rather than from the acquisition of C_4_ isoforms. It clearly outcompetes its parental lineages in terms of C_4_ physiology. Taniguchi et al. (2021), studying genome size evolution in *Flaveria*, showed dynamic alteration of genome size in the genus with *F. brownii* having the largest genome size among the 11 species investigated. Rewiring the parental genomes might have led to the C_4_-like phenotype found in *F. brownii* and might have facilitated adaptive divergence resulting in colonization of the coastal flats and islands of the lower Texas Gulf Coast. Similarly, our analyses revealed *F. pringlei* (C_3_ proto-kranz) as a recent hybrid between *F. cronquistii* (C_3_) and *F. angustifolia* (C_3_-C_4_; Fig. 1-3). Therefore, the proto-kranz phenotype in this species might be of different origin than that of *F. robusta* (Table 1). In fact, Lyu et al. (2015) showed the *F. pringlei* sample used in their study to be of artificial hybrid origin (possibly resulting from unintended crosses in the greenhouse). Trait assessments in these two species, e.g., of leaf anatomical and ultrastructural traits (McKown and Dengler, 2007; Sage et al., 2013) might lead to re-evaluation of their assessment as evolutionary “basal” as they likely acquired the proto-kranz phenotype differently. Also, the similarities between *F. cronquistii* and *F. pringlei* found by McKown and Dengler (2007) and their differences to *F. robusta* can now be explained. According to our analyses, *F. robusta* and *F. cronquistii* seem to be representatives of ancient C_3_ lineages in *Flaveria*, but not *F. pringlei*. The rare *Flaveria mcdougallii* (C_3_, not represented in this analysis), which is morphologically and geographically distinct and might be sister to all other species of *Flaveria* (Powell, 1978; McKown et al., 2005), would be an important species to add in further studies.

The most ancient hybridization event in *Flaveria* involved two C_3_ lineages, *F. robusta* and *F. cronquistii*. According to Powell (1978), *Flaveria cronquistii* most closely resembles *F. robusta*. The two are geographically separated, and the former is distributed in the Tehuacán Valley region and the latter in Colima and Michoacán (both Mexico). There are no reports of artificial hybridization between these two species. The resulting ancient hybrid lineage seems to include all C_3_-C_4_ intermediate, C_4_-like and C_4_ species (Fig. 2; Table 1). Concerning leaf anatomy, *Flaveria robusta* differs from *F. cronquistii* by a higher vein density achieved through higher vein branching, and by a higher number of organelles in the bundle sheath cells where the organelles are located in a centripetal position along the cell wall connecting bundle sheath cells and vascular tissue. Organelles are more evenly distributed in *F. cronquistii*, and its leaves are more succulent and show larger bundle sheath cells (McKown and Dengler, 2007; Sage et al., 2013). The combination of these traits in a hybrid lineage and subsequent segregation effectively leading to a higher bundle sheath to mesophyll ratio and activated large bundle sheath cells may have triggered the evolution of “pre kranz” cells in *Flaveria*. Possibly the strong leaf anatomical differences between the two parental lineages promoted the origin of novel traits which then allowed this lineage to occupy new niches eco-geographically separate from the parents. There are multiple examples for plant lineages in which homoploid hybridization resulted in potentially adaptive genotypes through transgressive segregation, eventually leading to speciation (Gross and Rieseberg, 2005; Nieto Feliner et al., 2020). An increase of the bundle sheath to mesophyll ratio has been suggested to play an initial role in the evolution of C_4_ in several plant lineages (Marshall et al., 2007; McKown and Dengler, 2007; Christin et al., 2013; Griffiths et al., 2013; Lauterbach et al., 2019).

Two C_3_-C_4_ intermediate lineages, *F. angustifolia* and *F. sonorensis*, are involved in the formation of clades A and B according to our results. It is remarkable that only these two and not any other species (especially not the C_4_-like species) contributed to the origin of these clades. In their extant distribution the two species do not overlap. *Flaveria angustifolia* grows in sclerophyllous scrub in Puebla and Oaxaca, while *F. sonorensis* is found only in the short-tree forests of tropical Sonora (Powell, 1978). Looking more closely at these two descendants of the lineages that seem to have hybridized in the past and possibly gave rise to C_4_ photosynthesis might give new insights into important preconditions for the evolution of C_4_ in *Flaveria*. Both *F. angustifolia* and *F. sonorensis* were categorized as C_2_ Type I species (Table 1), but with weaker C_2_ photosynthesis (relatively high amounts of glycine decarboxylase in the mesophyll cells and relatively high CO_2_ compensation points) than other C_2_ species of *Flaveria* (Sage et al., 2018). This finding implies the evolution of a C_2_ phenotype prior to C_4_, supporting the ‘photorespiratory bridge hypothesis’ (Sage et al., 2018) in case of *Flaveria*. However, it seems that C_4_ evolution in *Flaveria* was triggered by hybridization of C_2_ lineages, a scenario never suggested before. For *Flaveria* this somewhat shifts the focus to C_2_ photosynthesis and under which selective conditions this type of photosynthesis might have evolved. To gain further insights into the evolution of C_2_ photosynthesis, detailed studies of the origin, anatomy, and ecophysiology of *F. angustifolia* and *F. sonorensis* might be rewarding. Organelle enrichment and their flux-optimized positioning as well as glycine decarboxylase accumulation in the proto-kranz cells (Khoshravesh et al., 2016) seem to be essential for C_2_ photosynthesis (in eudicots and monocots). These traits enable species to more efficiently re-cycle respired CO_2_ and to longer maintain a positive assimilation rate under carbon deficient conditions. Since the majority of C_4_ lineages do not have any known C_2_ relatives (see Sage et al., 2018 for an overview), and C_2_ lineages without close C_4_ relative are known as well (see Lundgren, 2020 for an overview), the question remains whether there exist several evolutionary pathways to C_4_ photosynthesis (Edwards, 2019), and whether C_2_ should be considered an independent carbon concentrating mechanism not necessarily always intimately connected to C_4_ photosynthesis (Lundgren, 2020 and ref. therein). Finally, a C_2_ lineage might also be the result of ancient hybridization of a C_3_ and a C_4_ lineage, as has been suggested for *Salsola divaricata*, which then might have thrived through the large plasticity of photosynthetic traits inherited from photosynthetically divergent parental lineages (Tefarikis et al., 2021).

## CONCLUSIONS

The young genus *Flaveria* which includes four C_3_, four C_4_, three C_4_-like and c. ten C_3_-C_4_ intermediate species is remarkable for the high number of evolutionary reticulations creating an enormous diversity of phenotypes with different photosynthetic traits. Due to this and its strictly diploid chromosome number the genus is a highly interesting system to study the genetic basis of the C_4_ syndrome and the role of transgressive segregation in the origin of genotypes eventually leading to the evolution of C_4_ photosynthesis. We found evidence that homoploid hybridization of C_3_ lineages might have triggered the evolution of C_2_ photosynthesis, and that homoploid hybridization of C_2_ lineages gave rise to C_4_-like or C_4_ photosynthesis. In both cases the hybrid derivatives possibly surpassed the parental performance under conditions of high photorespiration and had an adaptive advantage. However, since reticulation occurred in recent as well as ancient lineages of the genus, the sequence of evolutionary events needs to be studied in the light of a carefully reconstructed phylogenetic history of the genus. Comparison of the entire genomes of parental and hybrid lineages should allow us to detect whether the origin of new photosynthetic traits is indeed the result of transgressive segregation (de los Reyes, 2019), as suggested here, or of stepwise mutational change of relevant genes.

## DATA AVAILABILITY STATEMENT

Analysis and results files can be accessed at the Dryad repository XXXX

## AUTHOR CONTRIBUTIONS

GK and DFM-B conceived the study. DFM-B performed all analyses. GK and DFM-B evaluated the results and wrote the paper.

## ACKNOWLEDGMENTS

We thank Joachim W. Kadereit (Mainz) for valuable comments on the manuscript.

## SUPPLEMENTARY MATERIAL

**Table S1**. Taxon sampling and source of data

**Table S2**. HyDe significant hybridization tests for all species of *Flaveria*.

**Table S3**. HyDe significant hybridization tests for all species of *Flaveria* excluding *F. pringlei* (C_3_).

**Fig. S1**. A. Maximum likelihood phylogeny of *Flaveria* inferred with IQ-TREE from the concatenated 1249-nuclear gene supermatrix. Numbers above branches represent bootstrap support (BS). Branch lengths as substitutions per site (scale bar on the bottom). B. ASTRAL tree of *Flaveria* inferred from the 1,295 nuclear gene trees. Local posterior probabilities (LLP) are shown next to nodes. Internal branch lengths are in coalescent units (scale bar on the bottom).

**Fig. S2**. A. Maximum likelihood cladogram of *Flaveria* inferred with IQ-TREE from the concatenated 1249-nuclear gene supermatrix. B. ASTRAL cladogram of *Flaveria* inferred from the 1,295 nuclear gene trees. Pie charts represent the proportion of gene trees that support that clade (blue), the main alternative bifurcation (green), the remaining alternatives (red), and conflict or support that have <50% bootstrap support (gray). Number above and below branches represent the number of concordant and discordant informative gene trees, respectively.

**Fig. S3**. A. Maximum likelihood cladogram of *Flaveria* inferred with IQ-TREE from the concatenated 1249-nuclear gene supermatrix. B. ASTRAL cladogram of *Flaveria* inferred from the 1,295 nuclear gene trees. Quartet Sampling (QS) scores are shown above branches. QS scores: Quartet concordance/Quartet differential/Quartet informativeness. Circles at nodes are colored by quartet concordance support.

**Fig. S4**. Distribution of tree-to-tree distances between empirical gene trees and the ASTRAL tree, compared to the distribution of tree-to-tree distances between simulated trees and the ASTRAL tree.

**Fig. S5**. Maximum pseudo-likelihood scores for species networks inferred with PhyloNet using the (A) 18-taxon, (B) and 16-taxon data sets. The x-axis notes the maximum number of reticulations for each of the network searches allowing up to ten reticulation events.

**Fig. S6**. Maximum pseudo-likelihood species networks inferred with PhyloNet using the (A) 18-taxon, (B) and 16-taxon data sets and allowing up to ten reticulation events. Red and blue curved branches indicate the minor and major edges, respectively of hybrid nodes. Numbers next to curved branches indicate inheritance probabilities for each hybrid node.

**Fig. S7**. Maximum likelihood cladogram of *Flaveria* inferred with IQ-TREE from the concatenated 1249-nuclear gene supermatrix. Numbers above branches are gene duplication counts and numbers below branches are gene duplication percentages.

