## Supplemental material for "Exploring the possible role of hybridization in the evolution of photosynthetic pathways in *Flaveria* (Asteraceae), the prime model of C_4_ photosynthesis evolution"

### SUPPLEMENTARY MATERIAL

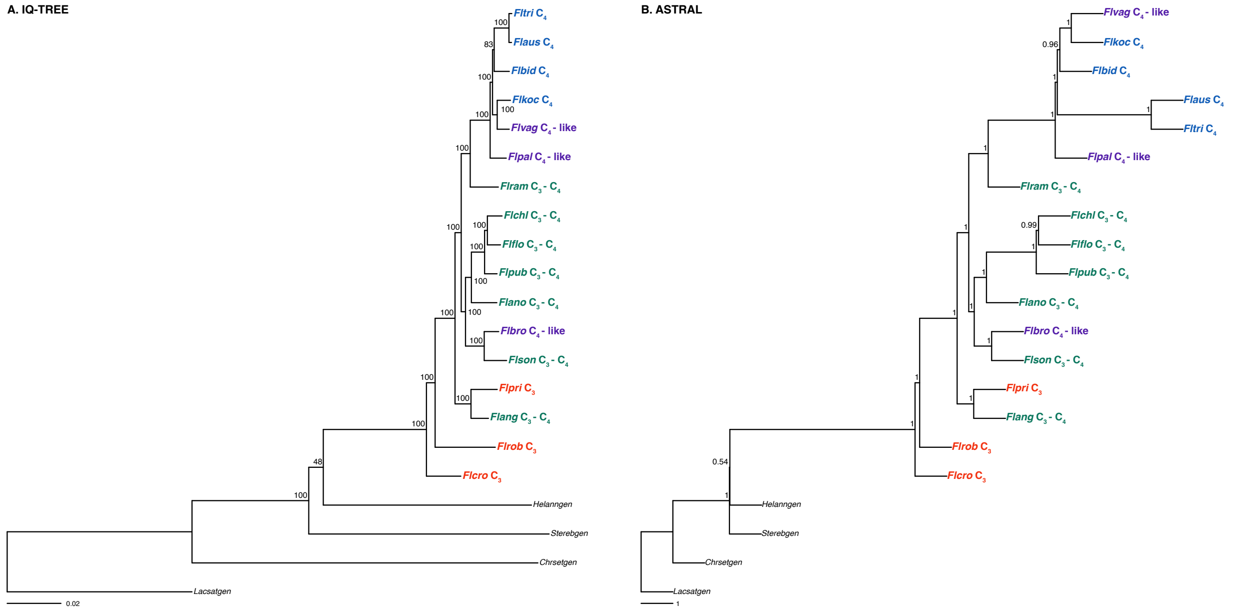


**Fig. S1.** A. Maximum likelihood phylogeny of *Flaveria* inferred with IQ-TREE from the concatenated 1249-nuclear gene supermatrix. Numbers above branches represent bootstrap support (BS). Branch lengths as substitutions per site (scale bar on the bottom). B. ASTRAL tree of *Flaveria* inferred from the 1,295 nuclear gene trees. Local posterior probabilities (LLP) are shown next to nodes. Internal branch lengths are in coalescent units (scale bar on the bottom).

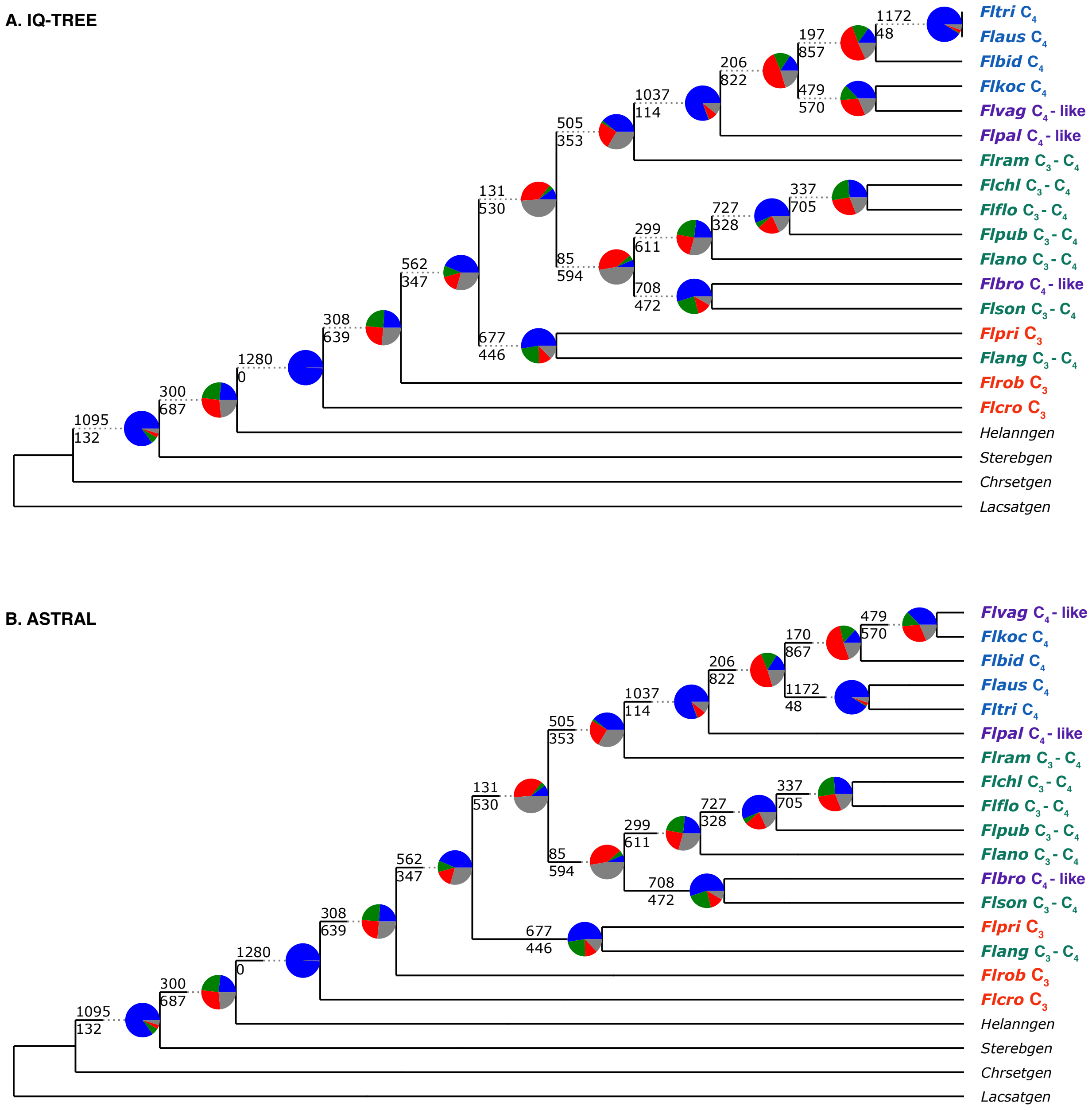


**Fig. S2.** A. Maximum likelihood cladogram of *Flaveria* inferred with IQ-TREE from the concatenated 1249-nuclear gene supermatrix. B. ASTRAL cladogram of *Flaveria* inferred from the 1,295 nuclear gene trees. Pie charts represent the proportion of gene trees that support that clade (blue), the main alternative bifurcation (green), the remaining alternatives (red), and conflict or support that have <50% bootstrap support (gray). Number above and below branches represent the number of concordant and discordant informative gene trees, respectively.

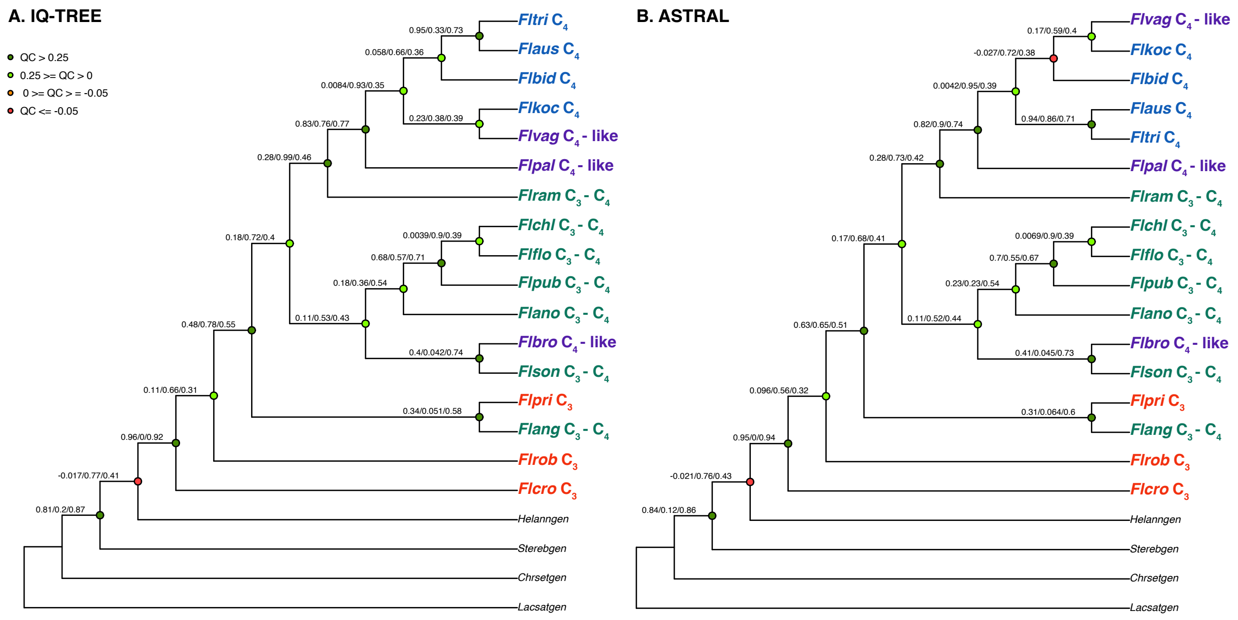


**Fig. S3.** A. Maximum likelihood cladogram of *Flaveria* inferred with IQ-TREE from the concatenated 1249-nuclear gene supermatrix. B. ASTRAL cladogram of *Flaveria* inferred from the 1,295 nuclear gene trees. Quartet Sampling (QS) scores are shown above branches. QS scores: Quartet concordance/Quartet differential/Quartet informativeness. Circles at nodes are colored by quartet concordance support.

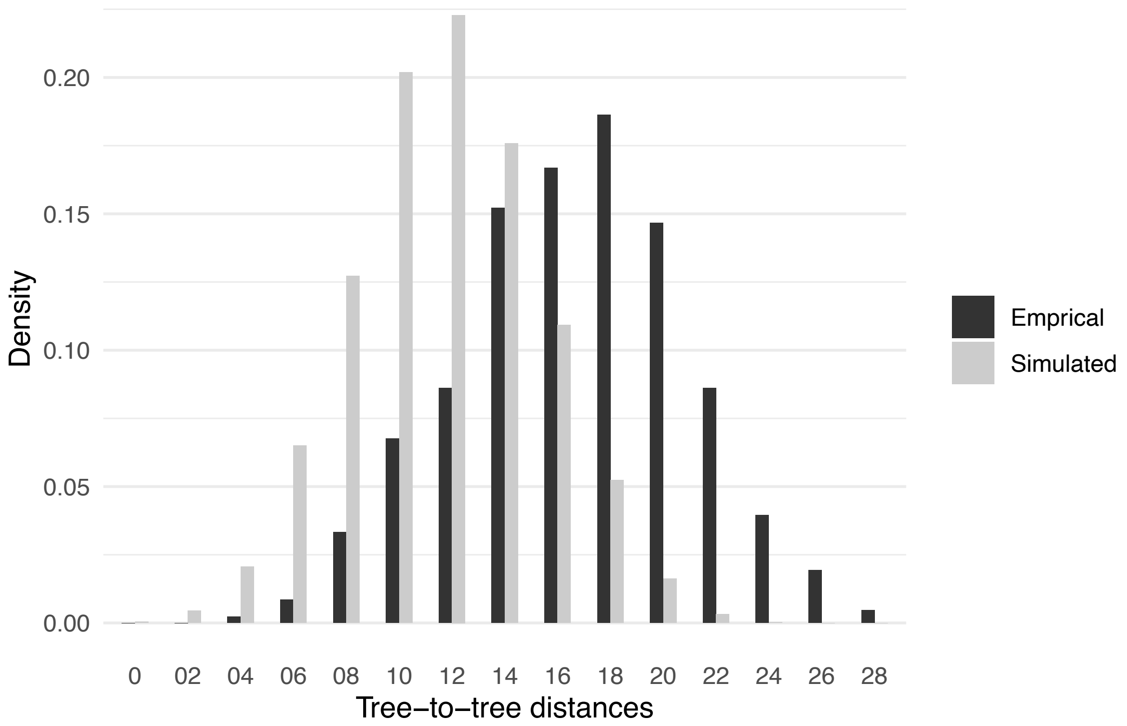


**Fig. S4.** Distribution of tree-to-tree distances between empirical gene trees and the ASTRAL tree, compared to the distribution of tree-to-tree distances between simulated trees and the ASTRAL tree.

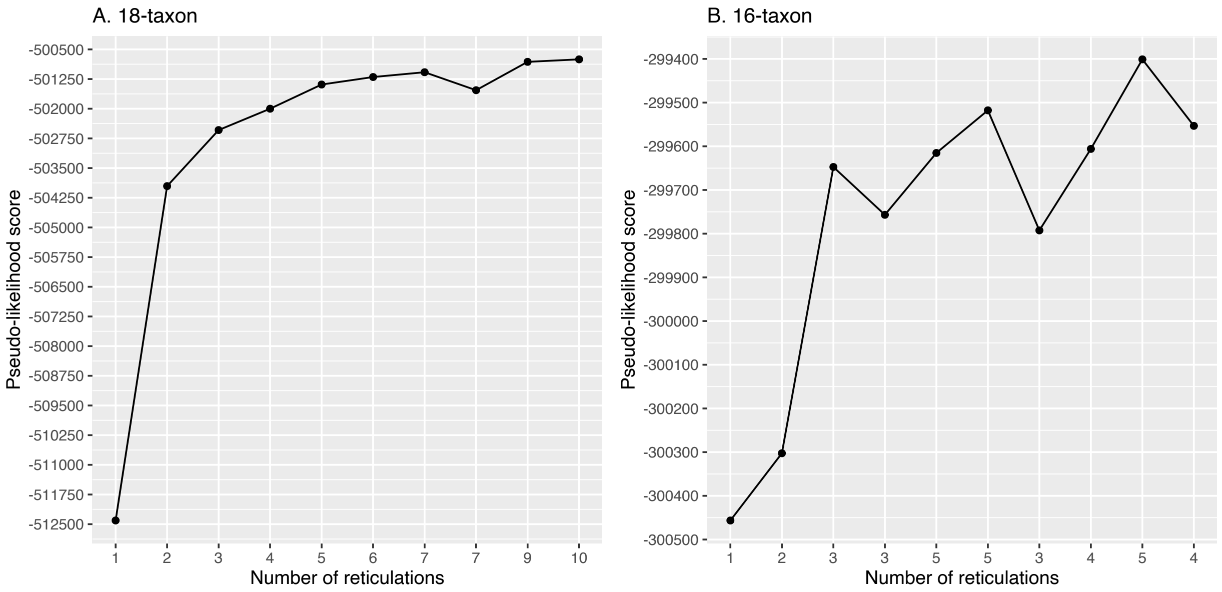


**Fig. S5.** Maximum pseudo-likelihood scores for species networks inferred with PhyloNet using the (A) 18-taxon and (B) 16-taxon data sets. The x-axis notes the maximum number of reticulations for each of the network searches allowing up to ten reticulation events.

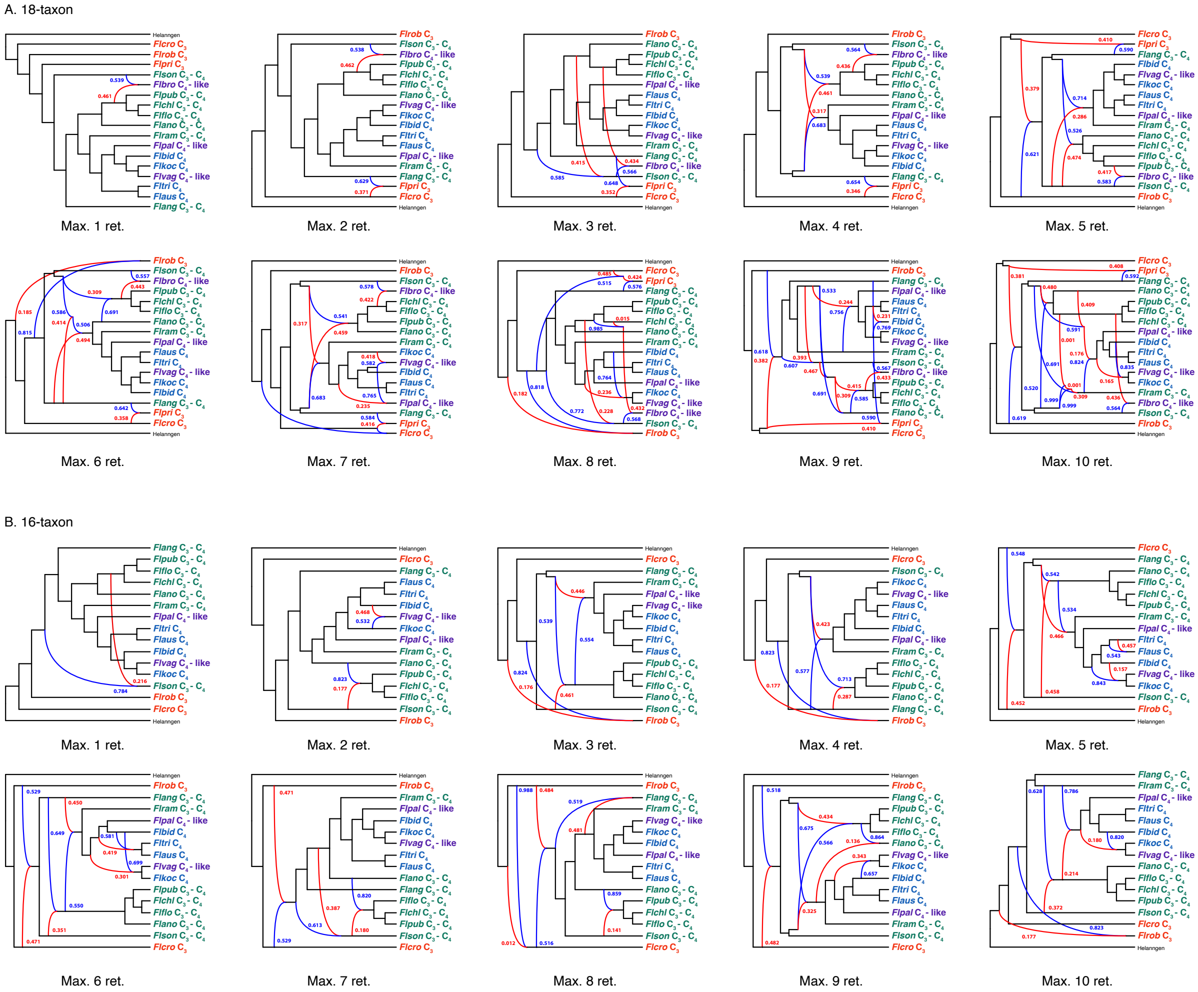


**Fig. S6.** Maximum pseudo-likelihood species networks inferred with PhyloNet using the (A) 18-taxon, (B) 16-taxon, and (C) 11-taxon data sets and allowing up to ten reticulation events. Red and blue curved branches indicate the minor and major edges, respectively of hybrid nodes. Numbers next to curved branches indicate inheritance probabilities for each hybrid node.

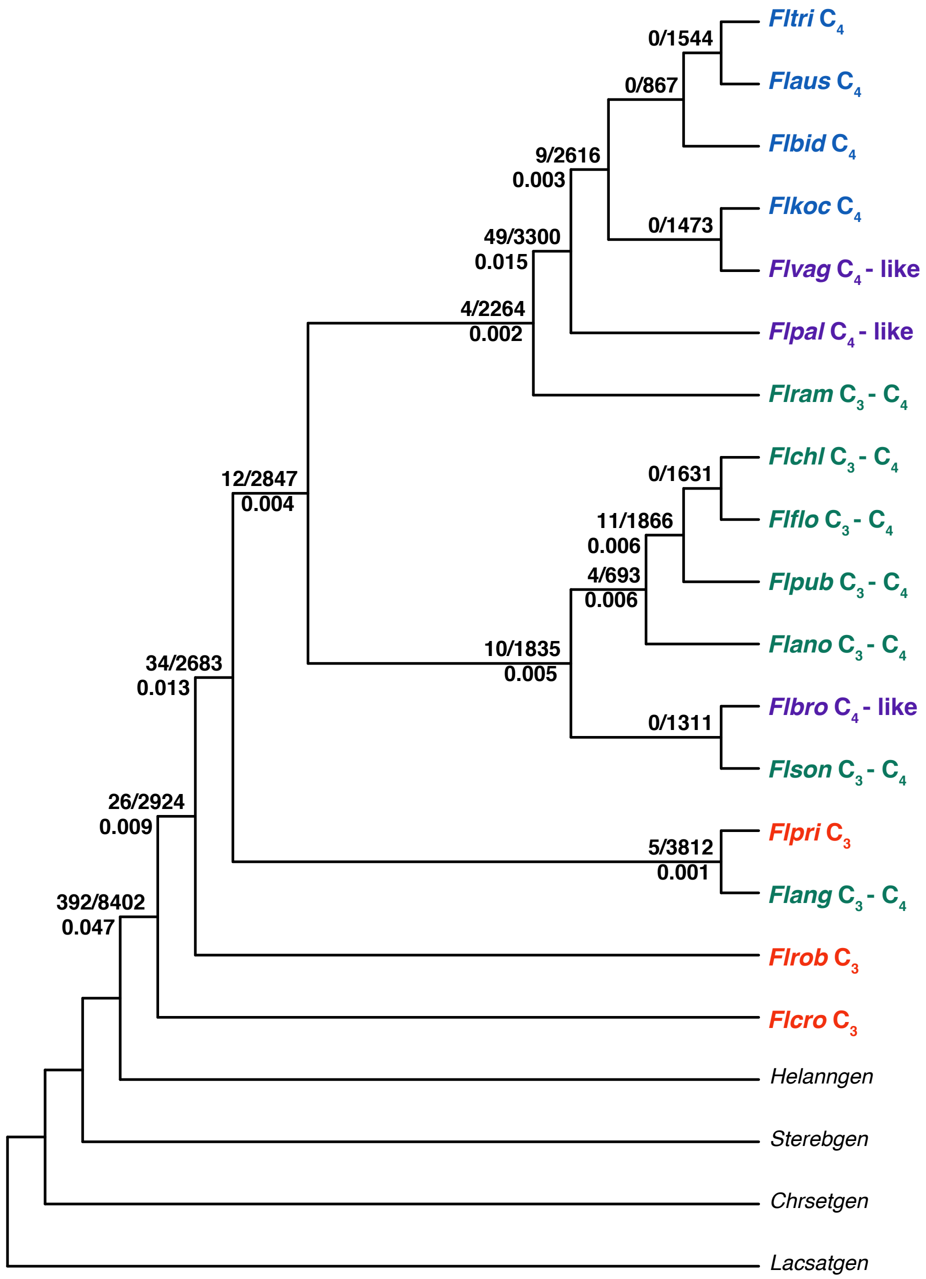


**Fig. S7.** Maximum likelihood cladogram of *Flaveria* inferred with IQ-TREE from the concatenated 1249-nuclear gene supermatrix. Numbers above branches are gene duplication counts and numbers below branches are gene duplication percentages.

**Table S1**. Taxon sampling and source of data

| Species | Code | Accession number | Library reads | Stranded or non-stranded library | Source | Source reference | Reference coverage for transcriptomes against *Helianthus annuus* | No. of final CDS used for BLASTN | No. orthologs | Perc. orthologs (total 5374) |
|---|---|---|---|---|---|---|---|---|---|---|
| *Flaveria angustifolia* | Flang | ERR2040662, ERR2040663 | paired end | non-stranded | NCBI-SRA | doi: 10.1093/gigascience/giz126 | 26% | 22543 | 4789 | 89.11 |
| *Flaveria anomala* | Flano | SRR1165193 | single end | non-stranded | NCBI-SRA | doi: 10.7554/eLife.02478 | 32% | 25189 | 4985 | 92.76 |
| *Flaveria australasica* | Flaus | SRR1692297 | single end | non-stranded | NCBI-SRA | doi: 10.1186/s12862-015-0399-9 | 29% | 25830 | 4966 | 92.41 |
| *Flaveria bidentis* | Flbid | ERR2040664, ERR2040665 | paired end | non-stranded | NCBI-SRA | doi: 10.1093/gigascience/giz126 | 29% | 23015 | 4984 | 92.74 |
| *Flaveria brownii* | Flbro | ERR2040666, ERR2040667 | paired end | non-stranded | NCBI-SRA | doi: 10.1093/gigascience/giz126 | 27% | 27927 | 4914 | 91.44 |
| *Flaveria chlorifolia* | Flchl | SRR1165206 | single end | non-stranded | NCBI-SRA | doi: 10.7554/eLife.02478 | 31% | 24872 | 4878 | 90.77 |
| *Flaveria cronquistii* | Flcro | ERR2040668, ERR2040669 | paired end | non-stranded | NCBI-SRA | doi: 10.1093/gigascience/giz126 | 27% | 25347 | 4682 | 87.12 |
| *Flaveria floridana* | Flflo | SRR1691226 | single end | non-stranded | NCBI-SRA | doi: 10.1186/s12862-015-0399-9 | 31% | 25228 | 4911 | 91.38 |
| *Flaveria kochiana* | Flkoc | ERR2040670 | paired end | non-stranded | NCBI-SRA | doi: 10.1093/gigascience/giz126 | 23% | 20827 | 4538 | 84.44 |
| *Flaveria palmeri* | Flpal | ERR2040671 | paired end | non-stranded | NCBI-SRA | doi: 10.1093/gigascience/giz126 | 25% | 22669 | 4833 | 89.93 |
| *Flaveria pringlei* | Flpri | ERR2040672, ERR2040673 | paired end | non-stranded | NCBI-SRA | doi: 10.1093/gigascience/giz126 | 27% | 30052 | 4823 | 89.75 |
| *Flaveria pubescens* | Flpub | ERR2040674, ERR2040675 | paired end | non-stranded | NCBI-SRA | doi: 10.1093/gigascience/giz126 | 28% | 24071 | 4896 | 91.11 |
| *Flaveria ramosissima* | Flram | SRR11124423 | paired end | non-stranded | NCBI-SRA | doi: 10.3389/fpls.2020.00935 | 35% | 24940 | 4927 | 91.68 |
| *Flaveria robusta* | Flrob | SRR11124411 | paired end | non-stranded | NCBI-SRA | doi: 10.3389/fpls.2020.00935 | 36% | 24557 | 4773 | 88.82 |

| | | | | | | | | | | |
|---|---|---|---|---|---|---|---|---|---|---|
| *Flaveria sonorensis* | Flson | SRR11124417 | paired end | non-stranded | NCBI-SRA | doi: 10.3389/fpls.2020.00935 | 35% | 24917 | 4490 | 83.55 |
| *Flaveria trinervia* | Fltri | SRR11124428 | paired end | non-stranded | NCBI-SRA | doi: 10.3389/fpls.2020.00935 | 35% | 23029 | 4875 | 90.71 |
| *Flaveria vaginata* | Flvag | ERR2040679 | paired end | non-stranded | NCBI-SRA | doi: 10.1093/gigascience/giz126 | 25% | 23596 | 4843 | 90.12 |
| *Chrysanthemum seticuspe* | Chrsetgen | Genome | N/A | N/A | http://mum-garden.kazusa.or.jp/ | doi: 10.1093/dnares/dsy048 | N/A | 71057 | 4213 | 78.40 |
| *Helianthus annuus* | Helanngen | Genome | N/A | N/A | Phytozome | doi: 10.1038/nature22380 | N/A | 52243 | 5145 | 95.74 |
| *Lactuca sativa* | Lacsatgen | Genome | N/A | N/A | Phytozome | doi: 10.1038/ncomms14953 | N/A | 38910 | 4487 | 83.49 |
| *Stevia rebaudiana* | Sterebgen | Genome | N/A | N/A | doi: 10.6084/m9.figshare.14169491.v1 | doi: 10.1038/s41438-021-00565-4 | N/A | 44112 | 5035 | 93.69 |

**Table S2**. HyDe significant hybridization tests for all species of *Flaveria*.

| P1 | Hybrid | P2 | Zscore | Pvalue | Gamma | AAAA | AAAB | AABA | AABB | AABC | ABAA | ABAB | ABAC | ABBA | BAAA | ABBC | CABC | BACA | BCAA | ABCD |
|---|---|---|---|---|---|---|---|---|---|---|---|---|---|---|---|---|---|---|---|---|
| vag_C4-like | ang_C3-C4 | pri_C3 | 39.3388236 | 0 | 0.35052831 | 1036100.25 | 5494.25 | 4275.25 | 7177.25 | 96.5 | 13175.75 | 1184.25 | 134 | 4418.75 | 98577.75 | 1167 | 219.75 | 580.75 | 2512.25 | 46.25 |
| vag_C4-like | ang_C3-C4 | son_C3-C4 | 4.21720008 | 1.24E-05 | 0.75901639 | 995854.75 | 9264.75 | 7812.75 | 3149 | 176 | 10981 | 2855 | 214 | 3781 | 95804.75 | 1509.5 | 237.5 | 1224 | 1713 | 68 |
| vag_C4-like | ang_C3-C4 | cro_C3 | 8.21280974 | 1.11E-16 | 0.9274543 | 1026894.94 | 11389.0625 | 8426.25 | 2836.75 | 216 | 11950.75 | 2266.5 | 282 | 9556.8125 | 91860.8125 | 2431.125 | 419.5 | 1189 | 1687.5 | 87 |
| vag_C4-like | ang_C3-C4 | rob_C3 | 6.94178987 | 1.95E-12 | 0.93796621 | 992896.5 | 16035.5 | 8111 | 2827.5 | 326.5 | 11348 | 2341 | 374 | 9697 | 88368.5 | 3040.5 | 419.5 | 1137 | 1596.5 | 110 |
| koc_C4 | ang_C3-C4 | pri_C3 | 39.1901793 | 0 | 0.35067791 | 1028441.44 | 5441.5 | 4262.3125 | 7113.75 | 97 | 13030.3125 | 1172.25 | 126 | 4381.0625 | 97962.25 | 1159.5 | 217.875 | 581.75 | 2365.75 | 44.25 |
| koc_C4 | ang_C3-C4 | son_C3-C4 | 4.6196644 | 1.92E-06 | 0.73230656 | 988616.313 | 9154.375 | 7790.0625 | 3146.75 | 175 | 10916.9375 | 2827.375 | 204.75 | 3701.0625 | 95207.75 | 1495.5 | 233.625 | 1187 | 1627.5 | 60 |
| koc_C4 | ang_C3-C4 | cro_C3 | 8.2091839 | 1.11E-16 | 0.92597499 | 1012797.31 | 11146.875 | 8349.5625 | 2799.25 | 208 | 11796.8125 | 2233.5 | 253 | 9310.4375 | 90740.875 | 2404.25 | 418.625 | 1129 | 1557.5 | 83 |
| koc_C4 | ang_C3-C4 | rob_C3 | 7.34363236 | 1.05E-13 | 0.93510788 | 986353.188 | 15976 | 8053.3125 | 2812.25 | 311.5 | 11289.8125 | 2300.75 | 362.5 | 9671.5625 | 87768 | 3049.5 | 419.625 | 1111 | 1489 | 92 |
| pri_C3 | ang_C3-C4 | son_C3-C4 | 35.0167372 | 0 | 0.69394435 | 1060238.75 | 11431 | 4813.75 | 4097.5 | 168.75 | 5505.5 | 1386 | 141 | 7534 | 101136.5 | 2216.25 | 107 | 679.25 | 1115.5 | 38.25 |
| pri_C3 | ang_C3-C4 | bro_C4-like | 37.6560343 | 0 | 0.66935329 | 1083002.25 | 11672 | 4793 | 4286.5 | 170.25 | 5738.25 | 1271.25 | 133 | 7375.25 | 103235.5 | 2322.25 | 102.5 | 653.75 | 1194 | 36.25 |
| pri_C3 | ang_C3-C4 | bid_C4 | 40.8945142 | 0 | 0.64910035 | 1116645.75 | 14336 | 4633 | 4774 | 219 | 6000.25 | 1278.25 | 136 | 7744.75 | 106342 | 2663 | 100.5 | 626.5 | 1264 | 41 |
| pri_C3 | ang_C3-C4 | flo_C3-C4 | 39.5766743 | 0 | 0.64367141 | 1119997.25 | 12173.75 | 4818 | 4614.75 | 177.75 | 5954.25 | 1278 | 114 | 7305.5 | 106379 | 2380.75 | 103.5 | 643.25 | 1226 | 37.25 |
| pri_C3 | ang_C3-C4 | chl_C3-C4 | 39.8196614 | 0 | 0.64624615 | 1120748 | 12447.25 | 4830.75 | 4638.75 | 180.25 | 5986.5 | 1278 | 113 | 7417.5 | 106536.75 | 2394.75 | 106.5 | 646.25 | 1222.5 | 36.25 |
| pri_C3 | ang_C3-C4 | ano_C3-C4 | 39.864224 | 0 | 0.63966301 | 1120789.25 | 10952.5 | 4727 | 4610.5 | 184.25 | 5950.25 | 1242.25 | 122 | 7221.5 | 106971.25 | 2288.75 | 102.5 | 662.75 | 1211 | 35.25 |
| pri_C3 | ang_C3-C4 | ram_C3-C4 | 40.4319606 | 0 | 0.6343049 | 1081497.5 | 11132 | 4492.5 | 4540.75 | 145.75 | 5840.25 | 1137.5 | 108.5 | 7040.5 | 103224.5 | 2230.25 | 93 | 601.75 | 1203 | 37.25 |
| pri_C3 | ang_C3-C4 | pub_C3-C4 | 40.21428 | 0 | 0.63704215 | 1111280.5 | 11708 | 4725.75 | 4644.25 | 177.25 | 6009 | 1228 | 116 | 7224 | 106182 | 2304.75 | 104.5 | 634.25 | 1248.5 | 37.25 |
| pri_C3 | ang_C3-C4 | pal_C4-like | 39.5338366 | 0 | 0.64948701 | 1063956 | 13453 | 4404 | 4522.5 | 208.25 | 5680.25 | 1234.25 | 137 | 7327.25 | 101369.75 | 2490.25 | 97.5 | 605.75 | 1195 | 39.25 |
| pri_C3 | ang_C3-C4 | aus_C4 | 40.1973998 | 0 | 0.65559526 | 1089415.75 | 14082 | 4462 | 4570.5 | 215.25 | 5702.25 | 1221.25 | 135 | 7596.75 | 102744.5 | 2595.25 | 104.5 | 590.75 | 1184 | 42.25 |
| pri_C3 | ang_C3-C4 | tri_C4 | 40.022285 | 0 | 0.65613197 | 1084874 | 14326.25 | 4509 | 4604.25 | 218.25 | 5771.25 | 1258.5 | 141.5 | 7642.5 | 103519.75 | 2623.75 | 104.5 | 594.75 | 1219.5 | 41.25 |
| son_C3-C4 | ang_C3-C4 | bid_C4 | 5.29507184 | 5.96E-08 | 0.28063096 | 1086819.25 | 12276 | 8571.5 | 4055.25 | 244 | 10080.75 | 3074.75 | 204.5 | 3457.25 | 104907.75 | 1850.5 | 186 | 1306 | 1652.5 | 71 |
| son_C3-C4 | ang_C3-C4 | aus_C4 | 6.49024782 | 4.31E-11 | 0.30777842 | 1060536.44 | 11952.0625 | 8270.5 | 3991 | 242 | 9862.0625 | 2947.5625 | 214.125 | 3411.5 | 101080.75 | 1816 | 186 | 1223 | 1580 | 67 |
| son_C3-C4 | ang_C3-C4 | tri_C4 | 6.59609589 | 2.12E-11 | 0.31435399 | 1063385.88 | 12289.625 | 8374.5 | 4054 | 241 | 9864.375 | 3013.875 | 226.75 | 3490.75 | 102737.75 | 1844.5 | 184 | 1276 | 1638 | 72 |
| bro_C4-like | ang_C3-C4 | bid_C4 | 4.09419512 | 2.12E-05 | 0.39649578 | 1099188 | 12309.25 | 8622.75 | 3732 | 235 | 9940 | 3267 | 225.5 | 3572.5 | 105763.5 | 1862 | 212 | 1377.5 | 1596 | 75 |
| bro_C4-like | ang_C3-C4 | aus_C4 | 4.644361 | 1.71E-06 | 0.38559322 | 1074325.25 | 11993 | 8386.75 | 3704 | 223 | 9703.25 | 3160.25 | 241 | 3501.5 | 102302.5 | 1828 | 194 | 1317.5 | 1547 | 64 |
| bro_C4-like | ang_C3-C4 | tri_C4 | 4.97281764 | 3.30E-07 | 0.42259414 | 1068437.75 | 12232.5 | 8403.75 | 3718 | 219 | 9703.5 | 3200.5 | 252.5 | 3579.25 | 102860.25 | 1837.5 | 195 | 1341.5 | 1590 | 66 |

| | | | | | | | | | | | | | | | | | | | | |
|---|---|---|---|---|---|---|---|---|---|---|---|---|---|---|---|---|---|---|---|---|
| cro_C3 | ang_C3-C4 | bid_C4 | 9.02138585 | 0 | 0.07679948 | 1102867.19 | 13131.75 | 9135 | 10207.3125 | 450 | 12267.0625 | 2414.75 | 298.5 | 3063 | 99119.8125 | 1781 | 228 | 1259 | 2626.625 | 86 |
| cro_C3 | ang_C3-C4 | rob_C3 | 5.16559135 | 1.20E-07 | 0.69780843 | 1048165.13 | 15326.5 | 13709.75 | 4670.125 | 494.75 | 9781.375 | 4237.5 | 383.25 | 5236.5 | 91674.875 | 2281 | 346.25 | 1979 | 1607.75 | 103.25 |
| cro_C3 | ang_C3-C4 | pal_C4-like | 7.3066077 | 1.38E-13 | 0.06499929 | 1047456.13 | 12128 | 8612.5 | 9751.375 | 436 | 11683.875 | 2354 | 273 | 2868.25 | 93871.625 | 1619.5 | 213 | 1207 | 2526.75 | 67 |
| cro_C3 | ang_C3-C4 | aus_C4 | 8.81849196 | 0 | 0.07721735 | 1076612.38 | 12736.75 | 8839.25 | 9887.625 | 434.5 | 11829.875 | 2379.75 | 281.5 | 3008 | 95635.625 | 1750 | 215 | 1193 | 2518.75 | 86 |
| cro_C3 | ang_C3-C4 | tri_C4 | 9.42926082 | 0 | 0.0819511 | 1072816.63 | 13040.5 | 8889 | 9997.625 | 433 | 11930.375 | 2408 | 290 | 3085.5 | 96458.125 | 1766 | 216 | 1217 | 2572.25 | 83 |
| bid_C4 | ang_C3-C4 | rob_C3 | 6.60007401 | 2.07E-11 | 0.94318365 | 1084336.94 | 17598.5625 | 8940.75 | 3067.5 | 339.5 | 12651.0625 | 2583.0625 | 390.125 | 10625 | 96868 | 3328 | 456 | 1235 | 1710.5 | 111 |
| rob_C3 | ang_C3-C4 | pal_C4-like | 4.82841354 | 6.89E-07 | 0.0431795 | 1021572.44 | 11564.0625 | 8323.25 | 10103.25 | 431 | 16562.8125 | 2470.8125 | 367.625 | 2815.25 | 91032.5 | 1556 | 331.5 | 1172 | 3149.5 | 93 |
| rob_C3 | ang_C3-C4 | aus_C4 | 7.13488409 | 4.88E-13 | 0.06208366 | 1059362 | 12371.75 | 8636 | 10321.25 | 453.5 | 17071 | 2507 | 359 | 3024.25 | 93445.75 | 1687 | 338.5 | 1155 | 3212 | 105 |
| rob_C3 | ang_C3-C4 | tri_C4 | 6.87647553 | 3.09E-12 | 0.05990988 | 1062773 | 12704.75 | 8754 | 10513 | 454.5 | 17230.75 | 2584.75 | 366.5 | 3090 | 94960.25 | 1716.5 | 344.5 | 1206 | 3279.5 | 104 |
| vag_C4-like | ano_C3-C4 | ang_C3-C4 | 4.89616627 | 4.89E-07 | 0.61034965 | 1052360.5 | 8262 | 7975.5 | 3266 | 185 | 11092 | 2917.75 | 219.5 | 3463.25 | 101604 | 1389.5 | 217 | 1394 | 1756 | 67 |
| vag_C4-like | ano_C3-C4 | pri_C3 | 6.67822416 | 1.22E-11 | 0.88863676 | 1057894.75 | 9762 | 8262.5 | 3059 | 205 | 11474.5 | 2580.5 | 242 | 6398.75 | 98675.5 | 1896.5 | 331 | 1315 | 1722 | 78 |
| vag_C4-like | ano_C3-C4 | son_C3-C4 | 6.6617848 | 1.36E-11 | 0.76266838 | 1039309.75 | 9706.5 | 7983.5 | 3144 | 191 | 11174 | 2681.5 | 226 | 4167.75 | 99946.5 | 1600.5 | 254 | 1333 | 1743 | 57 |
| vag_C4-like | ano_C3-C4 | bro_C4-like | 13.5551011 | 0 | 0.37566599 | 1040186 | 8666.75 | 7132.75 | 3954 | 182 | 11528 | 2342.75 | 211 | 3312.25 | 100536.5 | 1378.5 | 207 | 1162.5 | 1964 | 49 |
| vag_C4-like | ano_C3-C4 | cro_C3 | 6.3529216 | 1.06E-10 | 0.94811514 | 1053119.44 | 11902.8125 | 8576.5 | 2659.25 | 218 | 11726.5 | 2224.75 | 300 | 10164.5625 | 93832.0625 | 2539.125 | 485 | 1208.5 | 1641.5 | 101 |
| vag_C4-like | ano_C3-C4 | flo_C3-C4 | 4.95407463 | 3.64E-07 | 0.09954927 | 1100620 | 8011 | 6637.25 | 5016.75 | 185 | 12508.75 | 2120 | 172 | 2440.25 | 107166 | 1216.5 | 189 | 1077 | 2211.5 | 61 |
| vag_C4-like | ano_C3-C4 | chl_C3-C4 | 4.97056448 | 3.34E-07 | 0.10156373 | 1101780.5 | 8349.75 | 6760.5 | 4970 | 173.5 | 12590.75 | 2126 | 169 | 2447.5 | 107536 | 1238 | 192 | 1085 | 2225.5 | 55 |
| vag_C4-like | ano_C3-C4 | rob_C3 | 6.19129882 | 3.00E-10 | 0.94995501 | 1035270.25 | 16836.75 | 8360.5 | 2742.5 | 283.5 | 11384 | 2311.5 | 389 | 10492.75 | 91989.25 | 3201.5 | 457 | 1235 | 1590.5 | 109 |
| ang_C3-C4 | ano_C3-C4 | koc_C4 | 4.99551665 | 2.94E-07 | 0.38106565 | 1041120.69 | 10934.8125 | 7893.25 | 3421.5 | 228.5 | 8190.3125 | 2852.3125 | 217.625 | 3202.75 | 100559.25 | 1648.5 | 190 | 1391 | 1324.5 | 52 |
| ang_C3-C4 | ano_C3-C4 | bro_C4-like | 8.77250883 | 0 | 0.27799675 | 1104224.25 | 9171.5 | 7976 | 4072.5 | 187.5 | 9257.75 | 2515.5 | 165 | 3115 | 107039.5 | 1391 | 166 | 1318 | 1612.5 | 51 |
| ang_C3-C4 | ano_C3-C4 | bid_C4 | 5.48420551 | 2.08E-08 | 0.41258741 | 1145891.5 | 12308.5 | 8745.25 | 3728.5 | 216.5 | 9034.75 | 3140.5 | 242 | 3553.5 | 110949.5 | 1851 | 210 | 1511 | 1499.5 | 60 |
| ang_C3-C4 | ano_C3-C4 | aus_C4 | 5.01007627 | 2.72E-07 | 0.45748988 | 1129125 | 12144 | 8615 | 3639.25 | 203 | 8796 | 3170.25 | 225 | 3565.75 | 108344.25 | 1866 | 205.5 | 1470.5 | 1445.5 | 57 |
| ang_C3-C4 | ano_C3-C4 | tri_C4 | 5.2707167 | 6.81E-08 | 0.45675972 | 1118223 | 12343.25 | 8639.25 | 3670.5 | 202.5 | 8835 | 3174.25 | 220 | 3591.5 | 108225.75 | 1853.5 | 204.5 | 1504.5 | 1461.5 | 49 |
| koc_C4 | ano_C3-C4 | pri_C3 | 7.03972627 | 9.69E-13 | 0.88497923 | 1046194.25 | 9696.5 | 8158.75 | 3007.75 | 193 | 11327 | 2509.25 | 243 | 6344.75 | 97599.25 | 1829 | 339.5 | 1320.5 | 1576.5 | 73 |
| koc_C4 | ano_C3-C4 | son_C3-C4 | 6.13185552 | 4.36E-10 | 0.76692708 | 1027797.13 | 9543.125 | 7906.75 | 3121.75 | 168 | 11009.625 | 2696.625 | 219.75 | 4095.5 | 98848.75 | 1564.5 | 265.5 | 1313.5 | 1652.5 | 52 |
| koc_C4 | ano_C3-C4 | bro_C4-like | 13.5378381 | 0 | 0.36480972 | 1038837.5 | 8647.5 | 7096 | 4006.75 | 169 | 11552 | 2333.5 | 206 | 3294.5 | 100484.75 | 1384.5 | 223.5 | 1160 | 1886.5 | 46 |
| koc_C4 | ano_C3-C4 | cro_C3 | 6.09398672 | 5.53E-10 | 0.94886039 | 1035141.88 | 11593.375 | 8389.5 | 2618.75 | 206 | 11492.5 | 2204.5 | 285 | 9890.625 | 92364.125 | 2484.75 | 498.5 | 1186 | 1501.5 | 97 |
| koc_C4 | ano_C3-C4 | flo_C3-C4 | 5.70718663 | 5.76E-09 | 0.11140806 | 1087010 | 8037.25 | 6554.5 | 4972.5 | 168 | 12359.5 | 2063.25 | 166 | 2428 | 105703.5 | 1187.5 | 197.5 | 1055.5 | 2101 | 61 |
| koc_C4 | ano_C3-C4 | chl_C3-C4 | 6.3205008 | 1.31E-10 | 0.12371922 | 1089768.75 | 8318 | 6660.75 | 4932.75 | 168.5 | 12435 | 2067.75 | 173 | 2472.25 | 106208.25 | 1216 | 202.5 | 1061.5 | 2105 | 58 |

| | | | | | | | | | | | | | | | | | | | | |
|---|---|---|---|---|---|---|---|---|---|---|---|---|---|---|---|---|---|---|---|---|
| koc_C4 | ano_C3-C4 | rob_C3 | 6.55011048 | 2.89E-11 | 0.94729502 | 1025054.5 | 16681.5 | 8242 | 2730.5 | 292 | 11259.5 | 2277 | 403.5 | 10428 | 91019 | 3179.5 | 491.5 | 1212.5 | 1486 | 88 |
| pri_C3 | ano_C3-C4 | son_C3-C4 | 6.67437936 | 1.25E-11 | 0.13737459 | 1097462.5 | 10179 | 9490.25 | 5851.25 | 269.5 | 10109.25 | 2777.5 | 236 | 3267 | 102735.75 | 1474 | 218 | 1519 | 1829 | 66 |
| pri_C3 | ano_C3-C4 | bro_C4-like | 4.95134026 | 3.69E-07 | 0.06843345 | 1101006.5 | 9204.25 | 8275.75 | 7004.25 | 258 | 10584.75 | 2355.5 | 212 | 2697 | 103453.5 | 1346 | 170 | 1284 | 2125.5 | 54 |
| pri_C3 | ano_C3-C4 | bid_C4 | 7.19135685 | 3.23E-13 | 0.11401135 | 1148036.25 | 12643.75 | 9009 | 6976.5 | 326 | 10672.75 | 2801.5 | 257 | 3338.75 | 107162 | 1811.5 | 223 | 1422 | 2043 | 77 |
| pri_C3 | ano_C3-C4 | pal_C4-like | 5.03443955 | 2.40E-07 | 0.08629166 | 1083639.44 | 11656.8125 | 8507.0625 | 6584.8125 | 319.125 | 10056 | 2693.5 | 246.5 | 3061 | 101165.75 | 1638 | 207 | 1340.5 | 1993.5 | 62 |
| pri_C3 | ano_C3-C4 | aus_C4 | 7.28993118 | 1.56E-13 | 0.12042784 | 1131555.75 | 12558.5 | 8835 | 6766.25 | 329 | 10377.75 | 2798.5 | 255 | 3341.75 | 104861.5 | 1805.5 | 223 | 1397 | 1988.5 | 69 |
| pri_C3 | ano_C3-C4 | tri_C4 | 7.40835284 | 6.44E-14 | 0.12231487 | 1120587.5 | 12715 | 8855.25 | 6789.25 | 325.5 | 10410.75 | 2815.75 | 255.5 | 3369.5 | 104642 | 1811 | 219 | 1416 | 2013 | 62 |
| son_C3-C4 | ano_C3-C4 | bid_C4 | 8.32056014 | 0 | 0.28058633 | 1145108.75 | 12577.25 | 8852 | 4504.25 | 259 | 10597.25 | 2946 | 227 | 3553.75 | 110472.75 | 1885.5 | 188 | 1447 | 1763.5 | 60 |
| son_C3-C4 | ano_C3-C4 | pal_C4-like | 6.59343962 | 2.16E-11 | 0.23101571 | 1062687.31 | 11371.4375 | 8180.3125 | 4299.5625 | 258.125 | 9982.125 | 2754.625 | 218.75 | 3218.75 | 102211 | 1670 | 185.5 | 1329.5 | 1678 | 56 |
| son_C3-C4 | ano_C3-C4 | aus_C4 | 9.38021609 | 0 | 0.31133466 | 1130197.94 | 12434.5625 | 8685 | 4418 | 245 | 10429.8125 | 2899.0625 | 236.125 | 3585.75 | 108000.25 | 1902.5 | 193 | 1402 | 1714 | 54 |
| son_C3-C4 | ano_C3-C4 | tri_C4 | 9.5597462 | 0 | 0.31233407 | 1128638.63 | 12656.625 | 8731.25 | 4496.25 | 242.5 | 10390.875 | 2942.125 | 253.25 | 3648 | 108921.5 | 1917 | 193 | 1435 | 1748 | 57 |
| bro_C4-like | ano_C3-C4 | bid_C4 | 14.218639 | 0 | 0.65189927 | 1129096 | 12915.25 | 7774.25 | 3493.25 | 214 | 9428 | 2460 | 217.5 | 4395 | 109493.75 | 2067 | 191 | 1249.5 | 1499.5 | 49 |
| bro_C4-like | ano_C3-C4 | ram_C3-C4 | 12.8572941 | 0 | 0.63765267 | 1102985.19 | 9740.5625 | 7673.5625 | 3367.8125 | 202.125 | 9241 | 2446.25 | 183 | 4068 | 106952.5 | 1730 | 197 | 1254.5 | 1418.5 | 41 |
| bro_C4-like | ano_C3-C4 | pal_C4-like | 13.609537 | 0 | 0.62656677 | 1070450.94 | 11827.3125 | 7382.3125 | 3373.8125 | 215.125 | 8929.5 | 2392.5 | 212 | 4039 | 103624 | 1876 | 180 | 1176 | 1456.5 | 49 |
| bro_C4-like | ano_C3-C4 | aus_C4 | 14.0357438 | 0 | 0.65656566 | 1113724 | 12765.5 | 7663.5 | 3445 | 207 | 9282 | 2433.5 | 238 | 4367.25 | 107191.75 | 2073.5 | 187 | 1239 | 1464 | 46 |
| bro_C4-like | ano_C3-C4 | tri_C4 | 14.4002333 | 0 | 0.65491038 | 1104744.5 | 12912.75 | 7666.75 | 3486.25 | 197.5 | 9257.25 | 2441.75 | 244.5 | 4424 | 107130.75 | 2076 | 187 | 1249 | 1490 | 44 |
| cro_C3 | ano_C3-C4 | bid_C4 | 7.60591084 | 1.42E-14 | 0.05992912 | 1140195.19 | 12977.25 | 9372.75 | 10952.0625 | 509 | 12842.3125 | 2414.75 | 326 | 2959 | 102081.063 | 1730 | 226 | 1315.5 | 2771.125 | 104 |
| cro_C3 | ano_C3-C4 | rob_C3 | 4.35815963 | 6.56E-06 | 0.7149278 | 1089897.13 | 15863.5 | 14696.5 | 4837.625 | 500.5 | 10189.625 | 4465 | 412.25 | 5399.5 | 94843.875 | 2337.25 | 349.25 | 2154.25 | 1660.25 | 117.5 |
| cro_C3 | ano_C3-C4 | pal_C4-like | 5.17416768 | 1.15E-07 | 0.04278123 | 1069401.5 | 11870.875 | 8788.625 | 10342 | 482.75 | 12114.9375 | 2315.0625 | 284.625 | 2673.8125 | 95541.9375 | 1552.125 | 214.625 | 1211.625 | 2645.375 | 90.125 |
| cro_C3 | ano_C3-C4 | aus_C4 | 7.12101794 | 5.39E-13 | 0.05852903 | 1123942.88 | 12818.25 | 9211.75 | 10639.375 | 504.5 | 12497.375 | 2435.75 | 310.5 | 2945.75 | 99761.125 | 1750.5 | 225 | 1279 | 2678.25 | 99 |
| cro_C3 | ano_C3-C4 | tri_C4 | 7.22171247 | 2.59E-13 | 0.05923844 | 1115566.13 | 12985.5 | 9249 | 10764.625 | 493.5 | 12540.875 | 2478.75 | 304.5 | 3000.5 | 99727.875 | 1749.5 | 224.5 | 1306 | 2731.75 | 98 |
| bid_C4 | ano_C3-C4 | flo_C3-C4 | 5.43940759 | 2.68E-08 | 0.09946886 | 1212337.63 | 8906.625 | 7372.8125 | 5580.5625 | 193.125 | 14076.875 | 2274.375 | 186.75 | 2639.5625 | 118344.063 | 1335.125 | 195.125 | 1180.125 | 2371.125 | 57.125 |
| bid_C4 | ano_C3-C4 | chl_C3-C4 | 5.74930319 | 4.49E-09 | 0.10547833 | 1214274 | 9259.5 | 7486 | 5569 | 186.5 | 14151 | 2287 | 189 | 2674 | 118728.5 | 1370 | 198 | 1186 | 2390.5 | 51 |
| bid_C4 | ano_C3-C4 | rob_C3 | 6.21398609 | 2.59E-10 | 0.95157757 | 1142445.69 | 18594.8125 | 9305.5 | 3068 | 311 | 12806.5625 | 2609.3125 | 430.625 | 11623.25 | 101887.75 | 3556 | 496.5 | 1356 | 1694 | 113 |
| flo_C3-C4 | ano_C3-C4 | ram_C3-C4 | 5.93156133 | 1.51E-09 | 0.88582583 | 1188353.44 | 10793.3125 | 7243.8125 | 2568.0625 | 176.625 | 8801.5 | 2178.75 | 153.5 | 5199.25 | 115952.75 | 2033.5 | 192 | 1164 | 1279.5 | 55 |
| flo_C3-C4 | ano_C3-C4 | pal_C4-like | 4.79601096 | 8.10E-07 | 0.90238174 | 1126328.19 | 12762.3125 | 6833.0625 | 2498.3125 | 184.125 | 8319.75 | 2184 | 172 | 5089.5 | 109848.25 | 2125 | 172 | 1094 | 1254.5 | 53 |
| flo_C3-C4 | ano_C3-C4 | aus_C4 | 5.41227113 | 3.12E-08 | 0.9014885 | 1204797 | 14084.25 | 7296.75 | 2645.75 | 181 | 8830 | 2281.75 | 183 | 5612.75 | 116792.25 | 2429 | 194.5 | 1147.5 | 1325.5 | 52 |
| flo_C3-C4 | ano_C3-C4 | tri_C4 | 4.93810556 | 3.95E-07 | 0.9101331 | 1187638.38 | 14157.375 | 7245.0625 | 2607.8125 | 178.625 | 8698.875 | 2276.125 | 179.75 | 5635.3125 | 115864.063 | 2388.125 | 194.625 | 1161.625 | 1329.125 | 52.125 |

| | | | | | | | | | | | | | | | | | | | | |
|---|---|---|---|---|---|---|---|---|---|---|---|---|---|---|---|---|---|---|---|---|
| chl_C3-C4 | ano_C3-C4 | ram_C3-C4 | 5.98264019 | 1.10E-09 | 0.88251168 | 1188658.94 | 10879.3125 | 7347.0625 | 2600.3125 | 192.625 | 9192.25 | 2205.75 | 141.5 | 5169.5 | 116227.75 | 2012.5 | 196.5 | 1190 | 1289 | 45 |
| chl_C3-C4 | ano_C3-C4 | pal_C4-like | 5.06989727 | 1.99E-07 | 0.89534313 | 1127088.44 | 12787.5625 | 6943.3125 | 2535.5625 | 196.125 | 8585.75 | 2202.25 | 179.5 | 5053.75 | 110130.25 | 2141 | 170.5 | 1099 | 1276 | 53 |
| chl_C3-C4 | ano_C3-C4 | aus_C4 | 5.2564195 | 7.36E-08 | 0.90284068 | 1206073 | 14145.25 | 7424.25 | 2645.75 | 196 | 9164.75 | 2291.75 | 183 | 5581.25 | 117128.5 | 2432 | 187.5 | 1160.5 | 1352.5 | 50 |
| chl_C3-C4 | ano_C3-C4 | tri_C4 | 4.90792386 | 4.61E-07 | 0.90973076 | 1188792.75 | 14228.5 | 7367.5 | 2617 | 190.5 | 9021.75 | 2286.75 | 181 | 5615 | 116208.25 | 2405.5 | 185.5 | 1169.5 | 1354.5 | 50 |
| rob_C3 | ano_C3-C4 | aus_C4 | 6.06776269 | 6.51E-10 | 0.0488691 | 1128417 | 12748.25 | 9184.5 | 11313.25 | 523.5 | 18209.5 | 2608.5 | 394 | 3055.75 | 99767.25 | 1707.5 | 318 | 1288 | 3484 | 107 |
| rob_C3 | ano_C3-C4 | tri_C4 | 6.19208431 | 2.98E-10 | 0.04973434 | 1127364.5 | 12968.5 | 9261.5 | 11469 | 514.5 | 18300.75 | 2660.75 | 411.5 | 3121.75 | 100588.25 | 1716.5 | 327 | 1330 | 3502.5 | 111 |
| vag_C4-like | bid_C4 | pal_C4-like | 9.28173204 | 0 | 0.63898356 | 1056013.56 | 4618.625 | 4040.6875 | 1542.625 | 35.25 | 4265.75 | 1092.4375 | 32.375 | 1889.25 | 114642.938 | 796.875 | 48 | 620.875 | 798.375 | 13.375 |
| vag_C4-like | bid_C4 | aus_C4 | 5.52721295 | 1.63E-08 | 0.67372948 | 1080415.5 | 4750.6875 | 4207.5 | 1486.6875 | 32.375 | 4257.4375 | 1225.875 | 35.75 | 1764.4375 | 116047.875 | 824.25 | 35.875 | 676.25 | 743.25 | 14.25 |
| vag_C4-like | bid_C4 | tri_C4 | 6.20941665 | 2.67E-10 | 0.65539859 | 1080510.88 | 4900.8125 | 4206.875 | 1537.5625 | 34.125 | 4267.3125 | 1238.75 | 37.5 | 1807.0625 | 117421 | 820.5 | 39.125 | 703.5 | 754.5 | 17.5 |
| koc_C4 | bid_C4 | pal_C4-like | 8.88015047 | 0 | 0.43664706 | 1042382.25 | 4455.5 | 4216.1875 | 1752.6875 | 31.875 | 4611.375 | 1154.125 | 32.25 | 1618.0625 | 112747.563 | 734.625 | 34.875 | 660.375 | 808.875 | 12.375 |
| koc_C4 | bid_C4 | aus_C4 | 10.0717173 | 0 | 0.4125211 | 1075699.75 | 4816.4375 | 4277.5 | 1861.4375 | 56.375 | 4777.9375 | 1121.875 | 52.25 | 1641.1875 | 115174.875 | 796.75 | 33.875 | 678.75 | 813.75 | 11.25 |
| koc_C4 | bid_C4 | tri_C4 | 10.2486369 | 0 | 0.40301291 | 1071233.56 | 5012.1875 | 4280.375 | 1906.625 | 50.75 | 4768.6875 | 1126.4375 | 51.875 | 1653.125 | 116064.625 | 827.25 | 36.25 | 693.25 | 816.75 | 13.25 |
| pal_C4-like | bid_C4 | aus_C4 | 8.1377606 | 2.22E-16 | 0.46146965 | 1105778.06 | 4887.3125 | 4390.1875 | 1802.3125 | 47.125 | 4749.9375 | 1275.5625 | 30.125 | 1726.9375 | 118807.563 | 842.125 | 35.875 | 690.125 | 773.625 | 9.125 |
| pal_C4-like | bid_C4 | tri_C4 | 8.53329159 | 0 | 0.45749158 | 1102008.19 | 5114.75 | 4369.8125 | 1848.25 | 48.5 | 4765.375 | 1284.3125 | 33.125 | 1759.875 | 119804.063 | 872.125 | 36.75 | 697.625 | 785.125 | 12.125 |
| vag_C4-like | bro_C4-like | son_C3-C4 | 9.108525 | 0 | 0.07826831 | 989401.938 | 4754.3125 | 4514.0625 | 7584.8125 | 111.625 | 12758.25 | 1461 | 114 | 1981 | 96649 | 784.5 | 149.5 | 712 | 2552 | 29 |
| vag_C4-like | bro_C4-like | cro_C3 | 6.72339263 | 8.93E-12 | 0.93386393 | 1013846.44 | 11166.0625 | 9434.75 | 2888.75 | 249 | 12026.5 | 2412.5 | 293 | 9137.3125 | 90321.5625 | 2406.625 | 464 | 1276 | 1684.5 | 93 |
| vag_C4-like | bro_C4-like | rob_C3 | 7.72877921 | 5.55E-15 | 0.92513769 | 983783 | 15620.75 | 9004.75 | 2969.5 | 320.5 | 11618.25 | 2419 | 362 | 9222 | 87488.75 | 2947.5 | 446.5 | 1251 | 1635.5 | 108 |
| ang_C3-C4 | bro_C4-like | son_C3-C4 | 7.98740232 | 6.66E-16 | 0.06862902 | 1048290.94 | 5125.8125 | 4995.5625 | 7679.8125 | 99.625 | 10347.5 | 1488 | 97 | 1944.25 | 102934 | 806.5 | 121 | 795 | 2155 | 28 |
| koc_C4 | bro_C4-like | son_C3-C4 | 7.93771546 | 1.11E-15 | 0.06846534 | 992548.313 | 4699.9375 | 4565.8125 | 7652.5625 | 92.625 | 12839.375 | 1487.375 | 111.75 | 1940.5 | 97200.25 | 774.5 | 142.5 | 733 | 2485.5 | 30 |
| koc_C4 | bro_C4-like | cro_C3 | 6.63467488 | 1.63E-11 | 0.93368893 | 1005538.63 | 11023.375 | 9405.75 | 2860.25 | 243 | 11990 | 2392.5 | 279 | 8978.625 | 89893.125 | 2367.25 | 452 | 1278 | 1590.5 | 90 |
| koc_C4 | bro_C4-like | rob_C3 | 8.24512074 | 1.11E-16 | 0.92200353 | 984451.75 | 15706 | 9032.5 | 2974.25 | 328.5 | 11701.25 | 2388.75 | 379.5 | 9310 | 87644.5 | 2957.5 | 446.5 | 1273 | 1553 | 100 |
| son_C3-C4 | bro_C4-like | bid_C4 | 9.03029615 | 0 | 0.92533962 | 1077118.44 | 14240 | 5024.3125 | 2142.75 | 144 | 5187.0625 | 1604.5 | 113 | 8275.5625 | 105536.75 | 2713.5 | 104.625 | 782 | 879.5 | 29 |
| son_C3-C4 | bro_C4-like | flo_C3-C4 | 48.3247731 | 0 | 0.58265765 | 1085226.69 | 8852 | 2284.0625 | 4887.5 | 74.5 | 6361.5625 | 477 | 43 | 6634.5625 | 107565.5 | 1824.5 | 56.625 | 282 | 1425.5 | 18 |
| son_C3-C4 | bro_C4-like | chl_C3-C4 | 48.5109631 | 0 | 0.58342184 | 1086023.88 | 9121.8125 | 2250.125 | 4971.8125 | 67.125 | 6385.8125 | 510 | 38 | 6758.8125 | 107841.5 | 1861 | 49.625 | 308 | 1419.5 | 14 |
| son_C3-C4 | bro_C4-like | ano_C3-C4 | 28.0008013 | 0 | 0.77379515 | 1084780.44 | 9871 | 4271.0625 | 2912.5 | 117.5 | 5749.5625 | 1121 | 73 | 7249.3125 | 107124.5 | 2152 | 80.625 | 621 | 1066.5 | 20 |
| son_C3-C4 | bro_C4-like | ram_C3-C4 | 8.36869206 | 0 | 0.92951731 | 1063149.38 | 11254.375 | 5102.5625 | 1984.3125 | 110.625 | 5102.875 | 1502.625 | 105.25 | 7855.0625 | 104462.313 | 2373.625 | 107.625 | 776.125 | 830.125 | 29.125 |
| son_C3-C4 | bro_C4-like | pub_C3-C4 | 46.0853158 | 0 | 0.58721764 | 1075656.38 | 8445.5625 | 2484.9375 | 4741.125 | 57.75 | 6240.375 | 619.3125 | 42.625 | 6482.9375 | 107312.625 | 1836.25 | 49.875 | 348.25 | 1381.75 | 14.25 |
| son_C3-C4 | bro_C4-like | pal_C4-like | 10.2301296 | 0 | 0.91319574 | 1020944.31 | 13098.875 | 4699.0625 | 2088.25 | 145.5 | 4994.6875 | 1493.375 | 104.75 | 7751.5625 | 99828.5 | 2533 | 108.125 | 735 | 834 | 27 |

| | | | | | | | | | | | | | | | | | | | | |
|---|---|---|---|---|---|---|---|---|---|---|---|---|---|---|---|---|---|---|---|---|
| son_C3-C4 | bro_C4-like | aus_C4 | 8.59264504 | 0 | 0.92849359 | 1052917.38 | 13872.5625 | 4887.8125 | 2067 | 151 | 5102.625 | 1562.5625 | 107.125 | 8112.5625 | 101999.25 | 2676.5 | 106.625 | 744 | 848 | 27 |
| son_C3-C4 | bro_C4-like | tri_C4 | 8.80816861 | 0 | 0.92800381 | 1056781.31 | 14227.125 | 4879.8125 | 2098 | 156 | 5037.6875 | 1577.625 | 113.25 | 8285.0625 | 103722 | 2751.5 | 103.625 | 765 | 863 | 28 |
| cro_C3 | bro_C4-like | bid_C4 | 7.31566581 | 1.29E-13 | 0.06967861 | 1087597.19 | 13300.5 | 10227 | 9754.3125 | 476.5 | 11985.8125 | 2584.5 | 311 | 3121.5 | 97328.5625 | 1776 | 278 | 1378 | 2605.125 | 95 |
| cro_C3 | bro_C4-like | rob_C3 | 6.72701056 | 8.71E-12 | 0.61871102 | 1035343.63 | 15001.75 | 14298.5 | 4719.875 | 502.75 | 9770.875 | 4146.75 | 378.75 | 5076.75 | 90492.125 | 2221 | 371.75 | 2063 | 1640.25 | 118.25 |
| cro_C3 | bro_C4-like | pal_C4-like | 5.51620476 | 1.74E-08 | 0.05469197 | 1036788.13 | 12264.5 | 9725.75 | 9302.625 | 469 | 11437.875 | 2484 | 283 | 2878.5 | 92630.375 | 1626 | 249 | 1323 | 2495.25 | 74 |
| cro_C3 | bro_C4-like | aus_C4 | 6.98320055 | 1.45E-12 | 0.06980372 | 1062163.88 | 12971.25 | 9984 | 9343.375 | 493.5 | 11580.125 | 2560.5 | 285.5 | 3069.5 | 94098.625 | 1748.5 | 252.5 | 1322.5 | 2499.25 | 92 |
| cro_C3 | bro_C4-like | tri_C4 | 7.73638673 | 5.11E-15 | 0.07624319 | 1057423.38 | 13201.25 | 9987 | 9470.625 | 506.5 | 11638.625 | 2585.75 | 294 | 3154 | 94702.625 | 1771.5 | 262.5 | 1351.5 | 2528.75 | 92 |
| bid_C4 | bro_C4-like | rob_C3 | 7.19239405 | 3.21E-13 | 0.93230781 | 1071086.13 | 17080.875 | 9897.8125 | 3224.5625 | 350.125 | 12963.875 | 2686.875 | 405.75 | 10092.3125 | 95560.5625 | 3217.625 | 471.125 | 1370.125 | 1734.125 | 116.125 |
| rob_C3 | bro_C4-like | pal_C4-like | 5.44544651 | 2.59E-08 | 0.05327046 | 1012774.69 | 11838.0625 | 9322.75 | 9568.25 | 449 | 16141.0625 | 2553.8125 | 361.625 | 2948.5 | 90209.75 | 1594.5 | 337.5 | 1297 | 3052.5 | 102 |
| rob_C3 | bro_C4-like | aus_C4 | 7.12222373 | 5.35E-13 | 0.06916672 | 1048034.75 | 12722.25 | 9680.75 | 9740 | 497.5 | 16648.75 | 2641 | 370 | 3168.5 | 92536 | 1712.5 | 343.5 | 1315 | 3127.5 | 105 |
| rob_C3 | bro_C4-like | tri_C4 | 7.89484893 | 1.44E-15 | 0.07535843 | 1051600 | 13022.25 | 9765.75 | 9912.75 | 513.5 | 16750.5 | 2673.5 | 376 | 3263.5 | 93918.25 | 1758.5 | 355.5 | 1351 | 3159 | 112 |
| vag_C4-like | chl_C3-C4 | pri_C3 | 4.81867482 | 7.24E-07 | 0.90845387 | 1057882 | 9753.5 | 9782 | 3141.5 | 194 | 11640 | 2785.5 | 252 | 6318.25 | 98350.75 | 1869.5 | 358 | 1433 | 1724 | 69 |
| vag_C4-like | chl_C3-C4 | son_C3-C4 | 5.62785768 | 9.15E-09 | 0.74490456 | 1037024.25 | 9637.25 | 9405.5 | 3219.5 | 205 | 11393.75 | 2825.25 | 246 | 3976.5 | 99504.5 | 1543.5 | 250.5 | 1412.5 | 1771 | 50 |
| vag_C4-like | chl_C3-C4 | bro_C4-like | 10.6074974 | 0 | 0.13920495 | 1042511.88 | 6736.125 | 6695.625 | 5950.875 | 159.75 | 12474.5625 | 1832.5625 | 165.125 | 2498.5625 | 101332.313 | 1014.125 | 149.125 | 932.125 | 2385.125 | 36.125 |
| vag_C4-like | chl_C3-C4 | cro_C3 | 4.90410819 | 4.70E-07 | 0.95706608 | 1054562.44 | 11901.3125 | 10092.5 | 2726.5 | 242 | 11980.25 | 2382.5 | 301 | 10050.8125 | 93611.0625 | 2526.625 | 484 | 1322 | 1667 | 86 |
| vag_C4-like | chl_C3-C4 | rob_C3 | 5.00457425 | 2.80E-07 | 0.95617028 | 1034807 | 16776 | 9815.5 | 2814.75 | 296 | 11645.75 | 2458.25 | 387 | 10235.5 | 91776.25 | 3155.5 | 466.5 | 1324.5 | 1611.5 | 100 |
| ang_C3-C4 | chl_C3-C4 | bro_C4-like | 6.32632742 | 1.26E-10 | 0.08916067 | 1108935.94 | 7114.8125 | 7385.8125 | 6208.5625 | 172.125 | 10104.5 | 2038 | 129 | 2446.25 | 107999.5 | 1058.5 | 121 | 1046 | 2012 | 39 |
| koc_C4 | chl_C3-C4 | pri_C3 | 4.97309531 | 3.30E-07 | 0.90563607 | 1048543.75 | 9681 | 9727.75 | 3134.25 | 186 | 11530.25 | 2768 | 240 | 6283 | 97376.5 | 1819 | 336 | 1448 | 1581.5 | 74 |
| koc_C4 | chl_C3-C4 | son_C3-C4 | 4.76806045 | 9.31E-07 | 0.75448718 | 1028235.13 | 9476.875 | 9386 | 3215.25 | 193 | 11285.125 | 2880.125 | 228.75 | 3910 | 98560.75 | 1504.5 | 232.5 | 1401.5 | 1695.5 | 51 |
| koc_C4 | chl_C3-C4 | bro_C4-like | 9.437715 | 0 | 0.12378046 | 1045081.13 | 6727.875 | 6753.875 | 6027.625 | 161.75 | 12591.3125 | 1845.8125 | 160.125 | 2436.5625 | 101653.313 | 1033.125 | 160.125 | 941.625 | 2295.625 | 37.125 |
| koc_C4 | chl_C3-C4 | cro_C3 | 4.82377422 | 7.05E-07 | 0.95645925 | 1037087.88 | 11578.125 | 9960 | 2704 | 235 | 11776 | 2367 | 288 | 9769.875 | 92158.375 | 2464.75 | 479 | 1287 | 1535 | 86 |
| koc_C4 | chl_C3-C4 | rob_C3 | 4.74857359 | 1.03E-06 | 0.95781784 | 1025485.5 | 16609.5 | 9728.5 | 2818.5 | 311 | 11551 | 2479.25 | 396.5 | 10182.5 | 90758.75 | 3126.5 | 475.5 | 1312 | 1496 | 94 |
| pri_C3 | chl_C3-C4 | son_C3-C4 | 4.3324795 | 7.38E-06 | 0.09818401 | 1096784.75 | 10138 | 10878.5 | 5922.25 | 294.5 | 10188.25 | 2930.25 | 243 | 3256 | 102538 | 1421 | 193 | 1597.5 | 1864 | 69 |
| pri_C3 | chl_C3-C4 | bid_C4 | 5.0719278 | 1.97E-07 | 0.09285672 | 1150969.75 | 12897.25 | 10726 | 6887 | 346 | 10683.25 | 3055 | 265 | 3447.25 | 107106 | 1828.5 | 216 | 1580 | 2023 | 74 |
| pri_C3 | chl_C3-C4 | aus_C4 | 5.01902592 | 2.60E-07 | 0.0958642 | 1136001.25 | 12815 | 10528 | 6729.75 | 370 | 10418.25 | 3068 | 252 | 3456.25 | 105063.5 | 1828.5 | 204 | 1553 | 1976.5 | 71 |
| pri_C3 | chl_C3-C4 | tri_C4 | 5.55189264 | 1.42E-08 | 0.10499509 | 1119418.25 | 12867.25 | 10413 | 6690.5 | 358 | 10377.25 | 3044.25 | 246.5 | 3472 | 104235 | 1816 | 201 | 1557 | 1990 | 63 |
| son_C3-C4 | chl_C3-C4 | bid_C4 | 6.35578175 | 1.04E-10 | 0.28567078 | 1147513.25 | 12888.25 | 10507.5 | 4323 | 247.5 | 10592 | 3150.25 | 231 | 3619.25 | 110473.5 | 1941.5 | 209 | 1559.5 | 1716.5 | 54 |
| son_C3-C4 | chl_C3-C4 | pal_C4-like | 4.27931723 | 9.38E-06 | 0.20135932 | 1064418.63 | 11613.125 | 9700.75 | 4161 | 251 | 9961.625 | 2941.875 | 233.75 | 3249.25 | 102235.5 | 1748 | 216.5 | 1429 | 1611 | 44 |

| | | | | | | | | | | | | | | | | | | | | |
|---|---|---|---|---|---|---|---|---|---|---|---|---|---|---|---|---|---|---|---|---|
| son_C3-C4 | chl_C3-C4 | aus_C4 | 7.37402045 | 8.35E-14 | 0.31809559 | 1135126.94 | 12763.0625 | 10294.25 | 4271.5 | 262 | 10434.5625 | 3103.3125 | 248.125 | 3648.25 | 108368 | 1966.5 | 215 | 1513.5 | 1661 | 48 |
| son_C3-C4 | chl_C3-C4 | tri_C4 | 7.77912298 | 3.66E-15 | 0.3268986 | 1127152.88 | 12894.875 | 10255.25 | 4295.5 | 259 | 10346.875 | 3105.625 | 254.75 | 3683.5 | 108626.25 | 1962 | 211 | 1539.5 | 1694 | 50 |
| bro_C4-like | chl_C3-C4 | bid_C4 | 10.6106865 | 0 | 0.87149234 | 1135313.13 | 14001.0625 | 7313.375 | 2622.5625 | 162.125 | 7347.375 | 1937.0625 | 164.625 | 6585.875 | 110668.063 | 2526.125 | 172.75 | 1006.625 | 1107.125 | 41.125 |
| bro_C4-like | chl_C3-C4 | ano_C3-C4 | 24.8944856 | 0 | 0.59667949 | 1151076.13 | 8087.3125 | 6583.625 | 3466.5625 | 118.625 | 7811.875 | 1599.0625 | 106.125 | 4361.875 | 114813.063 | 1603.125 | 126.75 | 878.125 | 1294.625 | 40.125 |
| bro_C4-like | chl_C3-C4 | ram_C3-C4 | 9.46676443 | 0 | 0.8756551 | 1106496.63 | 10739.3125 | 7232.875 | 2549.0625 | 139.125 | 7155.625 | 1942.5625 | 168.625 | 6213.625 | 107913.313 | 2098.625 | 175.75 | 978.625 | 1067.125 | 35.125 |
| bro_C4-like | chl_C3-C4 | pal_C4-like | 10.5784055 | 0 | 0.86051639 | 1075513.63 | 12782.8125 | 6991.375 | 2556.8125 | 162.625 | 6974.875 | 1884.5625 | 170.125 | 6031.875 | 104724.063 | 2326.125 | 176.75 | 939.125 | 1075.125 | 31.125 |
| bro_C4-like | chl_C3-C4 | aus_C4 | 10.7039497 | 0 | 0.86970638 | 1121708.44 | 13837 | 7205.5625 | 2614.5 | 159 | 7232.3125 | 1924.75 | 186 | 6528.8125 | 108594.5 | 2542.5 | 181.125 | 993.5 | 1078 | 34 |
| bro_C4-like | chl_C3-C4 | tri_C4 | 10.8907634 | 0 | 0.86818423 | 1104156.44 | 13911.75 | 7148.5625 | 2597.5 | 152 | 7199.5625 | 1899 | 188.5 | 6499.5625 | 107708.5 | 2524 | 173.125 | 994.5 | 1100 | 33 |
| cro_C3 | chl_C3-C4 | bid_C4 | 5.61784149 | 9.69E-09 | 0.04783142 | 1143725.69 | 13317.25 | 11048.5 | 10806.5625 | 519 | 12890.0625 | 2600 | 331 | 3012.25 | 102095.063 | 1756.5 | 269 | 1437 | 2770.125 | 90 |
| cro_C3 | chl_C3-C4 | aus_C4 | 5.4628965 | 2.35E-08 | 0.04811656 | 1128774.13 | 13175 | 10880.75 | 10549.875 | 524.5 | 12576.625 | 2612 | 313.5 | 3013.25 | 100016.125 | 1785 | 248.5 | 1406.5 | 2681.25 | 90 |
| cro_C3 | chl_C3-C4 | tri_C4 | 5.8588251 | 2.34E-09 | 0.05155856 | 1113850.13 | 13228 | 10756.5 | 10556.875 | 513 | 12520.875 | 2619.25 | 308 | 3050.75 | 99394.875 | 1765 | 256.5 | 1417.5 | 2703.75 | 85 |
| bid_C4 | chl_C3-C4 | rob_C3 | 4.34760532 | 6.89E-06 | 0.96292571 | 1145098.94 | 18591.0625 | 10969.25 | 3133 | 340 | 13137.8125 | 2803.5625 | 424.125 | 11360 | 101913.25 | 3504 | 513 | 1473 | 1721 | 113 |
| rob_C3 | chl_C3-C4 | aus_C4 | 4.70621248 | 1.26E-06 | 0.04095534 | 1133528 | 13075.75 | 10817 | 11135.75 | 542.5 | 18278.5 | 2793.5 | 403 | 3149.75 | 100113.75 | 1736.5 | 324 | 1428 | 3440 | 112 |
| rob_C3 | chl_C3-C4 | tri_C4 | 5.06068911 | 2.09E-07 | 0.04383663 | 1125379.5 | 13234 | 10766 | 11185.5 | 532.5 | 18216.25 | 2804.25 | 404.5 | 3188.5 | 100226.5 | 1742 | 336 | 1453 | 3438.5 | 108 |
| vag_C4-like | flo_C3-C4 | pri_C3 | 5.23120533 | 8.44E-08 | 0.90192332 | 1055919.5 | 9644.25 | 9480.5 | 3147.5 | 207 | 11570 | 2762.5 | 248 | 6303 | 97873.75 | 1872 | 338 | 1423 | 1699 | 74 |
| vag_C4-like | flo_C3-C4 | son_C3-C4 | 6.74801309 | 7.54E-12 | 0.70691531 | 1035213 | 9546 | 9038.75 | 3254.5 | 212 | 11346.5 | 2783 | 239 | 3920.25 | 99057.5 | 1550.5 | 261 | 1398 | 1749 | 55 |
| vag_C4-like | flo_C3-C4 | bro_C4-like | 10.3832956 | 0 | 0.13380683 | 1039803.75 | 6668.25 | 6415.5 | 5969.5 | 167 | 12405.5 | 1792.5 | 172 | 2437.75 | 100820.75 | 992.5 | 158 | 892 | 2359 | 43 |
| vag_C4-like | flo_C3-C4 | cro_C3 | 5.26383933 | 7.07E-08 | 0.95335923 | 1050046.19 | 11676.5625 | 9749.5 | 2735.75 | 254.5 | 11867.25 | 2367.25 | 300 | 9899.5625 | 93085.8125 | 2524.125 | 490 | 1289.5 | 1629 | 96 |
| vag_C4-like | flo_C3-C4 | rob_C3 | 5.0200796 | 2.59E-07 | 0.95570287 | 1031005 | 16599 | 9456.25 | 2814.75 | 316 | 11526.5 | 2457.25 | 388 | 10170.25 | 91059.5 | 3163.5 | 460 | 1305.5 | 1572.5 | 111 |
| ang_C3-C4 | flo_C3-C4 | bro_C4-like | 6.44458608 | 5.82E-11 | 0.08888647 | 1105806.5 | 7111.75 | 7137.75 | 6144.25 | 172.5 | 10056.75 | 1957 | 126 | 2365.5 | 107650 | 1027.5 | 140.5 | 1008 | 2016 | 38 |
| koc_C4 | flo_C3-C4 | pri_C3 | 6.08447416 | 5.87E-10 | 0.88872814 | 1045901.5 | 9580.75 | 9484.75 | 3129.25 | 197 | 11484.75 | 2687 | 235 | 6219.25 | 96928.75 | 1820.5 | 324 | 1415 | 1574.5 | 72 |
| koc_C4 | flo_C3-C4 | son_C3-C4 | 6.80369068 | 5.13E-12 | 0.69080984 | 1027010.38 | 9417.625 | 9106.5 | 3247.25 | 194 | 11263.875 | 2772.875 | 230.75 | 3832.75 | 98227 | 1517.5 | 249 | 1382 | 1675.5 | 53 |
| koc_C4 | flo_C3-C4 | bro_C4-like | 9.98395398 | 0 | 0.1269302 | 1041289 | 6687 | 6548.5 | 6014.25 | 167 | 12504.75 | 1773.75 | 171 | 2390.25 | 101057 | 1008.5 | 173 | 893.5 | 2274.5 | 41 |
| koc_C4 | flo_C3-C4 | cro_C3 | 5.57983967 | 1.21E-08 | 0.94970332 | 1032498.13 | 11370.375 | 9691.25 | 2701.25 | 245.5 | 11686.5 | 2314.5 | 283 | 9617.125 | 91659.125 | 2461.25 | 473 | 1263 | 1508 | 91 |
| koc_C4 | flo_C3-C4 | rob_C3 | 5.33822821 | 4.70E-08 | 0.95313903 | 1022392.25 | 16469.25 | 9470.5 | 2803.5 | 317 | 11477.25 | 2425.25 | 385.75 | 10118.75 | 90170.25 | 3139.75 | 455 | 1299 | 1476.25 | 102.25 |
| pri_C3 | flo_C3-C4 | son_C3-C4 | 5.58413621 | 1.18E-08 | 0.11776447 | 1096702.25 | 10115.75 | 10548.25 | 5922 | 277.5 | 10150 | 2828 | 225 | 3241 | 102246.75 | 1424 | 198 | 1585 | 1864.5 | 75 |
| pri_C3 | flo_C3-C4 | bro_C4-like | 4.17218021 | 1.51E-05 | 0.03371564 | 1102137.25 | 7154.25 | 7406.5 | 9137 | 222 | 11243.5 | 1807.25 | 163 | 2063 | 103845.75 | 971 | 124 | 985.5 | 2535 | 47 |
| pri_C3 | flo_C3-C4 | bid_C4 | 5.74360262 | 4.65E-09 | 0.10265678 | 1148688.63 | 12801.125 | 10438.125 | 6844.125 | 329.75 | 10575.0625 | 2995.8125 | 259.125 | 3436.0625 | 106682.563 | 1800.125 | 231.125 | 1550.625 | 2026.625 | 76.125 |

| | | | | | | | | | | | | | | | | | | | | |
|---|---|---|---|---|---|---|---|---|---|---|---|---|---|---|---|---|---|---|---|---|
| pri_C3 | flo_C3-C4 | aus_C4 | 5.65791273 | 7.68E-09 | 0.10571481 | 1133163.75 | 12760 | 10269 | 6690.5 | 349 | 10318 | 3017 | 252 | 3451.25 | 104598 | 1811.5 | 225 | 1527 | 1964 | 72 |
| pri_C3 | flo_C3-C4 | tri_C4 | 6.37333999 | 9.29E-11 | 0.11677387 | 1117801.13 | 12788.125 | 10145.125 | 6664.625 | 343.75 | 10276.0625 | 2983.0625 | 248.625 | 3469.8125 | 103843.563 | 1783.625 | 224.125 | 1535.625 | 1986.625 | 66.125 |
| son_C3-C4 | flo_C3-C4 | bid_C4 | 7.84716938 | 2.22E-15 | 0.32505612 | 1145931.25 | 12838 | 10112.5 | 4261.75 | 266 | 10507.75 | 3059 | 234 | 3638.25 | 110141.5 | 1903.5 | 215 | 1527 | 1725.5 | 64 |
| son_C3-C4 | flo_C3-C4 | pal_C4-like | 6.36614238 | 9.74E-11 | 0.26628665 | 1064196.13 | 11628.625 | 9394.25 | 4084.5 | 261 | 9892.375 | 2845.625 | 232.75 | 3295.25 | 102017.5 | 1707 | 215.5 | 1412.5 | 1622 | 50 |
| son_C3-C4 | flo_C3-C4 | aus_C4 | 8.68630427 | 0 | 0.35446715 | 1134012.44 | 12756.0625 | 9927.25 | 4213.5 | 273 | 10369.3125 | 3033.0625 | 240.125 | 3681.25 | 108050.5 | 1940.5 | 226 | 1490 | 1663 | 62 |
| son_C3-C4 | flo_C3-C4 | tri_C4 | 9.07701033 | 0 | 0.35919578 | 1125758.63 | 12854.875 | 9904.5 | 4245.5 | 270 | 10280.625 | 3030.375 | 252.75 | 3711.5 | 108233.25 | 1927 | 222 | 1509 | 1699 | 62 |
| bro_C4-like | flo_C3-C4 | bid_C4 | 10.7479586 | 0 | 0.87145909 | 1131409.38 | 13898.875 | 7045.875 | 2582.125 | 171.75 | 7300.5625 | 1893.8125 | 168.625 | 6560.3125 | 110037.063 | 2497.625 | 172.125 | 956.125 | 1090.625 | 51.125 |
| bro_C4-like | flo_C3-C4 | ano_C3-C4 | 25.4142081 | 0 | 0.5922962 | 1148263.5 | 7988 | 6313 | 3454.5 | 128 | 7785 | 1544 | 105 | 4319.5 | 114321.5 | 1593 | 133 | 845 | 1278 | 35 |
| bro_C4-like | flo_C3-C4 | ram_C3-C4 | 9.18883667 | 0 | 0.88048919 | 1103914.5 | 10667.5 | 6980.75 | 2455.5 | 165.5 | 7165.75 | 1876.5 | 165.5 | 6142.25 | 107358.75 | 2111.5 | 178 | 947.5 | 1039.5 | 38 |
| bro_C4-like | flo_C3-C4 | pal_C4-like | 10.7895834 | 0 | 0.86238999 | 1073849.25 | 12743.5 | 6754.5 | 2491.75 | 171 | 6933 | 1815.5 | 178 | 6053.5 | 104328.5 | 2297 | 172 | 913 | 1057.5 | 33 |
| bro_C4-like | flo_C3-C4 | aus_C4 | 10.6162806 | 0 | 0.87287499 | 1118566.25 | 13783 | 6957 | 2554.5 | 172 | 7195.25 | 1877.75 | 187 | 6524.5 | 108172.25 | 2520.5 | 185.5 | 942.5 | 1064 | 44 |
| bro_C4-like | flo_C3-C4 | tri_C4 | 11.3529115 | 0 | 0.86591795 | 1101481.63 | 13821.625 | 6903.125 | 2555.875 | 166.75 | 7155.5625 | 1835.8125 | 193.625 | 6486.0625 | 107174.313 | 2493.625 | 181.625 | 939.125 | 1083.125 | 40.125 |
| cro_C3 | flo_C3-C4 | bid_C4 | 6.08253661 | 5.94E-10 | 0.05181665 | 1138725.31 | 13208.125 | 10721.625 | 10656.6875 | 514.75 | 12676.375 | 2550.3125 | 325.125 | 2993.3125 | 101598.125 | 1724.125 | 280.625 | 1395.625 | 2761.75 | 101.125 |
| cro_C3 | flo_C3-C4 | pal_C4-like | 4.32944734 | 7.48E-06 | 0.03841623 | 1068877.38 | 12060.5 | 10077.5 | 10080.875 | 496 | 11999.125 | 2427.75 | 294.5 | 2733.5 | 95184.125 | 1540.5 | 261.5 | 1297 | 2639.75 | 82 |
| cro_C3 | flo_C3-C4 | aus_C4 | 6.20960493 | 2.67E-10 | 0.05465584 | 1124934.38 | 13109 | 10567.75 | 10410.625 | 520.5 | 12377.625 | 2566.75 | 310.5 | 3020.25 | 99583.375 | 1764 | 269.5 | 1366 | 2665.75 | 108 |
| cro_C3 | flo_C3-C4 | tri_C4 | 6.63398807 | 1.64E-11 | 0.05823566 | 1109748.75 | 13124.875 | 10452.625 | 10400 | 508.75 | 12303.9375 | 2560.8125 | 307.125 | 3045.5625 | 98955.1875 | 1727.625 | 277.625 | 1372.625 | 2691.375 | 107.125 |
| bid_C4 | flo_C3-C4 | rob_C3 | 4.53360764 | 2.90E-06 | 0.96171499 | 1141618.06 | 18459.875 | 10616.125 | 3087.8125 | 355.125 | 13049.4375 | 2747.375 | 419.5 | 11299.125 | 101329.063 | 3517.375 | 513.25 | 1435.625 | 1692.875 | 124.375 |
| chl_C3-C4 | flo_C3-C4 | pub_C3-C4 | 5.47122468 | 2.24E-08 | 0.53474734 | 1207337.06 | 4012.1875 | 4042.125 | 1650.625 | 41.25 | 4559.125 | 1353.125 | 48.25 | 1695.0625 | 128489.563 | 664.625 | 30.125 | 629.625 | 743.625 | 10.625 |
| rob_C3 | flo_C3-C4 | aus_C4 | 4.69080587 | 1.36E-06 | 0.04063381 | 1131359.25 | 13024.75 | 10482 | 11103.75 | 524.5 | 18167.5 | 2763.5 | 387.25 | 3116.75 | 99619.5 | 1728.75 | 342 | 1408 | 3443.25 | 119.25 |
| rob_C3 | flo_C3-C4 | tri_C4 | 5.42932182 | 2.84E-08 | 0.04652121 | 1123122.38 | 13150.875 | 10437.125 | 11141.875 | 520.25 | 18101.0625 | 2754.0625 | 388.875 | 3163.3125 | 99724.3125 | 1719.875 | 358.125 | 1421.625 | 3452.875 | 117.375 |
| bro_C4-like | pal_C4-like | aus_C4 | 4.56924252 | 2.45E-06 | 0.02294693 | 1051071.63 | 5350.125 | 4932.375 | 10004.125 | 170.75 | 11778.3125 | 988.0625 | 98.125 | 1199.8125 | 102533.063 | 746.125 | 91.625 | 628.125 | 2738.625 | 31.125 |
| bro_C4-like | pal_C4-like | tri_C4 | 4.38951745 | 5.68E-06 | 0.02225858 | 1045421 | 5562.25 | 4924.5 | 10053.75 | 179 | 11802.625 | 1015.875 | 103.75 | 1221.625 | 103095.875 | 776.75 | 91.75 | 648.25 | 2767.75 | 26.25 |
| bid_C4 | pal_C4-like | chl_C3-C4 | 4.82879491 | 6.88E-07 | 0.97751141 | 1124148.44 | 12793.375 | 4966.5625 | 1143.875 | 88.75 | 5368.3125 | 926.125 | 94.75 | 10391.0625 | 111457.125 | 2924.75 | 169.625 | 627.75 | 723.25 | 25.25 |
| flo_C3-C4 | pal_C4-like | aus_C4 | 5.21493888 | 9.21E-08 | 0.02781522 | 1107732.13 | 5661.125 | 5182.625 | 9853.375 | 172.75 | 12203.0625 | 1063.0625 | 95.125 | 1314.5625 | 108660.563 | 806.125 | 95.125 | 641.125 | 2768.125 | 25.125 |
| flo_C3-C4 | pal_C4-like | tri_C4 | 5.52197857 | 1.68E-08 | 0.02971838 | 1091847.13 | 5801.125 | 5118.8125 | 9746.8125 | 178.125 | 12095 | 1056 | 93 | 1322.1875 | 107900.438 | 807.375 | 91.375 | 649.375 | 2730.875 | 24.375 |
| chl_C3-C4 | pal_C4-like | aus_C4 | 5.5936717 | 1.11E-08 | 0.02925984 | 1109658.88 | 5703.125 | 5192.125 | 9931.375 | 175.75 | 12456.0625 | 1040.0625 | 103.125 | 1308.0625 | 109049.813 | 810.125 | 98.125 | 649.125 | 2811.125 | 23.125 |
| chl_C3-C4 | pal_C4-like | tri_C4 | 6.19704863 | 2.89E-10 | 0.03240317 | 1093492.25 | 5839.25 | 5116.25 | 9869 | 179 | 12329.125 | 1022.625 | 97.25 | 1318.875 | 108295.625 | 815.75 | 97.25 | 655.25 | 2780.25 | 20.25 |
| ano_C3-C4 | pal_C4-like | aus_C4 | 5.01771349 | 2.62E-07 | 0.02766738 | 1102712.81 | 5620.125 | 5177.1875 | 9453.375 | 162.75 | 10735.375 | 1043.0625 | 74.125 | 1282.375 | 108713.563 | 817.125 | 95.75 | 642.125 | 2624.125 | 19.125 |

| | | | | | | | | | | | | | | | | | | | | |
|---|---|---|---|---|---|---|---|---|---|---|---|---|---|---|---|---|---|---|---|---|
| ano_C3-C4 | pal_C4-like | tri_C4 | 4.87678337 | 5.40E-07 | 0.02700865 | 1092662.44 | 5811 | 5137.3125 | 9458 | 169 | 10780.6875 | 1052.875 | 76.75 | 1286.1875 | 108579.375 | 832.75 | 98.875 | 647.25 | 2608.25 | 18.25 |
| vag_C4-like | pri_C3 | cro_C3 | 24.2645206 | 0 | 0.66468165 | 1033210.69 | 9697.8125 | 7957.75 | 4566.25 | 215 | 14963.75 | 2556.75 | 301 | 6540.0625 | 91911.8125 | 1843.125 | 360 | 1137.5 | 2284.5 | 95 |
| vag_C4-like | pri_C3 | rob_C3 | 4.30994118 | 8.17E-06 | 0.92474989 | 1002437.5 | 15323 | 8541.5 | 3626.5 | 348.75 | 13638.5 | 3280.5 | 472.25 | 7532.5 | 88226 | 2644 | 408.25 | 1321 | 1905.5 | 122.25 |
| ang_C3-C4 | pri_C3 | cro_C3 | 23.062522 | 0 | 0.86574486 | 1091405.63 | 11483.125 | 4100.75 | 3042.75 | 111 | 7769.5 | 1513.5 | 146 | 11374.875 | 99478.625 | 2702.25 | 138.5 | 592 | 1228.5 | 43 |
| koc_C4 | pri_C3 | cro_C3 | 24.2251769 | 0 | 0.65528197 | 1019607.88 | 9460.625 | 7818.25 | 4539.75 | 209 | 14788.5 | 2532.5 | 284 | 6348.125 | 90941.625 | 1821.25 | 353 | 1084 | 2149.5 | 88 |
| koc_C4 | pri_C3 | rob_C3 | 4.41117742 | 5.14E-06 | 0.9228963 | 994594.5 | 15239.5 | 8490.5 | 3613.25 | 323.75 | 13563 | 3260.25 | 465.75 | 7485.5 | 87499.5 | 2658 | 399.25 | 1276 | 1803 | 105.25 |
| son_C3-C4 | pri_C3 | cro_C3 | 17.5633383 | 0 | 0.70650063 | 1057208.88 | 9743.375 | 8716.75 | 4159.5 | 226 | 12753 | 2780.75 | 249 | 6099.625 | 95420.375 | 1832.25 | 323 | 1245.5 | 1862 | 71 |
| bro_C4-like | pri_C3 | cro_C3 | 20.1972934 | 0 | 0.69910065 | 1072753.63 | 9971.875 | 8547.25 | 4338.5 | 220 | 13111.5 | 2720 | 266 | 6480.375 | 96222.125 | 1906.75 | 358 | 1219 | 2032 | 73 |
| cro_C3 | pri_C3 | bid_C4 | 25.8427806 | 0 | 0.34750283 | 1109594.69 | 16365.25 | 8580.5 | 6940.8125 | 385 | 10374.5625 | 2728.25 | 318 | 4971.75 | 99087.5625 | 2439.5 | 240 | 1203.5 | 1988.625 | 92 |
| cro_C3 | pri_C3 | flo_C3-C4 | 21.6019708 | 0 | 0.30020258 | 1111782.88 | 14001 | 8567.5 | 6831.375 | 373 | 10335.875 | 2729.25 | 271 | 4489 | 99039.875 | 2087 | 232 | 1196.5 | 1969.75 | 84 |
| cro_C3 | pri_C3 | chl_C3-C4 | 22.3139659 | 0 | 0.30637996 | 1114232.88 | 14345.5 | 8668.5 | 6894.375 | 355 | 10449.625 | 2737.25 | 280 | 4573.5 | 99426.125 | 2091 | 235 | 1209.5 | 1967.75 | 85 |
| cro_C3 | pri_C3 | ano_C3-C4 | 21.5097448 | 0 | 0.29198213 | 1114530.38 | 12854.5 | 8598.25 | 6917.375 | 385 | 10461.375 | 2697.5 | 240 | 4437.75 | 99591.625 | 2007.5 | 226 | 1225 | 1988.75 | 79 |
| cro_C3 | pri_C3 | ram_C3-C4 | 21.8705408 | 0 | 0.30146414 | 1078314.88 | 13005.5 | 8339.75 | 6718.125 | 341 | 10144.375 | 2633 | 258 | 4396 | 96523.125 | 2001 | 212 | 1166 | 1960.25 | 78 |
| cro_C3 | pri_C3 | pub_C3-C4 | 19.6891034 | 0 | 0.285798 | 1098843.38 | 13394.25 | 8593.5 | 6764.375 | 349 | 10210.375 | 2803.5 | 272.5 | 4388.5 | 98273.375 | 2062 | 234 | 1198.5 | 1977.75 | 78 |
| cro_C3 | pri_C3 | pal_C4-like | 24.7750218 | 0 | 0.33617633 | 1055076.63 | 15258.5 | 8152.75 | 6685.625 | 379.5 | 9956.625 | 2589.75 | 300 | 4664 | 94193.375 | 2259 | 218 | 1158.5 | 1917.75 | 73 |
| cro_C3 | pri_C3 | aus_C4 | 25.2913807 | 0 | 0.3449957 | 1083260.63 | 15907.75 | 8274.75 | 6753.125 | 390.5 | 10026.625 | 2655.5 | 318.5 | 4813.75 | 95733.125 | 2356.5 | 226 | 1147.5 | 1921.75 | 81 |
| cro_C3 | pri_C3 | tri_C4 | 25.6175884 | 0 | 0.34614404 | 1080665.13 | 16196.75 | 8358.75 | 6855.625 | 386.5 | 10115.125 | 2689.5 | 317.5 | 4895 | 96523.875 | 2392 | 228 | 1181.5 | 1950.75 | 82 |
| bid_C4 | pri_C3 | rob_C3 | 4.07693707 | 2.28E-05 | 0.92983914 | 1091968.94 | 16744.5625 | 9418.25 | 3947.75 | 367.75 | 15146.0625 | 3604.8125 | 480.875 | 8149.75 | 96453.25 | 2872.5 | 441.25 | 1404 | 2053 | 134.25 |
| rob_C3 | pri_C3 | aus_C4 | 4.11826189 | 1.91E-05 | 0.07087268 | 1066708.75 | 14819.25 | 9115.5 | 7978 | 438.75 | 16281.25 | 3501 | 469.25 | 3842.5 | 93130.75 | 1998.5 | 359.75 | 1346 | 2790.5 | 116.25 |
| rob_C3 | pri_C3 | tri_C4 | 4.38168095 | 5.89E-06 | 0.07460786 | 1071406.25 | 15180.5 | 9267.5 | 8102 | 441.5 | 16413.75 | 3559.25 | 467.5 | 3925.5 | 94708.75 | 2056.5 | 374.5 | 1397 | 2832.5 | 119 |
| vag_C4-like | pub_C3-C4 | pri_C3 | 5.1601818 | 1.24E-07 | 0.90581978 | 1047484.75 | 9693.25 | 9043.25 | 3078.5 | 199 | 11398 | 2702.25 | 248 | 6321 | 97699.5 | 1908 | 349 | 1348.5 | 1689 | 73 |
| vag_C4-like | pub_C3-C4 | son_C3-C4 | 8.39165444 | 0 | 0.69138582 | 1016548.5 | 9358.5 | 8456.25 | 3224.25 | 220.5 | 11090.625 | 2641.625 | 221.25 | 3946.875 | 97844.875 | 1557.75 | 255.25 | 1302.25 | 1737.25 | 59.25 |
| vag_C4-like | pub_C3-C4 | bro_C4-like | 12.876327 | 0 | 0.16523572 | 1040508 | 6876 | 6174.5 | 5801.5 | 160 | 12344.5 | 1742.25 | 161 | 2545.75 | 101619.5 | 1078.5 | 176 | 871.5 | 2337 | 42 |
| vag_C4-like | pub_C3-C4 | cro_C3 | 7.73173791 | 5.33E-15 | 0.93527686 | 1042708.94 | 11671.8125 | 9222.5 | 2745.75 | 236.5 | 11819.25 | 2215 | 297 | 9884.5625 | 93084.5625 | 2575.125 | 480 | 1261 | 1612 | 88 |
| vag_C4-like | pub_C3-C4 | rob_C3 | 5.73846477 | 4.79E-09 | 0.95100352 | 1013313.5 | 16519.25 | 8949.25 | 2729.75 | 298 | 11316.5 | 2330 | 375 | 10089 | 90210.25 | 3157 | 444 | 1227 | 1562.5 | 108 |
| ang_C3-C4 | pub_C3-C4 | son_C3-C4 | 4.47352711 | 3.85E-06 | 0.74821745 | 1076679.19 | 9836.8125 | 9377.3125 | 3158.3125 | 214.625 | 8759.625 | 2847.125 | 178.25 | 3771.875 | 104270.125 | 1572.75 | 187.25 | 1472.25 | 1376.25 | 58.25 |
| ang_C3-C4 | pub_C3-C4 | bro_C4-like | 8.45034642 | 0 | 0.11489865 | 1105586.5 | 7269.5 | 6843.25 | 6028.25 | 177.5 | 10013.5 | 1912.75 | 134 | 2447 | 108355.25 | 1095 | 130 | 980.5 | 1996 | 47 |
| ang_C3-C4 | pub_C3-C4 | tri_C4 | 4.21814567 | 1.23E-05 | 0.53060517 | 1101629.31 | 12329.1875 | 9277.6875 | 3566.4375 | 210.875 | 8705.0625 | 3228.0625 | 217.125 | 3610.5625 | 106884.063 | 1853.125 | 188.125 | 1499.125 | 1439.125 | 61.125 |

| | | | | | | | | | | | | | | | | | | | | |
|---|---|---|---|---|---|---|---|---|---|---|---|---|---|---|---|---|---|---|---|---|
| koc_C4 | pub_C3-C4 | pri_C3 | 5.31635793 | 5.30E-08 | 0.90340375 | 1036894.25 | 9584.75 | 9008.5 | 3052.25 | 188 | 11338.25 | 2667 | 237 | 6270 | 96777.5 | 1839.5 | 322.5 | 1378 | 1575.5 | 69 |
| koc_C4 | pub_C3-C4 | son_C3-C4 | 8.20338024 | 1.11E-16 | 0.67348033 | 1007307.81 | 9182.1875 | 8432.3125 | 3235.5625 | 202.875 | 11058 | 2664.5 | 209.5 | 3842.375 | 97027.375 | 1505 | 233.25 | 1317 | 1673.75 | 55.5 |
| koc_C4 | pub_C3-C4 | bro_C4-like | 12.1632073 | 0 | 0.1546735 | 1039640.75 | 6854 | 6234.25 | 5866.25 | 162 | 12470.25 | 1740 | 159 | 2495 | 101641 | 1071.5 | 170.5 | 892 | 2268.5 | 38 |
| koc_C4 | pub_C3-C4 | cro_C3 | 7.61172103 | 1.37E-14 | 0.93463702 | 1026035.13 | 11391.625 | 9136.5 | 2703.25 | 228.5 | 11668 | 2184.75 | 277 | 9598.875 | 91706.125 | 2495.25 | 459.5 | 1261.5 | 1510 | 88 |
| koc_C4 | pub_C3-C4 | chl_C3-C4 | 4.11461073 | 1.94E-05 | 0.02337143 | 1066156.5 | 4610.5 | 3899.5 | 8459.25 | 125.5 | 13606.75 | 895.75 | 94 | 1076.75 | 105237.5 | 620 | 98.75 | 532.75 | 2735.25 | 26.25 |
| koc_C4 | pub_C3-C4 | rob_C3 | 6.06907788 | 6.46E-10 | 0.94811578 | 1002629.44 | 16377.0625 | 8887.8125 | 2731.0625 | 295.375 | 11281.5 | 2309.5 | 386 | 10013 | 89235.5 | 3103.25 | 443 | 1235.25 | 1462 | 98.25 |
| pri_C3 | pub_C3-C4 | son_C3-C4 | 5.34848068 | 4.44E-08 | 0.11041405 | 1073016.94 | 9707.5625 | 9901.5625 | 5890.3125 | 294.125 | 10033.125 | 2744.125 | 229.25 | 3134.625 | 100549.625 | 1403.25 | 185.25 | 1503.25 | 1858.75 | 71.25 |
| pri_C3 | pub_C3-C4 | bid_C4 | 5.69097014 | 6.33E-09 | 0.09918021 | 1129207.81 | 12580.4375 | 9797.6875 | 6828.6875 | 334.875 | 10525.0625 | 2914.0625 | 259.125 | 3345.0625 | 105560.063 | 1792.125 | 214.125 | 1454.125 | 2045.625 | 69.125 |
| pri_C3 | pub_C3-C4 | aus_C4 | 6.0028253 | 9.73E-10 | 0.10816019 | 1099931 | 12358.5 | 9565.75 | 6551.5 | 338 | 10064 | 2849.25 | 249 | 3298.25 | 101617.75 | 1737.5 | 207 | 1408.5 | 1951 | 70 |
| pri_C3 | pub_C3-C4 | tri_C4 | 6.03147913 | 8.15E-10 | 0.1084305 | 1098135.81 | 12585.6875 | 9521.6875 | 6630.4375 | 335.875 | 10173.3125 | 2893.3125 | 247.625 | 3347.8125 | 102673.313 | 1774.625 | 206.125 | 1433.125 | 1992.125 | 66.125 |
| son_C3-C4 | pub_C3-C4 | bid_C4 | 9.90906802 | 0 | 0.35127266 | 1113808.63 | 12542.1875 | 9267.125 | 4224.9375 | 248.375 | 10176.5625 | 2874.125 | 212.25 | 3605.5625 | 107736.625 | 1874.75 | 220.625 | 1419.25 | 1709.75 | 63.25 |
| son_C3-C4 | pub_C3-C4 | chl_C3-C4 | 6.12882822 | 4.44E-10 | 0.03047051 | 1123985.13 | 5063.4375 | 4162.625 | 9494.1875 | 125.375 | 12342.5625 | 865.375 | 69.25 | 1136.5625 | 110923.875 | 667.75 | 89.875 | 586 | 2698.5 | 26.5 |
| son_C3-C4 | pub_C3-C4 | ram_C3-C4 | 4.99212463 | 2.99E-07 | 0.23935763 | 1095367.25 | 9511.3125 | 9294.6875 | 3995.5 | 226 | 9927.6875 | 2876.5 | 207.5 | 3228.625 | 106070.688 | 1564.875 | 238.75 | 1444.875 | 1625.375 | 44.375 |
| son_C3-C4 | pub_C3-C4 | pal_C4-like | 8.72647907 | 0 | 0.31549215 | 1047230.94 | 11446.6875 | 8676 | 4074.625 | 250.25 | 9624.5625 | 2734.1875 | 217.875 | 3352 | 100956.375 | 1698.25 | 222 | 1344.75 | 1613.25 | 54.25 |
| son_C3-C4 | pub_C3-C4 | aus_C4 | 10.8763292 | 0 | 0.37664046 | 1084915.63 | 12230.6875 | 9027.0625 | 4087.125 | 245.25 | 9894.125 | 2754.1875 | 219.375 | 3559.5625 | 103460.875 | 1853.75 | 220.625 | 1348.25 | 1627.25 | 61.25 |
| son_C3-C4 | pub_C3-C4 | tri_C4 | 10.9920213 | 0 | 0.37387323 | 1090893.06 | 12552.3125 | 9055.8125 | 4191.6875 | 246.375 | 9945.125 | 2815.5 | 233 | 3637.25 | 105563.375 | 1881.25 | 219.5 | 1401.25 | 1681.25 | 65.25 |
| bro_C4-like | pub_C3-C4 | bid_C4 | 13.3808534 | 0 | 0.84186314 | 1125685.81 | 13826.9375 | 6651.6875 | 2690.6875 | 173.875 | 7460.3125 | 1831.3125 | 165.625 | 6406.3125 | 110371.813 | 2473.625 | 176.125 | 940.125 | 1168.625 | 47.125 |
| bro_C4-like | pub_C3-C4 | ano_C3-C4 | 26.2859875 | 0 | 0.56415175 | 1128420.75 | 7832 | 5867 | 3517.5 | 119 | 7807.75 | 1472.5 | 118 | 4119.5 | 112862 | 1500 | 125 | 794 | 1354 | 42 |
| bro_C4-like | pub_C3-C4 | ram_C3-C4 | 10.6371702 | 0 | 0.86240158 | 1095122.44 | 10638.8125 | 6557.0625 | 2509.0625 | 163.625 | 7230.25 | 1839.5 | 157 | 6036 | 107401.75 | 2088 | 173 | 943 | 1095.5 | 39 |
| bro_C4-like | pub_C3-C4 | pal_C4-like | 12.3483836 | 0 | 0.8431452 | 1074608.75 | 12814.5 | 6447.25 | 2548 | 178.5 | 7108 | 1774 | 175 | 5934.5 | 105216 | 2293 | 172 | 903 | 1119.5 | 35 |
| bro_C4-like | pub_C3-C4 | aus_C4 | 13.6032895 | 0 | 0.84002053 | 1098248.5 | 13512.5 | 6555 | 2601.5 | 170 | 7249 | 1744.25 | 179 | 6245.5 | 106492.25 | 2425.5 | 176.5 | 902.5 | 1120 | 42 |
| bro_C4-like | pub_C3-C4 | tri_C4 | 13.8174206 | 0 | 0.83796093 | 1093300.81 | 13760.9375 | 6512.9375 | 2638.4375 | 162.875 | 7301.3125 | 1762.3125 | 187.625 | 6293.0625 | 107245.063 | 2451.625 | 171.625 | 925.125 | 1155.125 | 42.125 |
| cro_C3 | pub_C3-C4 | bid_C4 | 8.27228539 | 0 | 0.06709513 | 1120497.75 | 13095.1875 | 9982.9375 | 10601.25 | 494.875 | 12562.625 | 2390.8125 | 314.125 | 2981.3125 | 100565.875 | 1703.125 | 256.625 | 1359.625 | 2786.75 | 90.125 |
| cro_C3 | pub_C3-C4 | chl_C3-C4 | 4.58975693 | 2.22E-06 | 0.01196722 | 1133488.56 | 5111.3125 | 4309.5625 | 16616.1875 | 226.625 | 14420.625 | 785 | 100 | 976.75 | 103464.125 | 576 | 100.75 | 546.25 | 3782 | 36.25 |
| cro_C3 | pub_C3-C4 | rob_C3 | 4.44017289 | 4.50E-06 | 0.72313878 | 1065895.56 | 15629.0625 | 14864.0625 | 4691.1875 | 528.125 | 9953.875 | 4316.75 | 409.5 | 5294.75 | 92957.375 | 2286.5 | 401 | 2124 | 1672.25 | 114 |
| cro_C3 | pub_C3-C4 | pal_C4-like | 6.58407824 | 2.30E-11 | 0.05573178 | 1062185.63 | 12028 | 9461 | 10092.375 | 474 | 11937.625 | 2315.5 | 292 | 2774.5 | 95122.625 | 1527.5 | 238.5 | 1305 | 2660.75 | 77 |
| cro_C3 | pub_C3-C4 | aus_C4 | 8.52064017 | 0 | 0.07155253 | 1092816.63 | 12774.75 | 9761.25 | 10170.625 | 486.5 | 12065.625 | 2346.25 | 302.5 | 2949.25 | 96883.375 | 1687 | 242.5 | 1301 | 2641.75 | 92 |
| cro_C3 | pub_C3-C4 | tri_C4 | 8.94768558 | 0 | 0.07416004 | 1091153.69 | 13032.4375 | 9706.6875 | 10357.8125 | 480.875 | 12190.1875 | 2380.3125 | 304.125 | 3019.3125 | 97930.1875 | 1699.625 | 256.125 | 1335.625 | 2713.875 | 92.125 |

| | | | | | | | | | | | | | | | | | | | | |
|---|---|---|---|---|---|---|---|---|---|---|---|---|---|---|---|---|---|---|---|---|
| bid_C4 | pub_C3-C4 | chl_C3-C4 | 4.2213234 | 1.22E-05 | 0.0225337 | 1180691.75 | 5168.375 | 4321.25 | 9373.875 | 133.75 | 15350.1875 | 974.8125 | 99.125 | 1168.4375 | 117004.563 | 717.125 | 105.125 | 567.875 | 3057.375 | 28.375 |
| bid_C4 | pub_C3-C4 | rob_C3 | 5.22905303 | 8.54E-08 | 0.95679251 | 1110372.19 | 18114.6875 | 9878.25 | 2988.375 | 326.25 | 12736.25 | 2604.625 | 405.75 | 11102.4375 | 99297.8125 | 3453.625 | 479.375 | 1340.625 | 1660.625 | 119.125 |
| flo_C3-C4 | pub_C3-C4 | aus_C4 | 4.39567401 | 5.53E-06 | 0.97798128 | 1159382.13 | 15191.8125 | 4094.625 | 1072.3125 | 90.125 | 4585.375 | 879.8125 | 98.125 | 9429.875 | 113820.563 | 3064.125 | 119.75 | 519.625 | 623.625 | 32.125 |
| chl_C3-C4 | pub_C3-C4 | ano_C3-C4 | 4.64454545 | 1.71E-06 | 0.96032524 | 1193493.19 | 8842 | 4353.3125 | 1214 | 60.75 | 5163.0625 | 999 | 75 | 6203.0625 | 121842.25 | 1913.25 | 107.125 | 581.25 | 738.5 | 22.25 |
| chl_C3-C4 | pub_C3-C4 | aus_C4 | 4.42193585 | 4.90E-06 | 0.97657897 | 1162855.44 | 15223.75 | 4292.5625 | 1134.25 | 106.25 | 5062.3125 | 935 | 106 | 9243.0625 | 114311 | 3051.25 | 142.125 | 547.25 | 686.5 | 29.25 |
| chl_C3-C4 | pub_C3-C4 | tri_C4 | 4.42099426 | 4.92E-06 | 0.97673888 | 1146901 | 15279.1875 | 4192.5 | 1114.1875 | 103.125 | 5014.375 | 916.8125 | 103.125 | 9204.625 | 113704.563 | 3039.375 | 135.25 | 554.875 | 686.625 | 27.375 |
| rob_C3 | pub_C3-C4 | aus_C4 | 5.96755914 | 1.21E-09 | 0.05032266 | 1082188.44 | 12487.25 | 9628.3125 | 10681.5 | 490.5 | 17505.0625 | 2523 | 383 | 2955.3125 | 95479.5 | 1632.5 | 330.125 | 1259 | 3305.5 | 116 |
| rob_C3 | pub_C3-C4 | tri_C4 | 6.20830313 | 2.69E-10 | 0.05158072 | 1088035.56 | 12787.9375 | 9666.9375 | 10944.4375 | 487.375 | 17697.5625 | 2578.3125 | 389.125 | 3033.3125 | 97307.3125 | 1670.625 | 340.125 | 1308.625 | 3372.625 | 117.125 |
| vag_C4-like | ram_C3-C4 | ang_C3-C4 | 5.19491173 | 1.03E-07 | 0.85604113 | 1020310.25 | 8452.25 | 7115 | 2751 | 156.5 | 9689.5 | 2401 | 191.5 | 4482.25 | 99120.25 | 1562.5 | 182 | 1138.5 | 1491.5 | 39 |
| vag_C4-like | ram_C3-C4 | pri_C3 | 4.57230569 | 2.41E-06 | 0.94449964 | 1029157.25 | 9966.25 | 7384.25 | 2525 | 181.5 | 9926.25 | 2217.75 | 187 | 7446.5 | 96543.25 | 2115.5 | 291.5 | 1100.5 | 1427.5 | 55 |
| vag_C4-like | ram_C3-C4 | son_C3-C4 | 4.60373126 | 2.08E-06 | 0.90729871 | 1021192.56 | 10014.9375 | 7166.1875 | 2633.4375 | 167.375 | 9763.5 | 2322.5 | 197 | 5365.75 | 98855.75 | 1838.5 | 211 | 1142 | 1442.5 | 52 |
| vag_C4-like | ram_C3-C4 | cro_C3 | 5.21742166 | 9.09E-08 | 0.96584771 | 1025279.94 | 11972.3125 | 7600.5 | 2233.75 | 213.5 | 10152.25 | 1903 | 251 | 11256.8125 | 91868.3125 | 2747.125 | 377.5 | 1045 | 1356 | 84 |
| ang_C3-C4 | ram_C3-C4 | koc_C4 | 5.55806808 | 1.37E-08 | 0.15619173 | 1009219.88 | 9639.125 | 7122.3125 | 4381.8125 | 183.125 | 8372.5625 | 2373.8125 | 199.625 | 2745.5 | 98011.75 | 1410.5 | 142.5 | 1157 | 1474.5 | 43 |
| ang_C3-C4 | ram_C3-C4 | bid_C4 | 5.74266835 | 4.67E-09 | 0.15232055 | 1112252.94 | 10797.8125 | 7775.8125 | 4837.5625 | 198.625 | 9269 | 2599 | 207 | 3001.25 | 108470 | 1618 | 164.5 | 1257 | 1671.5 | 41 |
| ang_C3-C4 | ram_C3-C4 | pal_C4-like | 4.21538991 | 1.25E-05 | 0.11835656 | 1049452.69 | 9889.5625 | 7326.3125 | 4607.5625 | 193.625 | 8729.625 | 2460.375 | 209.75 | 2748.625 | 102214.125 | 1444.25 | 140.25 | 1215.25 | 1584.75 | 40.25 |
| ang_C3-C4 | ram_C3-C4 | aus_C4 | 6.22382869 | 2.44E-10 | 0.17100538 | 1085740.06 | 10551.9375 | 7581.9375 | 4654.4375 | 194.375 | 8966.125 | 2562.625 | 193.25 | 2994.125 | 104545.375 | 1588.75 | 150.75 | 1221.25 | 1589.75 | 39.25 |
| ang_C3-C4 | ram_C3-C4 | tri_C4 | 6.24899283 | 2.07E-10 | 0.16700523 | 1087105.69 | 10815.3125 | 7569.0625 | 4774.8125 | 206.125 | 9098.375 | 2596.375 | 194.75 | 3033.125 | 106118.625 | 1611.75 | 154.75 | 1235.25 | 1642.75 | 35.25 |
| koc_C4 | ram_C3-C4 | pri_C3 | 4.7967379 | 8.07E-07 | 0.94244637 | 1015946.94 | 9857.25 | 7361.8125 | 2481.5 | 164.5 | 9832.8125 | 2162.75 | 191 | 7382.3125 | 95307.5 | 2020.5 | 285.125 | 1110.5 | 1334.5 | 59 |
| koc_C4 | ram_C3-C4 | son_C3-C4 | 4.59303016 | 2.19E-06 | 0.90400512 | 1009539.13 | 9853.125 | 7201.0625 | 2623.5625 | 144.625 | 9715.125 | 2314.375 | 200.75 | 5226.0625 | 97765.0625 | 1787.125 | 208.125 | 1146.125 | 1387.625 | 47.125 |
| koc_C4 | ram_C3-C4 | cro_C3 | 5.60484558 | 1.04E-08 | 0.96232276 | 1004645.56 | 11597.125 | 7493.5625 | 2226.25 | 201.5 | 10008.5625 | 1873 | 246 | 10895.4375 | 90116.125 | 2654.25 | 378.625 | 1031 | 1265 | 90 |
| pri_C3 | ram_C3-C4 | son_C3-C4 | 4.540425 | 2.81E-06 | 0.11029308 | 1077118.69 | 9922.0625 | 9876.5625 | 5719.0625 | 295.625 | 9883.75 | 2962.25 | 244 | 3304 | 101019 | 1457 | 210 | 1536.5 | 1780.5 | 66 |
| pri_C3 | ram_C3-C4 | bid_C4 | 5.64743375 | 8.17E-09 | 0.06333886 | 1118580.44 | 10969.3125 | 8007.0625 | 8192.5625 | 308.125 | 10943 | 2369.75 | 204 | 2763.5 | 105213.75 | 1554.5 | 191.5 | 1206.5 | 2256 | 59 |
| pri_C3 | ram_C3-C4 | pal_C4-like | 4.29855443 | 8.60E-06 | 0.05029602 | 1056439.44 | 10137.8125 | 7570.5625 | 7763.3125 | 282.125 | 10342.125 | 2259.125 | 200.25 | 2550.625 | 99383.875 | 1429.75 | 174.75 | 1163.25 | 2151.75 | 50.25 |
| pri_C3 | ram_C3-C4 | aus_C4 | 5.92191889 | 1.60E-09 | 0.06923915 | 1091276.31 | 10818.9375 | 7837.4375 | 7823.4375 | 313.375 | 10531.125 | 2325.375 | 194.25 | 2734.375 | 101513.625 | 1521.25 | 172.75 | 1165.75 | 2143.75 | 59.25 |
| pri_C3 | ram_C3-C4 | tri_C4 | 5.64209422 | 8.42E-09 | 0.06553284 | 1094010.94 | 11010.8125 | 7825.8125 | 7994.3125 | 313.625 | 10732.375 | 2379.625 | 199.25 | 2773.375 | 103038.125 | 1555.25 | 177.75 | 1193.25 | 2227.25 | 54.25 |
| son_C3-C4 | ram_C3-C4 | bid_C4 | 5.46943714 | 2.26E-08 | 0.10229192 | 1129208.44 | 11070.75 | 7970.5625 | 5919.75 | 224.5 | 11048.25 | 2531.6875 | 208.375 | 2917.75 | 109912.938 | 1616.875 | 183 | 1273.375 | 2015.375 | 52.375 |
| son_C3-C4 | ram_C3-C4 | aus_C4 | 6.87167056 | 3.19E-12 | 0.12651152 | 1103852.44 | 10860.8125 | 7759.5 | 5726 | 227.5 | 10796.5 | 2434.75 | 215.5 | 2911.4375 | 105931.688 | 1610.375 | 174.875 | 1215.375 | 1931.875 | 46.375 |
| son_C3-C4 | ram_C3-C4 | tri_C4 | 6.84221526 | 3.92E-12 | 0.12352373 | 1112146.88 | 11142.9375 | 7780.5 | 5890.5625 | 235.125 | 10904.1875 | 2486 | 227.5 | 2965.8125 | 108372.375 | 1642.75 | 174.125 | 1246.75 | 2004.25 | 49.25 |

| | | | | | | | | | | | | | | | | | | | | |
|---|---|---|---|---|---|---|---|---|---|---|---|---|---|---|---|---|---|---|---|---|
| bro_C4-like | ram_C3-C4 | tri_C4 | 4.33177385 | 7.40E-06 | 0.1021401 | 1076771.19 | 10716.0625 | 7587.5625 | 5268.5625 | 214.125 | 10507.375 | 2580.875 | 236.25 | 2886.625 | 104530.625 | 1568.25 | 176.25 | 1231.75 | 1885.25 | 47.25 |
| cro_C3 | ram_C3-C4 | bid_C4 | 7.12374628 | 5.29E-13 | 0.04409921 | 1111424.63 | 11317.3125 | 8225.8125 | 12222.625 | 411.625 | 12983.0625 | 2024 | 274 | 2494.5 | 100254.813 | 1485.5 | 219.5 | 1141 | 2984.625 | 91 |
| cro_C3 | ram_C3-C4 | rob_C3 | 4.35591672 | 6.63E-06 | 0.68116722 | 1067156.63 | 15573.25 | 14845.5 | 4815.125 | 505.75 | 10039.125 | 4445 | 393.75 | 5235.75 | 93209.875 | 2299 | 384.75 | 2156.5 | 1677.75 | 122.25 |
| cro_C3 | ram_C3-C4 | pal_C4-like | 4.48880604 | 3.58E-06 | 0.02924241 | 1042559.75 | 10289.625 | 7767.375 | 11492.5 | 398.25 | 12231.8125 | 1956.6875 | 251.875 | 2243.9375 | 93866.5625 | 1326.875 | 201.375 | 1096.875 | 2814.125 | 78.375 |
| cro_C3 | ram_C3-C4 | aus_C4 | 6.26076353 | 1.92E-10 | 0.04047309 | 1084897.19 | 11014.4375 | 8065.4375 | 11732.0625 | 434.375 | 12546.25 | 2011.875 | 261.75 | 2421.875 | 96631.25 | 1456.25 | 215.75 | 1088.25 | 2841 | 85.25 |
| cro_C3 | ram_C3-C4 | tri_C4 | 6.92531728 | 2.19E-12 | 0.04388335 | 1087904.56 | 11292.8125 | 8051.8125 | 12007.4375 | 421.625 | 12764.25 | 2034.125 | 269.75 | 2491.875 | 98207.25 | 1486.25 | 220.75 | 1119.75 | 2930.5 | 86.25 |
| ano_C3-C4 | ram_C3-C4 | aus_C4 | 4.59647494 | 2.15E-06 | 0.15631876 | 1156251.75 | 11091.6875 | 8061.25 | 4552.9375 | 188.875 | 9344.1875 | 2779.625 | 196.25 | 3108.1875 | 111755.875 | 1650.25 | 176.375 | 1344.25 | 1762.25 | 41.25 |
| ano_C3-C4 | ram_C3-C4 | tri_C4 | 4.22019765 | 1.22E-05 | 0.14250263 | 1151950.63 | 11211.5625 | 7982.125 | 4662.0625 | 197.125 | 9448.4375 | 2826.375 | 199.75 | 3131.4375 | 112406.625 | 1642.75 | 178.375 | 1354.25 | 1812.25 | 40.25 |
| rob_C3 | ram_C3-C4 | aus_C4 | 4.59084583 | 2.21E-06 | 0.03000793 | 1099121.56 | 11039.9375 | 8129.9375 | 12417.9375 | 419.875 | 18178.875 | 2251.875 | 319.25 | 2566.375 | 97599.125 | 1472.75 | 279.25 | 1106.75 | 3619.25 | 97.25 |
| rob_C3 | ram_C3-C4 | tri_C4 | 4.67304438 | 1.49E-06 | 0.02995864 | 1105808.94 | 11303.5625 | 8127.5625 | 12737.5625 | 415.625 | 18452.875 | 2287.125 | 328.25 | 2609.875 | 99529.375 | 1499.75 | 287.25 | 1148.25 | 3709.75 | 97.25 |
| vag_C4-like | son_C3-C4 | cro_C3 | 9.39108458 | 0 | 0.90250399 | 1000691.94 | 10826.8125 | 9331 | 3058 | 255.5 | 12391.5 | 2390 | 288 | 8573.5625 | 89536.0625 | 2246.125 | 423 | 1234 | 1763.5 | 96 |
| vag_C4-like | son_C3-C4 | rob_C3 | 9.06559781 | 0 | 0.9074808 | 997973 | 15623.25 | 9164 | 3172.25 | 353 | 12191 | 2512.75 | 401 | 8981.5 | 89174.25 | 2908 | 414 | 1257.5 | 1711.5 | 115 |
| koc_C4 | son_C3-C4 | cro_C3 | 9.6876877 | 0 | 0.89809843 | 985896.25 | 10601.625 | 9164.125 | 3025.5 | 231.5 | 12204.625 | 2342.25 | 276 | 8364 | 88396.125 | 2202.25 | 414.75 | 1197.5 | 1654.5 | 97 |
| koc_C4 | son_C3-C4 | rob_C3 | 9.63217461 | 0 | 0.90336352 | 989744 | 15538.875 | 9000 | 3181.375 | 348.25 | 12081.375 | 2482.5 | 394.5 | 9015.625 | 88355.25 | 2897 | 427.25 | 1231.5 | 1620.5 | 115 |
| pri_C3 | son_C3-C4 | rob_C3 | 4.70780815 | 1.25E-06 | 0.9176316 | 1054409.88 | 16026.875 | 11716.875 | 3628.375 | 439 | 9604.25 | 3252.25 | 345.75 | 7442.5 | 94085.75 | 2740 | 366.25 | 1604 | 1464 | 95.25 |
| bro_C4-like | son_C3-C4 | cro_C3 | 4.59191622 | 2.20E-06 | 0.98132237 | 1041908.06 | 12622.125 | 5124.3125 | 1459.25 | 125.5 | 5656.8125 | 1223.25 | 149.5 | 13622.6875 | 95646.125 | 3166.75 | 220.625 | 656 | 827 | 36 |
| cro_C3 | son_C3-C4 | bid_C4 | 10.6764286 | 0 | 0.10508862 | 1086638.19 | 13812.75 | 10185.25 | 9261.8125 | 444 | 11785.5625 | 2549.25 | 317 | 3337.5 | 97917.5625 | 1909.5 | 265.5 | 1338.5 | 2438.625 | 99 |
| cro_C3 | son_C3-C4 | flo_C3-C4 | 4.503906 | 3.34E-06 | 0.04543926 | 1091144.13 | 11258 | 10169.5 | 9227.625 | 420.25 | 11850.625 | 2484.25 | 254 | 2805.25 | 98114.375 | 1549.75 | 258 | 1298.75 | 2476.25 | 93.25 |
| cro_C3 | son_C3-C4 | chl_C3-C4 | 5.57209929 | 1.26E-08 | 0.05539475 | 1093332.63 | 11653.5 | 10240 | 9308.875 | 420 | 11996.875 | 2496.5 | 256.5 | 2896 | 98404.875 | 1568.5 | 261.5 | 1300.5 | 2479.75 | 85 |
| cro_C3 | son_C3-C4 | rob_C3 | 8.63551758 | 0 | 0.6036915 | 1053476 | 15256.625 | 14016.375 | 4837 | 502.75 | 9876.125 | 4085.5 | 376.25 | 5230.25 | 92778.625 | 2325.75 | 363.25 | 1953.25 | 1623.75 | 117.5 |
| cro_C3 | son_C3-C4 | ram_C3-C4 | 4.2420945 | 1.11E-05 | 0.04489893 | 1072368.38 | 10702.25 | 10099.75 | 9074.875 | 393.5 | 11679.125 | 2560.25 | 249.5 | 2866.5 | 96939.625 | 1580.5 | 266.5 | 1293 | 2412.25 | 92 |
| cro_C3 | son_C3-C4 | pal_C4-like | 8.81583277 | 0 | 0.09304539 | 1019420.75 | 12658.875 | 9630.125 | 8670.25 | 427.75 | 11055.625 | 2475.75 | 291 | 3111.25 | 91564.875 | 1731 | 254.5 | 1265.5 | 2302.75 | 82 |
| cro_C3 | son_C3-C4 | aus_C4 | 10.6168126 | 0 | 0.10890144 | 1060489.31 | 13398.3125 | 9935.0625 | 8873.1875 | 464.625 | 11342.625 | 2513.25 | 295.5 | 3290.5 | 94333.875 | 1855.5 | 246.5 | 1287 | 2334.75 | 95 |
| cro_C3 | son_C3-C4 | tri_C4 | 10.872083 | 0 | 0.11023542 | 1066255.25 | 13754.375 | 9969.625 | 9106.5 | 482.75 | 11544.125 | 2586.75 | 307 | 3394.5 | 96109.375 | 1910.5 | 258.5 | 1315.5 | 2392.25 | 101 |
| bid_C4 | son_C3-C4 | rob_C3 | 9.45191077 | 0 | 0.90883408 | 1103629.31 | 17373.9375 | 10112.875 | 3511.375 | 385.25 | 13764.0625 | 2787.3125 | 431.125 | 10005.5 | 99082.25 | 3209.5 | 449.5 | 1401.5 | 1870.5 | 128 |
| flo_C3-C4 | son_C3-C4 | rob_C3 | 4.71869245 | 1.19E-06 | 0.95512852 | 1110402.63 | 17636.375 | 10090.875 | 2997.875 | 392.5 | 11268 | 2650.25 | 338 | 10049.75 | 99338.5 | 3246.75 | 420 | 1336.75 | 1571.5 | 116.25 |
| chl_C3-C4 | son_C3-C4 | rob_C3 | 5.12006851 | 1.53E-07 | 0.95078941 | 1110850.38 | 17706.375 | 10199.125 | 3064.125 | 388.25 | 11689.25 | 2684.25 | 333 | 10023.75 | 99663.5 | 3252.5 | 428 | 1355.5 | 1584 | 95 |
| rob_C3 | son_C3-C4 | ram_C3-C4 | 5.14245578 | 1.36E-07 | 0.05073515 | 1095607.5 | 10788.875 | 10058 | 9853.875 | 398.25 | 17430.1875 | 2699.5625 | 329.125 | 3081.9375 | 98786.8125 | 1575.625 | 380.375 | 1361.125 | 3198.625 | 109.125 |

| | | | | | | | | | | | | | | | | | | | | |
|---|---|---|---|---|---|---|---|---|---|---|---|---|---|---|---|---|---|---|---|---|
| rob_C3 | son_C3-C4 | pal_C4-like | 7.86107451 | 2.00E-15 | 0.08114967 | 1023283.81 | 12486.625 | 9460.6875 | 9258.875 | 415.75 | 16106.5625 | 2651.25 | 385.5 | 3234.8125 | 91634.5 | 1723 | 362.625 | 1288 | 2981 | 112 |
| rob_C3 | son_C3-C4 | aus_C4 | 9.55408986 | 0 | 0.09514441 | 1080827.56 | 13440.3125 | 9903.1875 | 9637.5625 | 482.625 | 16867.375 | 2739 | 406 | 3464.375 | 95768.75 | 1844.5 | 376.25 | 1328 | 3094.5 | 119 |
| rob_C3 | son_C3-C4 | tri_C4 | 9.3782509 | 0 | 0.09247264 | 1090323.56 | 13824.375 | 9964.9375 | 9906.375 | 503.75 | 17141.3125 | 2822.5 | 415 | 3544.3125 | 98046.25 | 1892.5 | 391.125 | 1367 | 3154 | 130 |
| koc_C4 | vag_C4-like | bid_C4 | 11.0363814 | 0 | 0.70980569 | 1030906.69 | 4590.375 | 3231.25 | 1299.8125 | 45.125 | 3672.25 | 816.3125 | 32.625 | 1998.9375 | 112365.25 | 855.5 | 34.875 | 515.5 | 568 | 13.5 |
| koc_C4 | vag_C4-like | aus_C4 | 9.56525707 | 0 | 0.80378986 | 999709.438 | 5025.875 | 3154.5625 | 1155.875 | 38.75 | 3505.8125 | 757.875 | 43.25 | 2388.3125 | 107305.125 | 950.25 | 32.625 | 491.75 | 536.25 | 9.25 |
| koc_C4 | vag_C4-like | tri_C4 | 9.85086591 | 0 | 0.79187123 | 998778.313 | 5138.3125 | 3180.8125 | 1210.6875 | 39.875 | 3546.6875 | 792.0625 | 44.625 | 2384.8125 | 108406.688 | 958.375 | 35.625 | 494.875 | 547.875 | 10.375 |

**Table S3.** HyDe significant hybridization tests for all species of *Flaveria* excluding *F. pringlei* ($C_3$).

| P1 | Hybrid | P2 | Zscore | Pvalue | Gamma | AAAA | AAAB | AABA | AABB | AABC | ABAA | ABAB | ABAC | ABBA | BAAA | ABBC | CABC | BACA | BCAA | ABCD |
|---|---|---|---|---|---|---|---|---|---|---|---|---|---|---|---|---|---|---|---|---|
| vag_C4-like | ang_C3-C4 | son_C3-C4 | 4.21720008 | 1.24E-05 | 0.75901639 | 995854.75 | 9264.75 | 7812.75 | 3149 | 176 | 10981 | 2855 | 214 | 3781 | 95804.75 | 1509.5 | 237.5 | 1224 | 1713 | 68 |
| vag_C4-like | ang_C3-C4 | cro_C3 | 8.21280974 | 1.11E-16 | 0.9274543 | 1026894.94 | 11389.0625 | 8426.25 | 2836.75 | 216 | 11950.75 | 2266.5 | 282 | 9556.8125 | 91860.8125 | 2431.125 | 419.5 | 1189 | 1687.5 | 87 |
| vag_C4-like | ang_C3-C4 | rob_C3 | 6.94178987 | 1.95E-12 | 0.93796621 | 992896.5 | 16035.5 | 8111 | 2827.5 | 326.5 | 11348 | 2341 | 374 | 9697 | 88368.5 | 3040.5 | 419.5 | 1137 | 1596.5 | 110 |
| koc_C4 | ang_C3-C4 | son_C3-C4 | 4.6196644 | 1.92E-06 | 0.73230656 | 988616.313 | 9154.375 | 7790.0625 | 3146.75 | 175 | 10916.9375 | 2827.375 | 204.75 | 3701.0625 | 95207.75 | 1495.5 | 233.625 | 1187 | 1627.5 | 60 |
| koc_C4 | ang_C3-C4 | cro_C3 | 8.2091839 | 1.11E-16 | 0.92597499 | 1012797.31 | 11146.875 | 8349.5625 | 2799.25 | 208 | 11796.8125 | 2233.5 | 253 | 9310.4375 | 90740.875 | 2404.25 | 418.625 | 1129 | 1557.5 | 83 |
| koc_C4 | ang_C3-C4 | rob_C3 | 7.34363236 | 1.05E-13 | 0.93510788 | 986353.188 | 15976 | 8053.3125 | 2812.25 | 311.5 | 11289.8125 | 2300.75 | 362.5 | 9671.5625 | 87768 | 3049.5 | 419.625 | 1111 | 1489 | 92 |
| son_C3-C4 | ang_C3-C4 | bid_C4 | 5.29507184 | 5.96E-08 | 0.28063096 | 1086819.25 | 12276 | 8571.5 | 4055.25 | 244 | 10080.75 | 3074.75 | 204.5 | 3457.25 | 104907.75 | 1850.5 | 186 | 1306 | 1652.5 | 71 |
| son_C3-C4 | ang_C3-C4 | aus_C4 | 6.49024782 | 4.31E-11 | 0.30777842 | 1060536.44 | 11952.0625 | 8270.5 | 3991 | 242 | 9862.0625 | 2947.5625 | 214.125 | 3411.5 | 101080.75 | 1816 | 186 | 1223 | 1580 | 67 |
| son_C3-C4 | ang_C3-C4 | tri_C4 | 6.59609589 | 2.12E-11 | 0.31435399 | 1063385.88 | 12289.625 | 8374.5 | 4054 | 241 | 9864.375 | 3013.875 | 226.75 | 3490.75 | 102737.75 | 1844.5 | 184 | 1276 | 1638 | 72 |
| bro_C4 _-ike | ang_C3-C4 | bid_C4 | 4.09419512 | 2.12E-05 | 0.39649578 | 1099188 | 12309.25 | 8622.75 | 3732 | 235 | 9940 | 3267 | 225.5 | 3572.5 | 105763.5 | 1862 | 212 | 1377.5 | 1596 | 75 |
| bro_C4 -ike | ang_C3-C4 | aus_C4 | 4.644361 | 1.71E-06 | 0.38559322 | 1074325.25 | 11993 | 8386.75 | 3704 | 223 | 9703.25 | 3160.25 | 241 | 3501.5 | 102302.5 | 1828 | 194 | 1317.5 | 1547 | 64 |
| bro_C4 -ike | ang_C3-C4 | tri_C4 | 4.97281764 | 3.30E-07 | 0.42259414 | 1068437.75 | 12232.5 | 8403.75 | 3718 | 219 | 9703.5 | 3200.5 | 252.5 | 3579.25 | 102860.25 | 1837.5 | 195 | 1341.5 | 1590 | 66 |
| cro_C3 | ang_C3-C4 | bid_C4 | 9.02138585 | 0 | 0.07679948 | 1102867.19 | 13131.75 | 9135 | 10207.3125 | 450 | 12267.0625 | 2414.75 | 298.5 | 3063 | 99119.8125 | 1781 | 228 | 1259 | 2626.625 | 86 |
| cro_C3 | ang_C3-C4 | rob_C3 | 5.16559135 | 1.20E-07 | 0.69780843 | 1048165.13 | 15326.5 | 13709.75 | 4670.125 | 494.75 | 9781.375 | 4237.5 | 383.25 | 5236.5 | 91674.875 | 2281 | 346.25 | 1979 | 1607.75 | 103.25 |
| cro_C3 | ang_C3-C4 | pal_C4-like | 7.3066077 | 1.38E-13 | 0.06499929 | 1047456.13 | 12128 | 8612.5 | 9751.375 | 436 | 11683.875 | 2354 | 273 | 2868.25 | 93871.625 | 1619.5 | 213 | 1207 | 2526.75 | 67 |
| cro_C3 | ang_C3-C4 | aus_C4 | 8.81849196 | 0 | 0.07721735 | 1076612.38 | 12736.75 | 8839.25 | 9887.625 | 434.5 | 11829.875 | 2379.75 | 281.5 | 3008 | 95635.625 | 1750 | 215 | 1193 | 2518.75 | 86 |
| cro_C3 | ang_C3-C4 | tri_C4 | 9.42926082 | 0 | 0.0819511 | 1072816.63 | 13040.5 | 8889 | 9997.625 | 433 | 11930.375 | 2408 | 290 | 3085.5 | 96458.125 | 1766 | 216 | 1217 | 2572.25 | 83 |
| bid_C4 | ang_C3-C4 | rob_C3 | 6.60007401 | 2.07E-11 | 0.94318365 | 1084336.94 | 17598.5625 | 8940.75 | 3067.5 | 339.5 | 12651.0625 | 2583.0625 | 390.125 | 10625 | 96868 | 3328 | 456 | 1235 | 1710.5 | 111 |
| rob_C3 | ang_C3-C4 | pal_C4-like | 4.82841354 | 6.89E-07 | 0.0431795 | 1021572.44 | 11564.0625 | 8323.25 | 10103.25 | 431 | 16562.8125 | 2470.8125 | 367.625 | 2815.25 | 91032.5 | 1556 | 331.5 | 1172 | 3149.5 | 93 |
| rob_C3 | ang_C3-C4 | aus_C4 | 7.13488409 | 4.88E-13 | 0.06208366 | 1059362 | 12371.75 | 8636 | 10321.25 | 453.5 | 17071 | 2507 | 359 | 3024.25 | 93445.75 | 1687 | 338.5 | 1155 | 3212 | 105 |
| rob_C3 | ang_C3-C4 | tri_C4 | 6.87647553 | 3.09E-12 | 0.05990988 | 1062773 | 12704.75 | 8754 | 10513 | 454.5 | 17230.75 | 2584.75 | 366.5 | 3090 | 94960.25 | 1716.5 | 344.5 | 1206 | 3279.5 | 104 |
| vag_C4-like | ano_C3-C4 | ang_C3-C4 | 4.89616627 | 4.89E-07 | 0.61034965 | 1052360.5 | 8262 | 7975.5 | 3266 | 185 | 11092 | 2917.75 | 219.5 | 3463.25 | 101604 | 1389.5 | 217 | 1394 | 1756 | 67 |
| vag_C4-like | ano_C3-C4 | son_C3-C4 | 6.6617848 | 1.36E-11 | 0.76266838 | 1039309.75 | 9706.5 | 7983.5 | 3144 | 191 | 11174 | 2681.5 | 226 | 4167.75 | 99946.5 | 1600.5 | 254 | 1333 | 1743 | 57 |
| vag_C4-like | ano_C3-C4 | bro_C4-like | 13.5551011 | 0 | 0.37566599 | 1040186 | 8666.75 | 7132.75 | 3954 | 182 | 11528 | 2342.75 | 211 | 3312.25 | 100536.5 | 1378.5 | 207 | 1162.5 | 1964 | 49 |
| vag_C4-like | ano_C3-C4 | cro_C3 | 6.3529216 | 1.06E-10 | 0.94811514 | 1053119.44 | 11902.8125 | 8576.5 | 2659.25 | 218 | 11726.5 | 2224.75 | 300 | 10164.5625 | 93832.0625 | 2539.125 | 485 | 1208.5 | 1641.5 | 101 |

| | | | | | | | | | | | | | | | | | | | | |
|---|---|---|---|---|---|---|---|---|---|---|---|---|---|---|---|---|---|---|---|---|
| vag_C4-like | ano_C3-C4 | flo_C3-C4 | 4.95407463 | 3.64E-07 | 0.09954927 | 1100620 | 8011 | 6637.25 | 5016.75 | 185 | 12508.75 | 2120 | 172 | 2440.25 | 107166 | 1216.5 | 189 | 1077 | 2211.5 | 61 |
| vag_C4-like | ano_C3-C4 | chl_C3-C4 | 4.97056448 | 3.34E-07 | 0.10156373 | 1101780.5 | 8349.75 | 6760.5 | 4970 | 173.5 | 12590.75 | 2126 | 169 | 2447.5 | 107536 | 1238 | 192 | 1085 | 2225.5 | 55 |
| vag_C4-like | ano_C3-C4 | rob_C3 | 6.19129882 | 3.00E-10 | 0.94995501 | 1035270.25 | 16836.75 | 8360.5 | 2742.5 | 283.5 | 11384 | 2311.5 | 389 | 10492.75 | 91989.25 | 3201.5 | 457 | 1235 | 1590.5 | 109 |
| ang_C3-C4 | ano_C3-C4 | koc_C4 | 4.99551665 | 2.94E-07 | 0.38106565 | 1041120.69 | 10934.8125 | 7893.25 | 3421.5 | 228.5 | 8190.3125 | 2852.3125 | 217.625 | 3202.75 | 100559.25 | 1648.5 | 190 | 1391 | 1324.5 | 52 |
| ang_C3-C4 | ano_C3-C4 | bro_C4-like | 8.77250883 | 0 | 0.27799675 | 1104224.25 | 9171.5 | 7976 | 4072.5 | 187.5 | 9257.75 | 2515.5 | 165 | 3115 | 107039.5 | 1391 | 166 | 1318 | 1612.5 | 51 |
| ang_C3-C4 | ano_C3-C4 | bid_C4 | 5.48420551 | 2.08E-08 | 0.41258741 | 1145891.5 | 12308.5 | 8745.25 | 3728.5 | 216.5 | 9034.75 | 3140.5 | 242 | 3553.5 | 110949.5 | 1851 | 210 | 1511 | 1499.5 | 60 |
| ang_C3-C4 | ano_C3-C4 | aus_C4 | 5.01007627 | 2.72E-07 | 0.45748988 | 1129125 | 12144 | 8615 | 3639.25 | 203 | 8796 | 3170.25 | 225 | 3565.75 | 108344.25 | 1866 | 205.5 | 1470.5 | 1445.5 | 57 |
| ang_C3-C4 | ano_C3-C4 | tri_C4 | 5.2707167 | 6.81E-08 | 0.45675972 | 1118223 | 12343.25 | 8639.25 | 3670.5 | 202.5 | 8835 | 3174.25 | 220 | 3591.5 | 108225.75 | 1853.5 | 204.5 | 1504.5 | 1461.5 | 49 |
| koc_C4 | ano_C3-C4 | son_C3-C4 | 6.13185552 | 4.36E-10 | 0.76692708 | 1027797.13 | 9543.125 | 7906.75 | 3121.75 | 168 | 11009.625 | 2696.625 | 219.75 | 4095.5 | 98848.75 | 1564.5 | 265.5 | 1313.5 | 1652.5 | 52 |
| koc_C4 | ano_C3-C4 | bro_C4-like | 13.5378381 | 0 | 0.36480972 | 1038837.5 | 8647.5 | 7096 | 4006.75 | 169 | 11552 | 2333.5 | 206 | 3294.5 | 100484.75 | 1384.5 | 223.5 | 1160 | 1886.5 | 46 |
| koc_C4 | ano_C3-C4 | cro_C3 | 6.09398672 | 5.53E-10 | 0.94886039 | 1035141.88 | 11593.375 | 8389.5 | 2618.75 | 206 | 11492.5 | 2204.5 | 285 | 9890.625 | 92364.125 | 2484.75 | 498.5 | 1186 | 1501.5 | 97 |
| koc_C4 | ano_C3-C4 | flo_C3-C4 | 5.70718663 | 5.76E-09 | 0.11140806 | 1087010 | 8037.25 | 6554.5 | 4972.5 | 168 | 12359.5 | 2063.25 | 166 | 2428 | 105703.5 | 1187.5 | 197.5 | 1055.5 | 2101 | 61 |
| koc_C4 | ano_C3-C4 | chl_C3-C4 | 6.3205008 | 1.31E-10 | 0.12371922 | 1089768.75 | 8318 | 6660.75 | 4932.75 | 168.5 | 12435 | 2067.75 | 173 | 2472.25 | 106208.25 | 1216 | 202.5 | 1061.5 | 2105 | 58 |
| koc_C4 | ano_C3-C4 | rob_C3 | 6.55011048 | 2.89E-11 | 0.94729502 | 1025054.5 | 16681.5 | 8242 | 2730.5 | 292 | 11259.5 | 2277 | 403.5 | 10428 | 91019 | 3179.5 | 491.5 | 1212.5 | 1486 | 88 |
| son_C3-C4 | ano_C3-C4 | bid_C4 | 8.32056014 | 0 | 0.28058633 | 1145108.75 | 12577.25 | 8852 | 4504.25 | 259 | 10597.25 | 2946 | 227 | 3553.75 | 110472.75 | 1885.5 | 188 | 1447 | 1763.5 | 60 |
| son_C3-C4 | ano_C3-C4 | pal_C4-like | 6.59343962 | 2.16E-11 | 0.23101571 | 1062687.31 | 11371.4375 | 8180.3125 | 4299.5625 | 258.125 | 9982.125 | 2754.625 | 218.75 | 3218.75 | 102211 | 1670 | 185.5 | 1329.5 | 1678 | 56 |
| son_C3-C4 | ano_C3-C4 | aus_C4 | 9.38021609 | 0 | 0.31133466 | 1130197.94 | 12434.5625 | 8685 | 4418 | 245 | 10429.8125 | 2899.0625 | 236.125 | 3585.75 | 108000.25 | 1902.5 | 193 | 1402 | 1714 | 54 |
| son_C3-C4 | ano_C3-C4 | tri_C4 | 9.5597462 | 0 | 0.31233407 | 1128638.63 | 12656.625 | 8731.25 | 4496.25 | 242.5 | 10390.875 | 2942.125 | 253.25 | 3648 | 108921.5 | 1917 | 193 | 1435 | 1748 | 57 |
| bro_C4 _-ike | ano_C3-C4 | bid_C4 | 14.218639 | 0 | 0.65189927 | 1129096 | 12915.25 | 7774.25 | 3493.25 | 214 | 9428 | 2460 | 217.5 | 4395 | 109493.75 | 2067 | 191 | 1249.5 | 1499.5 | 49 |
| bro_C4 _-ike | ano_C3-C4 | ram_C3-C4 | 12.8572941 | 0 | 0.63765267 | 1102985.19 | 9740.5625 | 7673.5625 | 3367.8125 | 202.125 | 9241 | 2446.25 | 183 | 4068 | 106952.5 | 1730 | 197 | 1254.5 | 1418.5 | 41 |
| bro_C4 _-ike | ano_C3-C4 | pal_C4-like | 13.609537 | 0 | 0.62656677 | 1070450.94 | 11827.3125 | 7382.3125 | 3373.8125 | 215.125 | 8929.5 | 2392.5 | 212 | 4039 | 103624 | 1876 | 180 | 1176 | 1456.5 | 49 |
| bro_C4 _-ike | ano_C3-C4 | aus_C4 | 14.0357438 | 0 | 0.65656566 | 1113724 | 12765.5 | 7663.5 | 3445 | 207 | 9282 | 2433.5 | 238 | 4367.25 | 107191.75 | 2073.5 | 187 | 1239 | 1464 | 46 |
| bro_C4 _-ike | ano_C3-C4 | tri_C4 | 14.4002333 | 0 | 0.65491038 | 1104744.5 | 12912.75 | 7666.75 | 3486.25 | 197.5 | 9257.25 | 2441.75 | 244.5 | 4424 | 107130.75 | 2076 | 187 | 1249 | 1490 | 44 |
| cro_C3 | ano_C3-C4 | bid_C4 | 7.60591084 | 1.42E-14 | 0.05992912 | 1140195.19 | 12977.25 | 9372.75 | 10952.0625 | 509 | 12842.3125 | 2414.75 | 326 | 2959 | 102081.063 | 1730 | 226 | 1315.5 | 2771.125 | 104 |
| cro_C3 | ano_C3-C4 | rob_C3 | 4.35815963 | 6.56E-06 | 0.7149278 | 1089897.13 | 15863.5 | 14696.5 | 4837.625 | 500.5 | 10189.625 | 4465 | 412.25 | 5399.5 | 94843.875 | 2337.25 | 349.25 | 2154.25 | 1660.25 | 117.5 |
| cro_C3 | ano_C3-C4 | pal_C4-like | 5.17416768 | 1.15E-07 | 0.04278123 | 1069401.5 | 11870.875 | 8788.625 | 10342 | 482.75 | 12114.9375 | 2315.0625 | 284.625 | 2673.8125 | 95541.9375 | 1552.125 | 214.625 | 1211.625 | 2645.375 | 90.125 |
| cro_C3 | ano_C3-C4 | aus_C4 | 7.12101794 | 5.39E-13 | 0.05852903 | 1123942.88 | 12818.25 | 9211.75 | 10639.375 | 504.5 | 12497.375 | 2435.75 | 310.5 | 2945.75 | 99761.125 | 1750.5 | 225 | 1279 | 2678.25 | 99 |
| cro_C3 | ano_C3-C4 | tri_C4 | 7.22171247 | 2.59E-13 | 0.05923844 | 1115566.13 | 12985.5 | 9249 | 10764.625 | 493.5 | 12540.875 | 2478.75 | 304.5 | 3000.5 | 99727.875 | 1749.5 | 224.5 | 1306 | 2731.75 | 98 |

| | | | | | | | | | | | | | | | | | | | | |
|---|---|---|---|---|---|---|---|---|---|---|---|---|---|---|---|---|---|---|---|---|
| bid_C4 | ano_C3-C4 | flo_C3-C4 | 5.43940759 | 2.68E-08 | 0.09946886 | 1212337.63 | 8906.625 | 7372.8125 | 5580.5625 | 193.125 | 14076.875 | 2274.375 | 186.75 | 2639.5625 | 118344.063 | 1335.125 | 195.125 | 1180.125 | 2371.125 | 57.125 |
| bid_C4 | ano_C3-C4 | chl_C3-C4 | 5.74930319 | 4.49E-09 | 0.10547833 | 1214274 | 9259.5 | 7486 | 5569 | 186.5 | 14151 | 2287 | 189 | 2674 | 118728.5 | 1370 | 198 | 1186 | 2390.5 | 51 |
| bid_C4 | ano_C3-C4 | rob_C3 | 6.21398609 | 2.59E-10 | 0.95157757 | 1142445.69 | 18594.8125 | 9305.5 | 3068 | 311 | 12806.5625 | 2609.3125 | 430.625 | 11623.25 | 101887.75 | 3556 | 496.5 | 1356 | 1694 | 113 |
| flo_C3-C4 | ano_C3-C4 | ram_C3-C4 | 5.93156133 | 1.51E-09 | 0.88582583 | 1188353.44 | 10793.3125 | 7243.8125 | 2568.0625 | 176.625 | 8801.5 | 2178.75 | 153.5 | 5199.25 | 115952.75 | 2033.5 | 192 | 1164 | 1279.5 | 55 |
| flo_C3-C4 | ano_C3-C4 | pal_C4-like | 4.79601096 | 8.10E-07 | 0.90238174 | 1126328.19 | 12762.3125 | 6833.0625 | 2498.3125 | 184.125 | 8319.75 | 2184 | 172 | 5089.5 | 109848.25 | 2125 | 172 | 1094 | 1254.5 | 53 |
| flo_C3-C4 | ano_C3-C4 | aus_C4 | 5.41227113 | 3.12E-08 | 0.9014885 | 1204797 | 14084.25 | 7296.75 | 2645.75 | 181 | 8830 | 2281.75 | 183 | 5612.75 | 116792.25 | 2429 | 194.5 | 1147.5 | 1325.5 | 52 |
| flo_C3-C4 | ano_C3-C4 | tri_C4 | 4.93810556 | 3.95E-07 | 0.9101331 | 1187638.38 | 14157.375 | 7245.0625 | 2607.8125 | 178.625 | 8698.875 | 2276.125 | 179.75 | 5635.3125 | 115864.063 | 2388.125 | 194.625 | 1161.625 | 1329.125 | 52.125 |
| chl_C3-C4 | ano_C3-C4 | ram_C3-C4 | 5.98264019 | 1.10E-09 | 0.88251168 | 1188658.94 | 10879.3125 | 7347.0625 | 2600.3125 | 192.625 | 9192.25 | 2205.75 | 141.5 | 5169.5 | 116227.75 | 2012.5 | 196.5 | 1190 | 1289 | 45 |
| chl_C3-C4 | ano_C3-C4 | pal_C4-like | 5.06989727 | 1.99E-07 | 0.89534313 | 1127088.44 | 12787.5625 | 6943.3125 | 2535.5625 | 196.125 | 8585.75 | 2202.25 | 179.5 | 5053.75 | 110130.25 | 2141 | 170.5 | 1099 | 1276 | 53 |
| chl_C3-C4 | ano_C3-C4 | aus_C4 | 5.2564195 | 7.36E-08 | 0.90284068 | 1206073 | 14145.25 | 7424.25 | 2645.75 | 196 | 9164.75 | 2291.75 | 183 | 5581.25 | 117128.5 | 2432 | 187.5 | 1160.5 | 1352.5 | 50 |
| chl_C3-C4 | ano_C3-C4 | tri_C4 | 4.90792386 | 4.61E-07 | 0.90973076 | 1188792.75 | 14228.5 | 7367.5 | 2617 | 190.5 | 9021.75 | 2286.75 | 181 | 5615 | 116208.25 | 2405.5 | 185.5 | 1169.5 | 1354.5 | 50 |
| rob_C3 | ano_C3-C4 | aus_C4 | 6.06776269 | 6.51E-10 | 0.0488691 | 1128417 | 12748.25 | 9184.5 | 11313.25 | 523.5 | 18209.5 | 2608.5 | 394 | 3055.75 | 99767.25 | 1707.5 | 318 | 1288 | 3484 | 107 |
| rob_C3 | ano_C3-C4 | tri_C4 | 6.19208431 | 2.98E-10 | 0.04973434 | 1127364.5 | 12968.5 | 9261.5 | 11469 | 514.5 | 18300.75 | 2660.75 | 411.5 | 3121.75 | 100588.25 | 1716.5 | 327 | 1330 | 3502.5 | 111 |
| vag_C4-like | bid_C4 | pal_C4-like | 9.28173204 | 0 | 0.63898356 | 1056013.56 | 4618.625 | 4040.6875 | 1542.625 | 35.25 | 4265.75 | 1092.4375 | 32.375 | 1889.25 | 114642.938 | 796.875 | 48 | 620.875 | 798.375 | 13.375 |
| vag_C4-like | bid_C4 | aus_C4 | 5.52721295 | 1.63E-08 | 0.67372948 | 1080415.5 | 4750.6875 | 4207.5 | 1486.6875 | 32.375 | 4257.4375 | 1225.875 | 35.75 | 1764.4375 | 116047.875 | 824.25 | 35.875 | 676.25 | 743.25 | 14.25 |
| vag_C4-like | bid_C4 | tri_C4 | 6.20941665 | 2.67E-10 | 0.65539859 | 1080510.88 | 4900.8125 | 4206.875 | 1537.5625 | 34.125 | 4267.3125 | 1238.75 | 37.5 | 1807.0625 | 117421 | 820.5 | 39.125 | 703.5 | 754.5 | 17.5 |
| koc_C4 | bid_C4 | pal_C4-like | 8.88015047 | 0 | 0.43664706 | 1042382.25 | 4455.5 | 4216.1875 | 1752.6875 | 31.875 | 4611.375 | 1154.125 | 32.25 | 1618.0625 | 112747.563 | 734.625 | 34.875 | 660.375 | 808.875 | 12.375 |
| koc_C4 | bid_C4 | aus_C4 | 10.0717173 | 0 | 0.4125211 | 1075699.75 | 4816.4375 | 4277.5 | 1861.4375 | 56.375 | 4777.9375 | 1121.875 | 52.25 | 1641.1875 | 115174.875 | 796.75 | 33.875 | 678.75 | 813.75 | 11.25 |
| koc_C4 | bid_C4 | tri_C4 | 10.2486369 | 0 | 0.40301291 | 1071233.56 | 5012.1875 | 4280.375 | 1906.625 | 50.75 | 4768.6875 | 1126.4375 | 51.875 | 1653.125 | 116064.625 | 827.25 | 36.25 | 693.25 | 816.75 | 13.25 |
| pal_C4-like | bid_C4 | aus_C4 | 8.1377606 | 2.22E-16 | 0.46146965 | 1105778.06 | 4887.3125 | 4390.1875 | 1802.3125 | 47.125 | 4749.9375 | 1275.5625 | 30.125 | 1726.9375 | 118807.563 | 842.125 | 35.875 | 690.125 | 773.625 | 9.125 |
| pal_C4-like | bid_C4 | tri_C4 | 8.53329159 | 0 | 0.45749158 | 1102008.19 | 5114.75 | 4369.8125 | 1848.25 | 48.5 | 4765.375 | 1284.3125 | 33.125 | 1759.875 | 119804.063 | 872.125 | 36.75 | 697.625 | 785.125 | 12.125 |
| vag_C4-like | bro_C4_-ike | son_C3-C4 | 9.108525 | 0 | 0.07826831 | 989401.938 | 4754.3125 | 4514.0625 | 7584.8125 | 111.625 | 12758.25 | 1461 | 114 | 1981 | 96649 | 784.5 | 149.5 | 712 | 2552 | 29 |
| vag_C4-like | bro_C4_-ike | cro_C3 | 6.72339263 | 8.93E-12 | 0.93386393 | 1013846.44 | 11166.0625 | 9434.75 | 2888.75 | 249 | 12026.5 | 2412.5 | 293 | 9137.3125 | 90321.5625 | 2406.625 | 464 | 1276 | 1684.5 | 93 |
| vag_C4-like | bro_C4_-ike | rob_C3 | 7.72877921 | 5.55E-15 | 0.92513769 | 983783 | 15620.75 | 9004.75 | 2969.5 | 320.5 | 11618.25 | 2419 | 362 | 9222 | 87488.75 | 2947.5 | 446.5 | 1251 | 1635.5 | 108 |
| ang_C3-C4 | bro_C4_-ike | son_C3-C4 | 7.98740232 | 6.66E-16 | 0.06862902 | 1048290.94 | 5125.8125 | 4995.5625 | 7679.8125 | 99.625 | 10347.5 | 1488 | 97 | 1944.25 | 102934 | 806.5 | 121 | 795 | 2155 | 28 |
| koc_C4 | bro_C4_-ike | son_C3-C4 | 7.93771546 | 1.11E-15 | 0.06846534 | 992548.313 | 4699.9375 | 4565.8125 | 7652.5625 | 92.625 | 12839.375 | 1487.375 | 111.75 | 1940.5 | 97200.25 | 774.5 | 142.5 | 733 | 2485.5 | 30 |
| koc_C4 | bro_C4_-ike | cro_C3 | 6.63467488 | 1.63E-11 | 0.93368893 | 1005538.63 | 11023.375 | 9405.75 | 2860.25 | 243 | 11990 | 2392.5 | 279 | 8978.625 | 89893.125 | 2367.25 | 452 | 1278 | 1590.5 | 90 |
| koc_C4 | bro_C4_-ike | rob_C3 | 8.24512074 | 1.11E-16 | 0.92200353 | 984451.75 | 15706 | 9032.5 | 2974.25 | 328.5 | 11701.25 | 2388.75 | 379.5 | 9310 | 87644.5 | 2957.5 | 446.5 | 1273 | 1553 | 100 |

| | | | | | | | | | | | | | | | | | | | | |
|---|---|---|---|---|---|---|---|---|---|---|---|---|---|---|---|---|---|---|---|---|
| son_C3-C4 | bro_C4_-ike | bid_C4 | 9.03029615 | 0 | 0.92533962 | 1077118.44 | 14240 | 5024.3125 | 2142.75 | 144 | 5187.0625 | 1604.5 | 113 | 8275.5625 | 105536.75 | 2713.5 | 104.625 | 782 | 879.5 | 29 |
| son_C3-C4 | bro_C4_-ike | flo_C3-C4 | 48.3247731 | 0 | 0.58265765 | 1085226.69 | 8852 | 2284.0625 | 4887.5 | 74.5 | 6361.5625 | 477 | 43 | 6634.5625 | 107565.5 | 1824.5 | 56.625 | 282 | 1425.5 | 18 |
| son_C3-C4 | bro_C4_-ike | chl_C3-C4 | 48.5109631 | 0 | 0.58342184 | 1086023.88 | 9121.8125 | 2250.125 | 4971.8125 | 67.125 | 6385.8125 | 510 | 38 | 6758.8125 | 107841.5 | 1861 | 49.625 | 308 | 1419.5 | 14 |
| son_C3-C4 | bro_C4_-ike | ano_C3-C4 | 28.0008013 | 0 | 0.77379515 | 1084780.44 | 9871 | 4271.0625 | 2912.5 | 117.5 | 5749.5625 | 1121 | 73 | 7249.3125 | 107124.5 | 2152 | 80.625 | 621 | 1066.5 | 20 |
| son_C3-C4 | bro_C4_-ike | ram_C3-C4 | 8.36869206 | 0 | 0.92951731 | 1063149.38 | 11254.375 | 5102.5625 | 1984.3125 | 110.625 | 5102.875 | 1502.625 | 105.25 | 7855.0625 | 104462.313 | 2373.625 | 107.625 | 776.125 | 830.125 | 29.125 |
| son_C3-C4 | bro_C4_-ike | pub_C3-C4 | 46.0853158 | 0 | 0.58721764 | 1075656.38 | 8445.5625 | 2484.9375 | 4741.125 | 57.75 | 6240.375 | 619.3125 | 42.625 | 6482.9375 | 107312.625 | 1836.25 | 49.875 | 348.25 | 1381.75 | 14.25 |
| son_C3-C4 | bro_C4_-ike | pal_C4-like | 10.2301296 | 0 | 0.91319574 | 1020944.31 | 13098.875 | 4699.0625 | 2088.25 | 145.5 | 4994.6875 | 1493.375 | 104.75 | 7751.5625 | 99828.5 | 2533 | 108.125 | 735 | 834 | 27 |
| son_C3-C4 | bro_C4_-ike | aus_C4 | 8.59264504 | 0 | 0.92849359 | 1052917.38 | 13872.5625 | 4887.8125 | 2067 | 151 | 5102.625 | 1562.5625 | 107.125 | 8112.5625 | 101999.25 | 2676.5 | 106.625 | 744 | 848 | 27 |
| son_C3-C4 | bro_C4_-ike | tri_C4 | 8.80816861 | 0 | 0.92800381 | 1056781.31 | 14227.125 | 4879.8125 | 2098 | 156 | 5037.6875 | 1577.625 | 113.25 | 8285.0625 | 103722 | 2751.5 | 103.625 | 765 | 863 | 28 |
| cro_C3 | bro_C4_-ike | bid_C4 | 7.31566581 | 1.29E-13 | 0.06967861 | 1087597.19 | 13300.5 | 10227 | 9754.3125 | 476.5 | 11985.8125 | 2584.5 | 311 | 3121.5 | 97328.5625 | 1776 | 278 | 1378 | 2605.125 | 95 |
| cro_C3 | bro_C4_-ike | rob_C3 | 6.72701056 | 8.71E-12 | 0.61871102 | 1035343.63 | 15001.75 | 14298.5 | 4719.875 | 502.75 | 9770.875 | 4146.75 | 378.75 | 5076.75 | 90492.125 | 2221 | 371.75 | 2063 | 1640.25 | 118.25 |
| cro_C3 | bro_C4_-ike | pal_C4-like | 5.51620476 | 1.74E-08 | 0.05469197 | 1036788.13 | 12264.5 | 9725.75 | 9302.625 | 469 | 11437.875 | 2484 | 283 | 2878.5 | 92630.375 | 1626 | 249 | 1323 | 2495.25 | 74 |
| cro_C3 | bro_C4_-ike | aus_C4 | 6.98320055 | 1.45E-12 | 0.06980372 | 1062163.88 | 12971.25 | 9984 | 9343.375 | 493.5 | 11580.125 | 2560.5 | 285.5 | 3069.5 | 94098.625 | 1748.5 | 252.5 | 1322.5 | 2499.25 | 92 |
| cro_C3 | bro_C4_-ike | tri_C4 | 7.73638673 | 5.11E-15 | 0.07624319 | 1057423.38 | 13201.25 | 9987 | 9470.625 | 506.5 | 11638.625 | 2585.75 | 294 | 3154 | 94702.625 | 1771.5 | 262.5 | 1351.5 | 2528.75 | 92 |
| bid_C4 | bro_C4_-ike | rob_C3 | 7.19239405 | 3.21E-13 | 0.93230781 | 1071086.13 | 17080.875 | 9897.8125 | 3224.5625 | 350.125 | 12963.875 | 2686.875 | 405.75 | 10092.3125 | 95560.5625 | 3217.625 | 471.125 | 1370.125 | 1734.125 | 116.125 |
| rob_C3 | bro_C4_-ike | pal_C4-like | 5.44544651 | 2.59E-08 | 0.05327046 | 1012774.69 | 11838.0625 | 9322.75 | 9568.25 | 449 | 16141.0625 | 2553.8125 | 361.625 | 2948.5 | 90209.75 | 1594.5 | 337.5 | 1297 | 3052.5 | 102 |
| rob_C3 | bro_C4_-ike | aus_C4 | 7.12222373 | 5.35E-13 | 0.06916672 | 1048034.75 | 12722.25 | 9680.75 | 9740 | 497.5 | 16648.75 | 2641 | 370 | 3168.5 | 92536 | 1712.5 | 343.5 | 1315 | 3127.5 | 105 |
| rob_C3 | bro_C4_-ike | tri_C4 | 7.89484893 | 1.44E-15 | 0.07535843 | 1051600 | 13022.25 | 9765.75 | 9912.75 | 513.5 | 16750.5 | 2673.5 | 376 | 3263.5 | 93918.25 | 1758.5 | 355.5 | 1351 | 3159 | 112 |
| vag_C4-like | chl_C3-C4 | son_C3-C4 | 5.62785768 | 9.15E-09 | 0.74490456 | 1037024.25 | 9637.25 | 9405.5 | 3219.5 | 205 | 11393.75 | 2825.25 | 246 | 3976.5 | 99504.5 | 1543.5 | 250.5 | 1412.5 | 1771 | 50 |
| vag_C4-like | chl_C3-C4 | bro_C4-like | 10.6074974 | 0 | 0.13920495 | 1042511.88 | 6736.125 | 6695.625 | 5950.875 | 159.75 | 12474.5625 | 1832.5625 | 165.125 | 2498.5625 | 101332.313 | 1014.125 | 149.125 | 932.125 | 2385.125 | 36.125 |
| vag_C4-like | chl_C3-C4 | cro_C3 | 4.90410819 | 4.70E-07 | 0.95706608 | 1054562.44 | 11901.3125 | 10092.5 | 2726.5 | 242 | 11980.25 | 2382.5 | 301 | 10050.8125 | 93611.0625 | 2526.625 | 484 | 1322 | 1667 | 86 |
| vag_C4-like | chl_C3-C4 | rob_C3 | 5.00457425 | 2.80E-07 | 0.95617028 | 1034807 | 16776 | 9815.5 | 2814.75 | 296 | 11645.75 | 2458.25 | 387 | 10235.5 | 91776.25 | 3155.5 | 466.5 | 1324.5 | 1611.5 | 100 |
| ang_C3-C4 | chl_C3-C4 | bro_C4-like | 6.32632742 | 1.26E-10 | 0.08916067 | 1108935.94 | 7114.8125 | 7385.8125 | 6208.5625 | 172.125 | 10104.5 | 2038 | 129 | 2446.25 | 107999.5 | 1058.5 | 121 | 1046 | 2012 | 39 |
| koc_C4 | chl_C3-C4 | son_C3-C4 | 4.76806045 | 9.31E-07 | 0.75448718 | 1028235.13 | 9476.875 | 9386 | 3215.25 | 193 | 11285.125 | 2880.125 | 228.75 | 3910 | 98560.75 | 1504.5 | 232.5 | 1401.5 | 1695.5 | 51 |
| koc_C4 | chl_C3-C4 | bro_C4-like | 9.437715 | 0 | 0.12378046 | 1045081.13 | 6727.875 | 6753.875 | 6027.625 | 161.75 | 12591.3125 | 1845.8125 | 160.125 | 2436.5625 | 101653.313 | 1033.125 | 160.125 | 941.625 | 2295.625 | 37.125 |
| koc_C4 | chl_C3-C4 | cro_C3 | 4.82377422 | 7.05E-07 | 0.95645925 | 1037087.88 | 11578.125 | 9960 | 2704 | 235 | 11776 | 2367 | 288 | 9769.875 | 92158.375 | 2464.75 | 479 | 1287 | 1535 | 86 |
| koc_C4 | chl_C3-C4 | rob_C3 | 4.74857359 | 1.03E-06 | 0.95781784 | 1025485.5 | 16609.5 | 9728.5 | 2818.5 | 311 | 11551 | 2479.25 | 396.5 | 10182.5 | 90758.75 | 3126.5 | 475.5 | 1312 | 1496 | 94 |
| son_C3-C4 | chl_C3-C4 | bid_C4 | 6.35578175 | 1.04E-10 | 0.28567078 | 1147513.25 | 12888.25 | 10507.5 | 4323 | 247.5 | 10592 | 3150.25 | 231 | 3619.25 | 110473.5 | 1941.5 | 209 | 1559.5 | 1716.5 | 54 |

| | | | | | | | | | | | | | | | | | | | | |
|---|---|---|---|---|---|---|---|---|---|---|---|---|---|---|---|---|---|---|---|---|
| son_C3-C4 | chl_C3-C4 | pal_C4-like | 4.27931723 | 9.38E-06 | 0.20135932 | 1064418.63 | 11613.125 | 9700.75 | 4161 | 251 | 9961.625 | 2941.875 | 233.75 | 3249.25 | 102235.5 | 1748 | 216.5 | 1429 | 1611 | 44 |
| son_C3-C4 | chl_C3-C4 | aus_C4 | 7.37402045 | 8.35E-14 | 0.31809559 | 1135126.94 | 12763.0625 | 10294.25 | 4271.5 | 262 | 10434.5625 | 3103.3125 | 248.125 | 3648.25 | 108368 | 1966.5 | 215 | 1513.5 | 1661 | 48 |
| son_C3-C4 | chl_C3-C4 | tri_C4 | 7.77912298 | 3.66E-15 | 0.3268986 | 1127152.88 | 12894.875 | 10255.25 | 4295.5 | 259 | 10346.875 | 3105.625 | 254.75 | 3683.5 | 108626.25 | 1962 | 211 | 1539.5 | 1694 | 50 |
| bro_C4 -ike | chl_C3-C4 | bid_C4 | 10.6106865 | 0 | 0.87149234 | 1135313.13 | 14001.0625 | 7313.375 | 2622.5625 | 162.125 | 7347.375 | 1937.0625 | 164.625 | 6585.875 | 110668.063 | 2526.125 | 172.75 | 1006.625 | 1107.125 | 41.125 |
| bro_C4 -ike | chl_C3-C4 | ano_C3-C4 | 24.8944856 | 0 | 0.59667949 | 1151076.13 | 8087.3125 | 6583.625 | 3466.5625 | 118.625 | 7811.875 | 1599.0625 | 106.125 | 4361.875 | 114813.063 | 1603.125 | 126.75 | 878.125 | 1294.625 | 40.125 |
| bro_C4 -ike | chl_C3-C4 | ram_C3-C4 | 9.46676443 | 0 | 0.8756551 | 1106496.63 | 10739.3125 | 7232.875 | 2549.0625 | 139.125 | 7155.625 | 1942.5625 | 168.625 | 6213.625 | 107913.313 | 2098.625 | 175.75 | 978.625 | 1067.125 | 35.125 |
| bro_C4 -ike | chl_C3-C4 | pal_C4-like | 10.5784055 | 0 | 0.86051639 | 1075513.63 | 12782.8125 | 6991.375 | 2556.8125 | 162.625 | 6974.875 | 1884.5625 | 170.125 | 6031.875 | 104724.063 | 2326.125 | 176.75 | 939.125 | 1075.125 | 31.125 |
| bro_C4 -ike | chl_C3-C4 | aus_C4 | 10.7039497 | 0 | 0.86970638 | 1121708.44 | 13837 | 7205.5625 | 2614.5 | 159 | 7232.3125 | 1924.75 | 186 | 6528.8125 | 108594.5 | 2542.5 | 181.125 | 993.5 | 1078 | 34 |
| bro_C4 -ike | chl_C3-C4 | tri_C4 | 10.8907634 | 0 | 0.86818423 | 1104156.44 | 13911.75 | 7148.5625 | 2597.5 | 152 | 7199.5625 | 1899 | 188.5 | 6499.5625 | 107708.5 | 2524 | 173.125 | 994.5 | 1100 | 33 |
| cro_C3 | chl_C3-C4 | bid_C4 | 5.61784149 | 9.69E-09 | 0.04783142 | 1143725.69 | 13317.25 | 11048.5 | 10806.5625 | 519 | 12890.0625 | 2600 | 331 | 3012.25 | 102095.063 | 1756.5 | 269 | 1437 | 2770.125 | 90 |
| cro_C3 | chl_C3-C4 | aus_C4 | 5.4628965 | 2.35E-08 | 0.04811656 | 1128774.13 | 13175 | 10880.75 | 10549.875 | 524.5 | 12576.625 | 2612 | 313.5 | 3013.25 | 100016.125 | 1785 | 248.5 | 1406.5 | 2681.25 | 90 |
| cro_C3 | chl_C3-C4 | tri_C4 | 5.8588251 | 2.34E-09 | 0.05155856 | 1113850.13 | 13228 | 10756.5 | 10556.875 | 513 | 12520.875 | 2619.25 | 308 | 3050.75 | 99394.875 | 1765 | 256.5 | 1417.5 | 2703.75 | 85 |
| bid_C4 | chl_C3-C4 | rob_C3 | 4.34760532 | 6.89E-06 | 0.96292571 | 1145098.94 | 18591.0625 | 10969.25 | 3133 | 340 | 13137.8125 | 2803.5625 | 424.125 | 11360 | 101913.25 | 3504 | 513 | 1473 | 1721 | 113 |
| rob_C3 | chl_C3-C4 | aus_C4 | 4.70621248 | 1.26E-06 | 0.04095534 | 1133528 | 13075.75 | 10817 | 11135.75 | 542.5 | 18278.5 | 2793.5 | 403 | 3149.75 | 100113.75 | 1736.5 | 324 | 1428 | 3440 | 112 |
| rob_C3 | chl_C3-C4 | tri_C4 | 5.06068911 | 2.09E-07 | 0.04383663 | 1125379.5 | 13234 | 10766 | 11185.5 | 532.5 | 18216.25 | 2804.25 | 404.5 | 3188.5 | 100226.5 | 1742 | 336 | 1453 | 3438.5 | 108 |
| vag_C4-like | flo_C3-C4 | son_C3-C4 | 6.74801309 | 7.54E-12 | 0.70691531 | 1035213 | 9546 | 9038.75 | 3254.5 | 212 | 11346.5 | 2783 | 239 | 3920.25 | 99057.5 | 1550.5 | 261 | 1398 | 1749 | 55 |
| vag_C4-like | flo_C3-C4 | bro_C4-like | 10.3832956 | 0 | 0.13380683 | 1039803.75 | 6668.25 | 6415.5 | 5969.5 | 167 | 12405.5 | 1792.5 | 172 | 2437.75 | 100820.75 | 992.5 | 158 | 892 | 2359 | 43 |
| vag_C4-like | flo_C3-C4 | cro_C3 | 5.26383933 | 7.07E-08 | 0.95335923 | 1050046.19 | 11676.5625 | 9749.5 | 2735.75 | 254.5 | 11867.25 | 2367.25 | 300 | 9899.5625 | 93085.8125 | 2524.125 | 490 | 1289.5 | 1629 | 96 |
| vag_C4-like | flo_C3-C4 | rob_C3 | 5.0200796 | 2.59E-07 | 0.95570287 | 1031005 | 16599 | 9456.25 | 2814.75 | 316 | 11526.5 | 2457.25 | 388 | 10170.25 | 91059.5 | 3163.5 | 460 | 1305.5 | 1572.5 | 111 |
| ang_C3-C4 | flo_C3-C4 | bro_C4-like | 6.44458608 | 5.82E-11 | 0.08888647 | 1105806.5 | 7111.75 | 7137.75 | 6144.25 | 172.5 | 10056.75 | 1957 | 126 | 2365.5 | 107650 | 1027.5 | 140.5 | 1008 | 2016 | 38 |
| koc_C4 | flo_C3-C4 | son_C3-C4 | 6.80369068 | 5.13E-12 | 0.69080984 | 1027010.38 | 9417.625 | 9106.5 | 3247.25 | 194 | 11263.875 | 2772.875 | 230.75 | 3832.75 | 98227 | 1517.5 | 249 | 1382 | 1675.5 | 53 |
| koc_C4 | flo_C3-C4 | bro_C4-like | 9.98395398 | 0 | 0.1269302 | 1041289 | 6687 | 6548.5 | 6014.25 | 167 | 12504.75 | 1773.75 | 171 | 2390.25 | 101057 | 1008.5 | 173 | 893.5 | 2274.5 | 41 |
| koc_C4 | flo_C3-C4 | cro_C3 | 5.57983967 | 1.21E-08 | 0.94970332 | 1032498.13 | 11370.375 | 9691.25 | 2701.25 | 245.5 | 11686.5 | 2314.5 | 283 | 9617.125 | 91659.125 | 2461.25 | 473 | 1263 | 1508 | 91 |
| koc_C4 | flo_C3-C4 | rob_C3 | 5.33822821 | 4.70E-08 | 0.95313903 | 1022392.25 | 16469.25 | 9470.5 | 2803.5 | 317 | 11477.25 | 2425.25 | 385.75 | 10118.75 | 90170.25 | 3139.75 | 455 | 1299 | 1476.25 | 102.25 |
| son_C3-C4 | flo_C3-C4 | bid_C4 | 7.84716938 | 2.22E-15 | 0.32505612 | 1145931.25 | 12838 | 10112.5 | 4261.75 | 266 | 10507.75 | 3059 | 234 | 3638.25 | 110141.5 | 1903.5 | 215 | 1527 | 1725.5 | 64 |
| son_C3-C4 | flo_C3-C4 | pal_C4-like | 6.36614238 | 9.74E-11 | 0.26628665 | 1064196.13 | 11628.625 | 9394.25 | 4084.5 | 261 | 9892.375 | 2845.625 | 232.75 | 3295.25 | 102017.5 | 1707 | 215.5 | 1412.5 | 1622 | 50 |
| son_C3-C4 | flo_C3-C4 | aus_C4 | 8.68630427 | 0 | 0.35446715 | 1134012.44 | 12756.0625 | 9927.25 | 4213.5 | 273 | 10369.3125 | 3033.0625 | 240.125 | 3681.25 | 108050.5 | 1940.5 | 226 | 1490 | 1663 | 62 |
| son_C3-C4 | flo_C3-C4 | tri_C4 | 9.07701033 | 0 | 0.35919578 | 1125758.63 | 12854.875 | 9904.5 | 4245.5 | 270 | 10280.625 | 3030.375 | 252.75 | 3711.5 | 108233.25 | 1927 | 222 | 1509 | 1699 | 62 |

| | | | | | | | | | | | | | | | | | | | | |
|---|---|---|---|---|---|---|---|---|---|---|---|---|---|---|---|---|---|---|---|---|
| bro_C4_-ike | flo_C3-C4 | bid_C4 | 10.7479586 | 0 | 0.87145909 | 1131409.38 | 13898.875 | 7045.875 | 2582.125 | 171.75 | 7300.5625 | 1893.8125 | 168.625 | 6560.3125 | 110037.063 | 2497.625 | 172.125 | 956.125 | 1090.625 | 51.125 |
| bro_C4_-ike | flo_C3-C4 | ano_C3-C4 | 25.4142081 | 0 | 0.5922962 | 1148263.5 | 7988 | 6313 | 3454.5 | 128 | 7785 | 1544 | 105 | 4319.5 | 114321.5 | 1593 | 133 | 845 | 1278 | 35 |
| bro_C4_-ike | flo_C3-C4 | ram_C3-C4 | 9.18883667 | 0 | 0.88048919 | 1103914.5 | 10667.5 | 6980.75 | 2455.5 | 165.5 | 7165.75 | 1876.5 | 165.5 | 6142.25 | 107358.75 | 2111.5 | 178 | 947.5 | 1039.5 | 38 |
| bro_C4_-ike | flo_C3-C4 | pal_C4-like | 10.7895834 | 0 | 0.86238999 | 1073849.25 | 12743.5 | 6754.5 | 2491.75 | 171 | 6933 | 1815.5 | 178 | 6053.5 | 104328.5 | 2297 | 172 | 913 | 1057.5 | 33 |
| bro_C4_-ike | flo_C3-C4 | aus_C4 | 10.6162806 | 0 | 0.87287499 | 1118566.25 | 13783 | 6957 | 2554.5 | 172 | 7195.25 | 1877.75 | 187 | 6524.5 | 108172.25 | 2520.5 | 185.5 | 942.5 | 1064 | 44 |
| bro_C4_-ike | flo_C3-C4 | tri_C4 | 11.3529115 | 0 | 0.86591795 | 1101481.63 | 13821.625 | 6903.125 | 2555.875 | 166.75 | 7155.5625 | 1835.8125 | 193.625 | 6486.0625 | 107174.313 | 2493.625 | 181.625 | 939.125 | 1083.125 | 40.125 |
| cro_C3 | flo_C3-C4 | bid_C4 | 6.08253661 | 5.94E-10 | 0.05181665 | 1138725.31 | 13208.125 | 10721.625 | 10656.6875 | 514.75 | 12676.375 | 2550.3125 | 325.125 | 2993.3125 | 101598.125 | 1724.125 | 280.625 | 1395.625 | 2761.75 | 101.125 |
| cro_C3 | flo_C3-C4 | pal_C4-like | 4.32944734 | 7.48E-06 | 0.03841623 | 1068877.38 | 12060.5 | 10077.5 | 10080.875 | 496 | 11999.125 | 2427.75 | 294.5 | 2733.5 | 95184.125 | 1540.5 | 261.5 | 1297 | 2639.75 | 82 |
| cro_C3 | flo_C3-C4 | aus_C4 | 6.20960493 | 2.67E-10 | 0.05465584 | 1124934.38 | 13109 | 10567.75 | 10410.625 | 520.5 | 12377.625 | 2566.75 | 310.5 | 3020.25 | 99583.375 | 1764 | 269.5 | 1366 | 2665.75 | 108 |
| cro_C3 | flo_C3-C4 | tri_C4 | 6.63398807 | 1.64E-11 | 0.05823566 | 1109748.75 | 13124.875 | 10452.625 | 10400 | 508.75 | 12303.9375 | 2560.8125 | 307.125 | 3045.5625 | 98955.1875 | 1727.625 | 277.625 | 1372.625 | 2691.375 | 107.125 |
| bid_C4 | flo_C3-C4 | rob_C3 | 4.53360764 | 2.90E-06 | 0.96171499 | 1141618.06 | 18459.875 | 10616.125 | 3087.8125 | 355.125 | 13049.4375 | 2747.375 | 419.5 | 11299.125 | 101329.063 | 3517.375 | 513.25 | 1435.625 | 1692.875 | 124.375 |
| chl_C3-C4 | flo_C3-C4 | pub_C3-C4 | 5.47122468 | 2.24E-08 | 0.53474734 | 1207337.06 | 4012.1875 | 4042.125 | 1650.625 | 41.25 | 4559.125 | 1353.125 | 48.25 | 1695.0625 | 128489.563 | 664.625 | 30.125 | 629.625 | 743.625 | 10.625 |
| rob_C3 | flo_C3-C4 | aus_C4 | 4.69080587 | 1.36E-06 | 0.04063381 | 1131359.25 | 13024.75 | 10482 | 11103.75 | 524.5 | 18167.5 | 2763.5 | 387.25 | 3116.75 | 99619.5 | 1728.75 | 342 | 1408 | 3443.25 | 119.25 |
| rob_C3 | flo_C3-C4 | tri_C4 | 5.42932182 | 2.84E-08 | 0.04652121 | 1123122.38 | 13150.875 | 10437.125 | 11141.875 | 520.25 | 18101.0625 | 2754.0625 | 388.875 | 3163.3125 | 99724.3125 | 1719.875 | 358.125 | 1421.625 | 3452.875 | 117.375 |
| bro_C4_-ike | pal_C4-like | aus_C4 | 4.56924252 | 2.45E-06 | 0.02294693 | 1051071.63 | 5350.125 | 4932.375 | 10004.125 | 170.75 | 11778.3125 | 988.0625 | 98.125 | 1199.8125 | 102533.063 | 746.125 | 91.625 | 628.125 | 2738.625 | 31.125 |
| bro_C4_-ike | pal_C4-like | tri_C4 | 4.38951745 | 5.68E-06 | 0.02225858 | 1045421 | 5562.25 | 4924.5 | 10053.75 | 179 | 11802.625 | 1015.875 | 103.75 | 1221.625 | 103095.875 | 776.75 | 91.75 | 648.25 | 2767.75 | 26.25 |
| bid_C4 | pal_C4-like | flo_C3-C4 | 4.01615065 | 2.96E-05 | 0.98088704 | 1121906.31 | 12512.25 | 4958.375 | 1133.4375 | 86.375 | 5308.1875 | 951.5 | 90.5 | 10288.625 | 111032.688 | 2868.875 | 170.75 | 622.875 | 714.875 | 29.375 |
| bid_C4 | pal_C4-like | chl_C3-C4 | 4.82879491 | 6.88E-07 | 0.97751141 | 1124148.44 | 12793.375 | 4966.5625 | 1143.875 | 88.75 | 5368.3125 | 926.125 | 94.75 | 10391.0625 | 111457.125 | 2924.75 | 169.625 | 627.75 | 723.25 | 25.25 |
| flo_C3-C4 | pal_C4-like | aus_C4 | 5.21493888 | 9.21E-08 | 0.02781522 | 1107732.13 | 5661.125 | 5182.625 | 9853.375 | 172.75 | 12203.0625 | 1063.0625 | 95.125 | 1314.5625 | 108660.563 | 806.125 | 95.125 | 641.125 | 2768.125 | 25.125 |
| flo_C3-C4 | pal_C4-like | tri_C4 | 5.52197857 | 1.68E-08 | 0.02971838 | 1091847.13 | 5801.125 | 5118.8125 | 9746.8125 | 178.125 | 12095 | 1056 | 93 | 1322.1875 | 107900.438 | 807.375 | 91.375 | 649.375 | 2730.875 | 24.375 |
| chl_C3-C4 | pal_C4-like | aus_C4 | 5.5936717 | 1.11E-08 | 0.02925984 | 1109658.88 | 5703.125 | 5192.125 | 9931.375 | 175.75 | 12456.0625 | 1040.0625 | 103.125 | 1308.0625 | 109049.813 | 810.125 | 98.125 | 649.125 | 2811.125 | 23.125 |
| chl_C3-C4 | pal_C4-like | tri_C4 | 6.19704863 | 2.89E-10 | 0.03240317 | 1093492.25 | 5839.25 | 5116.25 | 9869 | 179 | 12329.125 | 1022.625 | 97.25 | 1318.875 | 108295.625 | 815.75 | 97.25 | 655.25 | 2780.25 | 20.25 |
| ano_C3-C4 | pal_C4-like | aus_C4 | 5.01771349 | 2.62E-07 | 0.02766738 | 1102712.81 | 5620.125 | 5177.1875 | 9453.375 | 162.75 | 10735.375 | 1043.0625 | 74.125 | 1282.375 | 108713.563 | 817.125 | 95.75 | 642.125 | 2624.125 | 19.125 |
| ano_C3-C4 | pal_C4-like | tri_C4 | 4.87678337 | 5.40E-07 | 0.02700865 | 1092662.44 | 5811 | 5137.3125 | 9458 | 169 | 10780.6875 | 1052.875 | 76.75 | 1286.1875 | 108579.375 | 832.75 | 98.875 | 647.25 | 2608.25 | 18.25 |
| vag_C4-like | pub_C3-C4 | son_C3-C4 | 8.39165444 | 0 | 0.69138582 | 1016548.5 | 9358.5 | 8456.25 | 3224.25 | 220.5 | 11090.625 | 2641.625 | 221.25 | 3946.875 | 97844.875 | 1557.75 | 255.25 | 1302.25 | 1737.25 | 59.25 |
| vag_C4-like | pub_C3-C4 | bro_C4-like | 12.876327 | 0 | 0.16523572 | 1040508 | 6876 | 6174.5 | 5801.5 | 160 | 12344.5 | 1742.25 | 161 | 2545.75 | 101619.5 | 1078.5 | 176 | 871.5 | 2337 | 42 |
| vag_C4-like | pub_C3-C4 | cro_C3 | 7.73173791 | 5.33E-15 | 0.93527686 | 1042708.94 | 11671.8125 | 9222.5 | 2745.75 | 236.5 | 11819.25 | 2215 | 297 | 9884.5625 | 93084.5625 | 2575.125 | 480 | 1261 | 1612 | 88 |
| vag_C4-like | pub_C3-C4 | rob_C3 | 5.73846477 | 4.79E-09 | 0.95100352 | 1013313.5 | 16519.25 | 8949.25 | 2729.75 | 298 | 11316.5 | 2330 | 375 | 10089 | 90210.25 | 3157 | 444 | 1227 | 1562.5 | 108 |

| | | | | | | | | | | | | | | | | | | | | |
|---|---|---|---|---|---|---|---|---|---|---|---|---|---|---|---|---|---|---|---|---|
| ang_C3-C4 | pub_C3-C4 | son_C3-C4 | 4.47352711 | 3.85E-06 | 0.74821745 | 1076679.19 | 9836.8125 | 9377.3125 | 3158.3125 | 214.625 | 8759.625 | 2847.125 | 178.25 | 3771.875 | 104270.125 | 1572.75 | 187.25 | 1472.25 | 1376.25 | 58.25 |
| ang_C3-C4 | pub_C3-C4 | bro_C4-like | 8.45034642 | 0 | 0.11489865 | 1105586.5 | 7269.5 | 6843.25 | 6028.25 | 177.5 | 10013.5 | 1912.75 | 134 | 2447 | 108355.25 | 1095 | 130 | 980.5 | 1996 | 47 |
| ang_C3-C4 | pub_C3-C4 | tri_C4 | 4.21814567 | 1.23E-05 | 0.53060517 | 1101629.31 | 12329.1875 | 9277.6875 | 3566.4375 | 210.875 | 8705.0625 | 3228.0625 | 217.125 | 3610.5625 | 106884.063 | 1853.125 | 188.125 | 1499.125 | 1439.125 | 61.125 |
| koc_C4 | pub_C3-C4 | son_C3-C4 | 8.20338024 | 1.11E-16 | 0.67348033 | 1007307.81 | 9182.1875 | 8432.3125 | 3235.5625 | 202.875 | 11058 | 2664.5 | 209.5 | 3842.375 | 97027.375 | 1505 | 233.25 | 1317 | 1673.75 | 55.5 |
| koc_C4 | pub_C3-C4 | bro_C4-like | 12.1632073 | 0 | 0.1546735 | 1039640.75 | 6854 | 6234.25 | 5866.25 | 162 | 12470.25 | 1740 | 159 | 2495 | 101641 | 1071.5 | 170.5 | 892 | 2268.5 | 38 |
| koc_C4 | pub_C3-C4 | cro_C3 | 7.61172103 | 1.37E-14 | 0.93463702 | 1026035.13 | 11391.625 | 9136.5 | 2703.25 | 228.5 | 11668 | 2184.75 | 277 | 9598.875 | 91706.125 | 2495.25 | 459.5 | 1261.5 | 1510 | 88 |
| koc_C4 | pub_C3-C4 | chl_C3-C4 | 4.11461073 | 1.94E-05 | 0.02337143 | 1066156.5 | 4610.5 | 3899.5 | 8459.25 | 125.5 | 13606.75 | 895.75 | 94 | 1076.75 | 105237.5 | 620 | 98.75 | 532.75 | 2735.25 | 26.25 |
| koc_C4 | pub_C3-C4 | rob_C3 | 6.06907788 | 6.46E-10 | 0.94811578 | 1002629.44 | 16377.0625 | 8887.8125 | 2731.0625 | 295.375 | 11281.5 | 2309.5 | 386 | 10013 | 89235.5 | 3103.25 | 443 | 1235.25 | 1462 | 98.25 |
| son_C3-C4 | pub_C3-C4 | bid_C4 | 9.90906802 | 0 | 0.35127266 | 1113808.63 | 12542.1875 | 9267.125 | 4224.9375 | 248.375 | 10176.5625 | 2874.125 | 212.25 | 3605.5625 | 107736.625 | 1874.75 | 220.625 | 1419.25 | 1709.75 | 63.25 |
| son_C3-C4 | pub_C3-C4 | chl_C3-C4 | 6.12882822 | 4.44E-10 | 0.03047051 | 1123985.13 | 5063.4375 | 4162.625 | 9494.1875 | 125.375 | 12342.5625 | 865.375 | 69.25 | 1136.5625 | 110923.875 | 667.75 | 89.875 | 586 | 2698.5 | 26.5 |
| son_C3-C4 | pub_C3-C4 | ram_C3-C4 | 4.99212463 | 2.99E-07 | 0.23935763 | 1095367.25 | 9511.3125 | 9294.6875 | 3995.5 | 226 | 9927.6875 | 2876.5 | 207.5 | 3228.625 | 106070.688 | 1564.875 | 238.75 | 1444.875 | 1625.375 | 44.375 |
| son_C3-C4 | pub_C3-C4 | pal_C4-like | 8.72647907 | 0 | 0.31549215 | 1047230.94 | 11446.6875 | 8676 | 4074.625 | 250.25 | 9624.5625 | 2734.1875 | 217.875 | 3352 | 100956.375 | 1698.25 | 222 | 1344.75 | 1613.25 | 54.25 |
| son_C3-C4 | pub_C3-C4 | aus_C4 | 10.8763292 | 0 | 0.37664046 | 1084915.63 | 12230.6875 | 9027.0625 | 4087.125 | 245.25 | 9894.125 | 2754.1875 | 219.375 | 3559.5625 | 103460.875 | 1853.75 | 220.625 | 1348.25 | 1627.25 | 61.25 |
| son_C3-C4 | pub_C3-C4 | tri_C4 | 10.9920213 | 0 | 0.37387323 | 1090893.06 | 12552.3125 | 9055.8125 | 4191.6875 | 246.375 | 9945.125 | 2815.5 | 233 | 3637.25 | 105563.375 | 1881.25 | 219.5 | 1401.25 | 1681.25 | 65.25 |
| bro_C4_-ike | pub_C3-C4 | bid_C4 | 13.3808534 | 0 | 0.84186314 | 1125685.81 | 13826.9375 | 6651.6875 | 2690.6875 | 173.875 | 7460.3125 | 1831.3125 | 165.625 | 6406.3125 | 110371.813 | 2473.625 | 176.125 | 940.125 | 1168.625 | 47.125 |
| bro_C4_-ike | pub_C3-C4 | ano_C3-C4 | 26.2859875 | 0 | 0.56415175 | 1128420.75 | 7832 | 5867 | 3517.5 | 119 | 7807.75 | 1472.5 | 118 | 4119.5 | 112862 | 1500 | 125 | 794 | 1354 | 42 |
| bro_C4_-ike | pub_C3-C4 | ram_C3-C4 | 10.6371702 | 0 | 0.86240158 | 1095122.44 | 10638.8125 | 6557.0625 | 2509.0625 | 163.625 | 7230.25 | 1839.5 | 157 | 6036 | 107401.75 | 2088 | 173 | 943 | 1095.5 | 39 |
| bro_C4_-ike | pub_C3-C4 | pal_C4-like | 12.3483836 | 0 | 0.8431452 | 1074608.75 | 12814.5 | 6447.25 | 2548 | 178.5 | 7108 | 1774 | 175 | 5934.5 | 105216 | 2293 | 172 | 903 | 1119.5 | 35 |
| bro_C4_-ike | pub_C3-C4 | aus_C4 | 13.6032895 | 0 | 0.84002053 | 1098248.5 | 13512.5 | 6555 | 2601.5 | 170 | 7249 | 1744.25 | 179 | 6245.5 | 106492.25 | 2425.5 | 176.5 | 902.5 | 1120 | 42 |
| bro_C4_-ike | pub_C3-C4 | tri_C4 | 13.8174206 | 0 | 0.83796093 | 1093300.81 | 13760.9375 | 6512.9375 | 2638.4375 | 162.875 | 7301.3125 | 1762.3125 | 187.625 | 6293.0625 | 107245.063 | 2451.625 | 171.625 | 925.125 | 1155.125 | 42.125 |
| cro_C3 | pub_C3-C4 | bid_C4 | 8.27228539 | 0 | 0.06709513 | 1120497.75 | 13095.1875 | 9982.9375 | 10601.25 | 494.875 | 12562.625 | 2390.8125 | 314.125 | 2981.3125 | 100565.875 | 1703.125 | 256.625 | 1359.625 | 2786.75 | 90.125 |
| cro_C3 | pub_C3-C4 | chl_C3-C4 | 4.58975693 | 2.22E-06 | 0.01196722 | 1133488.56 | 5111.3125 | 4309.5625 | 16616.1875 | 226.625 | 14420.625 | 785 | 100 | 976.75 | 103464.125 | 576 | 100.75 | 546.25 | 3782 | 36.25 |
| cro_C3 | pub_C3-C4 | rob_C3 | 4.44017289 | 4.50E-06 | 0.72313878 | 1065895.56 | 15629.0625 | 14864.0625 | 4691.1875 | 528.125 | 9953.875 | 4316.75 | 409.5 | 5294.75 | 92957.375 | 2286.5 | 401 | 2124 | 1672.25 | 114 |
| cro_C3 | pub_C3-C4 | pal_C4-like | 6.58407824 | 2.30E-11 | 0.05573178 | 1062185.63 | 12028 | 9461 | 10092.375 | 474 | 11937.625 | 2315.5 | 292 | 2774.5 | 95122.625 | 1527.5 | 238.5 | 1305 | 2660.75 | 77 |
| cro_C3 | pub_C3-C4 | aus_C4 | 8.52064017 | 0 | 0.07155253 | 1092816.63 | 12774.75 | 9761.25 | 10170.625 | 486.5 | 12065.625 | 2346.25 | 302.5 | 2949.25 | 96883.375 | 1687 | 242.5 | 1301 | 2641.75 | 92 |
| cro_C3 | pub_C3-C4 | tri_C4 | 8.94768558 | 0 | 0.07416004 | 1091153.69 | 13032.4375 | 9706.6875 | 10357.8125 | 480.875 | 12190.1875 | 2380.3125 | 304.125 | 3019.3125 | 97930.1875 | 1699.625 | 256.125 | 1335.625 | 2713.875 | 92.125 |
| bid_C4 | pub_C3-C4 | chl_C3-C4 | 4.2213234 | 1.22E-05 | 0.0225337 | 1180691.75 | 5168.375 | 4321.25 | 9373.875 | 133.75 | 15350.1875 | 974.8125 | 99.125 | 1168.4375 | 117004.563 | 717.125 | 105.125 | 567.875 | 3057.375 | 28.375 |
| bid_C4 | pub_C3-C4 | rob_C3 | 5.22905303 | 8.54E-08 | 0.95679251 | 1110372.19 | 18114.6875 | 9878.25 | 2988.375 | 326.25 | 12736.25 | 2604.625 | 405.75 | 11102.4375 | 99297.8125 | 3453.625 | 479.375 | 1340.625 | 1660.625 | 119.125 |

| | | | | | | | | | | | | | | | | | | | | |
|---|---|---|---|---|---|---|---|---|---|---|---|---|---|---|---|---|---|---|---|---|
| flo_C3-C4 | pub_C3-C4 | aus_C4 | 4.39567401 | 5.53E-06 | 0.97798128 | 1159382.13 | 15191.8125 | 4094.625 | 1072.3125 | 90.125 | 4585.375 | 879.8125 | 98.125 | 9429.875 | 113820.563 | 3064.125 | 119.75 | 519.625 | 623.625 | 32.125 |
| chl_C3-C4 | pub_C3-C4 | ano_C3-C4 | 4.64454545 | 1.71E-06 | 0.96032524 | 1193493.19 | 8842 | 4353.3125 | 1214 | 60.75 | 5163.0625 | 999 | 75 | 6203.0625 | 121842.25 | 1913.25 | 107.125 | 581.25 | 738.5 | 22.25 |
| chl_C3-C4 | pub_C3-C4 | aus_C4 | 4.42193585 | 4.90E-06 | 0.97657897 | 1162855.44 | 15223.75 | 4292.5625 | 1134.25 | 106.25 | 5062.3125 | 935 | 106 | 9243.0625 | 114311 | 3051.25 | 142.125 | 547.25 | 686.5 | 29.25 |
| chl_C3-C4 | pub_C3-C4 | tri_C4 | 4.42099426 | 4.92E-06 | 0.97673888 | 1146901 | 15279.1875 | 4192.5 | 1114.1875 | 103.125 | 5014.375 | 916.8125 | 103.125 | 9204.625 | 113704.563 | 3039.375 | 135.25 | 554.875 | 686.625 | 27.375 |
| rob_C3 | pub_C3-C4 | aus_C4 | 5.96755914 | 1.21E-09 | 0.05032266 | 1082188.44 | 12487.25 | 9628.3125 | 10681.5 | 490.5 | 17505.0625 | 2523 | 383 | 2955.3125 | 95479.5 | 1632.5 | 330.125 | 1259 | 3305.5 | 116 |
| rob_C3 | pub_C3-C4 | tri_C4 | 6.20830313 | 2.69E-10 | 0.05158072 | 1088035.56 | 12787.9375 | 9666.9375 | 10944.4375 | 487.375 | 17697.5625 | 2578.3125 | 389.125 | 3033.3125 | 97307.3125 | 1670.625 | 340.125 | 1308.625 | 3372.625 | 117.125 |
| vag_C4-like | ram_C3-C4 | ang_C3-C4 | 5.19491173 | 1.03E-07 | 0.85604113 | 1020310.25 | 8452.25 | 7115 | 2751 | 156.5 | 9689.5 | 2401 | 191.5 | 4482.25 | 99120.25 | 1562.5 | 182 | 1138.5 | 1491.5 | 39 |
| vag_C4-like | ram_C3-C4 | son_C3-C4 | 4.60373126 | 2.08E-06 | 0.90729871 | 1021192.56 | 10014.9375 | 7166.1875 | 2633.4375 | 167.375 | 9763.5 | 2322.5 | 197 | 5365.75 | 98855.75 | 1838.5 | 211 | 1142 | 1442.5 | 52 |
| vag_C4-like | ram_C3-C4 | cro_C3 | 5.21742166 | 9.09E-08 | 0.96584771 | 1025279.94 | 11972.3125 | 7600.5 | 2233.75 | 213.5 | 10152.25 | 1903 | 251 | 11256.8125 | 91868.3125 | 2747.125 | 377.5 | 1045 | 1356 | 84 |
| ang_C3-C4 | ram_C3-C4 | koc_C4 | 5.55806808 | 1.37E-08 | 0.15619173 | 1009219.88 | 9639.125 | 7122.3125 | 4381.8125 | 183.125 | 8372.5625 | 2373.8125 | 199.625 | 2745.5 | 98011.75 | 1410.5 | 142.5 | 1157 | 1474.5 | 43 |
| ang_C3-C4 | ram_C3-C4 | bid_C4 | 5.74266835 | 4.67E-09 | 0.15232055 | 1112252.94 | 10797.8125 | 7775.8125 | 4837.5625 | 198.625 | 9269 | 2599 | 207 | 3001.25 | 108470 | 1618 | 164.5 | 1257 | 1671.5 | 41 |
| ang_C3-C4 | ram_C3-C4 | pal_C4-like | 4.21538991 | 1.25E-05 | 0.11835656 | 1049452.69 | 9889.5625 | 7326.3125 | 4607.5625 | 193.625 | 8729.625 | 2460.375 | 209.75 | 2748.625 | 102214.125 | 1444.25 | 140.25 | 1215.25 | 1584.75 | 40.25 |
| ang_C3-C4 | ram_C3-C4 | aus_C4 | 6.22382869 | 2.44E-10 | 0.17100538 | 1085740.06 | 10551.9375 | 7581.9375 | 4654.4375 | 194.375 | 8966.125 | 2562.625 | 193.25 | 2994.125 | 104545.375 | 1588.75 | 150.75 | 1221.25 | 1589.75 | 39.25 |
| ang_C3-C4 | ram_C3-C4 | tri_C4 | 6.24899283 | 2.07E-10 | 0.16700523 | 1087105.69 | 10815.3125 | 7569.0625 | 4774.8125 | 206.125 | 9098.375 | 2596.375 | 194.75 | 3033.125 | 106118.625 | 1611.75 | 154.75 | 1235.25 | 1642.75 | 35.25 |
| koc_C4 | ram_C3-C4 | son_C3-C4 | 4.59303016 | 2.19E-06 | 0.90400512 | 1009539.13 | 9853.125 | 7201.0625 | 2623.5625 | 144.625 | 9715.125 | 2314.375 | 200.75 | 5226.0625 | 97765.0625 | 1787.125 | 208.125 | 1146.125 | 1387.625 | 47.125 |
| koc_C4 | ram_C3-C4 | cro_C3 | 5.60484558 | 1.04E-08 | 0.96232276 | 1004645.56 | 11597.125 | 7493.5625 | 2226.25 | 201.5 | 10008.5625 | 1873 | 246 | 10895.4375 | 90116.125 | 2654.25 | 378.625 | 1031 | 1265 | 90 |
| koc_C4 | ram_C3-C4 | rob_C3 | 4.05927336 | 2.46E-05 | 0.97248747 | 1003908.94 | 16726.75 | 7460.8125 | 2320.25 | 256 | 9919.5625 | 2055 | 315.5 | 11430.8125 | 89739.75 | 3364.5 | 360.125 | 1033.5 | 1281.5 | 96 |
| son_C3-C4 | ram_C3-C4 | bid_C4 | 5.46943714 | 2.26E-08 | 0.10229192 | 1129208.44 | 11070.75 | 7970.5625 | 5919.75 | 224.5 | 11048.25 | 2531.6875 | 208.375 | 2917.75 | 109912.938 | 1616.875 | 183 | 1273.375 | 2015.375 | 52.375 |
| son_C3-C4 | ram_C3-C4 | pal_C4-like | 4.05390877 | 2.52E-05 | 0.08005122 | 1048214.31 | 9980.6875 | 7391.625 | 5568.375 | 223.75 | 10358.3125 | 2380.0625 | 202.625 | 2657.5 | 101759.25 | 1434 | 163 | 1192.5 | 1884 | 50 |
| son_C3-C4 | ram_C3-C4 | aus_C4 | 6.87167056 | 3.19E-12 | 0.12651152 | 1103852.44 | 10860.8125 | 7759.5 | 5726 | 227.5 | 10796.5 | 2434.75 | 215.5 | 2911.4375 | 105931.688 | 1610.375 | 174.875 | 1215.375 | 1931.875 | 46.375 |
| son_C3-C4 | ram_C3-C4 | tri_C4 | 6.84221526 | 3.92E-12 | 0.12352373 | 1112146.88 | 11142.9375 | 7780.5 | 5890.5625 | 235.125 | 10904.1875 | 2486 | 227.5 | 2965.8125 | 108372.375 | 1642.75 | 174.125 | 1246.75 | 2004.25 | 49.25 |
| bro_C4_-ike | ram_C3-C4 | tri_C4 | 4.33177385 | 7.40E-06 | 0.1021401 | 1076771.19 | 10716.0625 | 7587.5625 | 5268.5625 | 214.125 | 10507.375 | 2580.875 | 236.25 | 2886.625 | 104530.625 | 1568.25 | 176.25 | 1231.75 | 1885.25 | 47.25 |
| cro_C3 | ram_C3-C4 | bid_C4 | 7.12374628 | 5.29E-13 | 0.04409921 | 1111424.63 | 11317.3125 | 8225.8125 | 12222.625 | 411.625 | 12983.0625 | 2024 | 274 | 2494.5 | 100254.813 | 1485.5 | 219.5 | 1141 | 2984.625 | 91 |
| cro_C3 | ram_C3-C4 | rob_C3 | 4.35591672 | 6.63E-06 | 0.68116722 | 1067156.63 | 15573.25 | 14845.5 | 4815.125 | 505.75 | 10039.125 | 4445 | 393.75 | 5235.75 | 93209.875 | 2299 | 384.75 | 2156.5 | 1677.75 | 122.25 |
| cro_C3 | ram_C3-C4 | pal_C4-like | 4.48880604 | 3.58E-06 | 0.02924241 | 1042559.75 | 10289.625 | 7767.375 | 11492.5 | 398.25 | 12231.8125 | 1956.6875 | 251.875 | 2243.9375 | 93866.5625 | 1326.875 | 201.375 | 1096.875 | 2814.125 | 78.375 |
| cro_C3 | ram_C3-C4 | aus_C4 | 6.26076353 | 1.92E-10 | 0.04047309 | 1084897.19 | 11014.4375 | 8065.4375 | 11732.0625 | 434.375 | 12546.25 | 2011.875 | 261.75 | 2421.875 | 96631.25 | 1456.25 | 215.75 | 1088.25 | 2841 | 85.25 |
| cro_C3 | ram_C3-C4 | tri_C4 | 6.92531728 | 2.19E-12 | 0.04388335 | 1087904.56 | 11292.8125 | 8051.8125 | 12007.4375 | 421.625 | 12764.25 | 2034.125 | 269.75 | 2491.875 | 98207.25 | 1486.25 | 220.75 | 1119.75 | 2930.5 | 86.25 |
| ano_C3-C4 | ram_C3-C4 | aus_C4 | 4.59647494 | 2.15E-06 | 0.15631876 | 1156251.75 | 11091.6875 | 8061.25 | 4552.9375 | 188.875 | 9344.1875 | 2779.625 | 196.25 | 3108.1875 | 111755.875 | 1650.25 | 176.375 | 1344.25 | 1762.25 | 41.25 |

| | | | | | | | | | | | | | | | | | | | | |
|---|---|---|---|---|---|---|---|---|---|---|---|---|---|---|---|---|---|---|---|---|
| ano_C3-C4 | ram_C3-C4 | tri_C4 | 4.22019765 | 1.22E-05 | 0.14250263 | 1151950.63 | 11211.5625 | 7982.125 | 4662.0625 | 197.125 | 9448.4375 | 2826.375 | 199.75 | 3131.4375 | 112406.625 | 1642.75 | 178.375 | 1354.25 | 1812.25 | 40.25 |
| rob_C3 | ram_C3-C4 | aus_C4 | 4.59084583 | 2.21E-06 | 0.03000793 | 1099121.56 | 11039.9375 | 8129.9375 | 12417.9375 | 419.875 | 18178.875 | 2251.875 | 319.25 | 2566.375 | 97599.125 | 1472.75 | 279.25 | 1106.75 | 3619.25 | 97.25 |
| rob_C3 | ram_C3-C4 | tri_C4 | 4.67304438 | 1.49E-06 | 0.02995864 | 1105808.94 | 11303.5625 | 8127.5625 | 12737.5625 | 415.625 | 18452.875 | 2287.125 | 328.25 | 2609.875 | 99529.375 | 1499.75 | 287.25 | 1148.25 | 3709.75 | 97.25 |
| vag_C4-like | son_C3-C4 | cro_C3 | 9.39108458 | 0 | 0.90250399 | 1000691.94 | 10826.8125 | 9331 | 3058 | 255.5 | 12391.5 | 2390 | 288 | 8573.5625 | 89536.0625 | 2246.125 | 423 | 1234 | 1763.5 | 96 |
| vag_C4-like | son_C3-C4 | rob_C3 | 9.06559781 | 0 | 0.9074808 | 997973 | 15623.25 | 9164 | 3172.25 | 353 | 12191 | 2512.75 | 401 | 8981.5 | 89174.25 | 2908 | 414 | 1257.5 | 1711.5 | 115 |
| koc_C4 | son_C3-C4 | cro_C3 | 9.6876877 | 0 | 0.89809843 | 985896.25 | 10601.625 | 9164.125 | 3025.5 | 231.5 | 12204.625 | 2342.25 | 276 | 8364 | 88396.125 | 2202.25 | 414.75 | 1197.5 | 1654.5 | 97 |
| koc_C4 | son_C3-C4 | rob_C3 | 9.63217461 | 0 | 0.90336352 | 989744 | 15538.875 | 9000 | 3181.375 | 348.25 | 12081.375 | 2482.5 | 394.5 | 9015.625 | 88355.25 | 2897 | 427.25 | 1231.5 | 1620.5 | 115 |
| bro_C4 -ike | son_C3-C4 | cro_C3 | 4.59191622 | 2.20E-06 | 0.98132237 | 1041908.06 | 12622.125 | 5124.3125 | 1459.25 | 125.5 | 5656.8125 | 1223.25 | 149.5 | 13622.6875 | 95646.125 | 3166.75 | 220.625 | 656 | 827 | 36 |
| cro_C3 | son_C3-C4 | bid_C4 | 10.6764286 | 0 | 0.10508862 | 1086638.19 | 13812.75 | 10185.25 | 9261.8125 | 444 | 11785.5625 | 2549.25 | 317 | 3337.5 | 97917.5625 | 1909.5 | 265.5 | 1338.5 | 2438.625 | 99 |
| cro_C3 | son_C3-C4 | flo_C3-C4 | 4.503906 | 3.34E-06 | 0.04543926 | 1091144.13 | 11258 | 10169.5 | 9227.625 | 420.25 | 11850.625 | 2484.25 | 254 | 2805.25 | 98114.375 | 1549.75 | 258 | 1298.75 | 2476.25 | 93.25 |
| cro_C3 | son_C3-C4 | chl_C3-C4 | 5.57209929 | 1.26E-08 | 0.05539475 | 1093332.63 | 11653.5 | 10240 | 9308.875 | 420 | 11996.875 | 2496.5 | 256.5 | 2896 | 98404.875 | 1568.5 | 261.5 | 1300.5 | 2479.75 | 85 |
| cro_C3 | son_C3-C4 | rob_C3 | 8.63551758 | 0 | 0.6036915 | 1053476 | 15256.625 | 14016.375 | 4837 | 502.75 | 9876.125 | 4085.5 | 376.25 | 5230.25 | 92778.625 | 2325.75 | 363.25 | 1953.25 | 1623.75 | 117.5 |
| cro_C3 | son_C3-C4 | ram_C3-C4 | 4.2420945 | 1.11E-05 | 0.04489893 | 1072368.38 | 10702.25 | 10099.75 | 9074.875 | 393.5 | 11679.125 | 2560.25 | 249.5 | 2866.5 | 96939.625 | 1580.5 | 266.5 | 1293 | 2412.25 | 92 |
| cro_C3 | son_C3-C4 | pal_C4-like | 8.81583277 | 0 | 0.09304539 | 1019420.75 | 12658.875 | 9630.125 | 8670.25 | 427.75 | 11055.625 | 2475.75 | 291 | 3111.25 | 91564.875 | 1731 | 254.5 | 1265.5 | 2302.75 | 82 |
| cro_C3 | son_C3-C4 | aus_C4 | 10.6168126 | 0 | 0.10890144 | 1060489.31 | 13398.3125 | 9935.0625 | 8873.1875 | 464.625 | 11342.625 | 2513.25 | 295.5 | 3290.5 | 94333.875 | 1855.5 | 246.5 | 1287 | 2334.75 | 95 |
| cro_C3 | son_C3-C4 | tri_C4 | 10.872083 | 0 | 0.11023542 | 1066255.25 | 13754.375 | 9969.625 | 9106.5 | 482.75 | 11544.125 | 2586.75 | 307 | 3394.5 | 96109.375 | 1910.5 | 258.5 | 1315.5 | 2392.25 | 101 |
| bid_C4 | son_C3-C4 | rob_C3 | 9.45191077 | 0 | 0.90883408 | 1103629.31 | 17373.9375 | 10112.875 | 3511.375 | 385.25 | 13764.0625 | 2787.3125 | 431.125 | 10005.5 | 99082.25 | 3209.5 | 449.5 | 1401.5 | 1870.5 | 128 |
| flo_C3-C4 | son_C3-C4 | rob_C3 | 4.71869245 | 1.19E-06 | 0.95512852 | 1110402.63 | 17636.375 | 10090.875 | 2997.875 | 392.5 | 11268 | 2650.25 | 338 | 10049.75 | 99338.5 | 3246.75 | 420 | 1336.75 | 1571.5 | 116.25 |
| chl_C3-C4 | son_C3-C4 | rob_C3 | 5.12006851 | 1.53E-07 | 0.95078941 | 1110850.38 | 17706.375 | 10199.125 | 3064.125 | 388.25 | 11689.25 | 2684.25 | 333 | 10023.75 | 99663.5 | 3252.5 | 428 | 1355.5 | 1584 | 95 |
| rob_C3 | son_C3-C4 | ram_C3-C4 | 5.14245578 | 1.36E-07 | 0.05073515 | 1095607.5 | 10788.875 | 10058 | 9853.875 | 398.25 | 17430.1875 | 2699.5625 | 329.125 | 3081.9375 | 98786.8125 | 1575.625 | 380.375 | 1361.125 | 3198.625 | 109.125 |
| rob_C3 | son_C3-C4 | pal_C4-like | 7.86107451 | 2.00E-15 | 0.08114967 | 1023283.81 | 12486.625 | 9460.6875 | 9258.875 | 415.75 | 16106.5625 | 2651.25 | 385.5 | 3234.8125 | 91634.5 | 1723 | 362.625 | 1288 | 2981 | 112 |
| rob_C3 | son_C3-C4 | aus_C4 | 9.55408986 | 0 | 0.09514441 | 1080827.56 | 13440.3125 | 9903.1875 | 9637.5625 | 482.625 | 16867.375 | 2739 | 406 | 3464.375 | 95768.75 | 1844.5 | 376.25 | 1328 | 3094.5 | 119 |
| rob_C3 | son_C3-C4 | tri_C4 | 9.3782509 | 0 | 0.09247264 | 1090323.56 | 13824.375 | 9964.9375 | 9906.375 | 503.75 | 17141.3125 | 2822.5 | 415 | 3544.3125 | 98046.25 | 1892.5 | 391.125 | 1367 | 3154 | 130 |
| koc_C4 | vag_C4-like | bid_C4 | 11.0363814 | 0 | 0.70980569 | 1030906.69 | 4590.375 | 3231.25 | 1299.8125 | 45.125 | 3672.25 | 816.3125 | 32.625 | 1998.9375 | 112365.25 | 855.5 | 34.875 | 515.5 | 568 | 13.5 |
| koc_C4 | vag_C4-like | aus_C4 | 9.56525707 | 0 | 0.80378986 | 999709.438 | 5025.875 | 3154.5625 | 1155.875 | 38.75 | 3505.8125 | 757.875 | 43.25 | 2388.3125 | 107305.125 | 950.25 | 32.625 | 491.75 | 536.25 | 9.25 |
| koc_C4 | vag_C4-like | tri_C4 | 9.85086591 | 0 | 0.79187123 | 998778.313 | 5138.3125 | 3180.8125 | 1210.6875 | 39.875 | 3546.6875 | 792.0625 | 44.625 | 2384.8125 | 108406.688 | 958.375 | 35.625 | 494.875 | 547.875 | 10.375 |
